# Microengineered transplantation of human solid tumors for in vitro studies of CAR T immunotherapy

**DOI:** 10.1101/2025.04.25.650709

**Authors:** Haijiao Liu, Estela Noguera-Ortega, Xuanqi Dong, Won Dong Lee, Jeehan Chang, Sezin Aday Aydin, Yumei Li, Yonghee Shin, Xinyi Shi, Maria Liousia, Marina C. Martinez, Joshua J. Brotman, Soyeon Kim, Zeyu Chen, Anni Wang, Zirui Ou, Jungwook Paek, Ju Young Park, Aidi Liu, Haonan Hu, Zebin Xiao, Dora Maria Racca, Se-jeong Kim, G. Scott Worthen, Wei Guo, Ellen Puré, Taewook Kang, Joshua D. Rabinowitz, E. John Wherry, Edmund K. Moon, Steven M. Albelda, Dan Dongeun Huh

## Abstract

Treatment of solid malignancies using chimeric antigen receptor (CAR) T cells remains a significant challenge, but current efforts to advance this therapy are challenged by our limited capacity to probe and understand cancer-immune interactions in human solid tumors. Here, we present a microengineered platform for in vitro modeling of malignant solid tumors during CAR T therapy. This system makes it possible to vascularize human tumor explants and perfuse them with blood-borne immune cells in a controlled manner. We first present a microphysiological model of human lung adenocarcinomas infused with CAR-T cells and show how this system can be used to simulate, visualize, and interrogate tumor-directed trafficking and effector function of CAR T cells. We then demonstrate the proof-of-principle of testing a chemokine-directed CAR T cell engineering strategy in a model of malignant pleural mesothelioma and validating our in vitro assessment using a matching in vivo mouse model. Finally, we describe a potential therapeutic target discovered by single-cell RNA sequencing that can be pharmacologically modulated to increase the efficacy of CAR T cells for lung adenocarcinoma, for which we also present specific biomarkers identified by global metabolomics analysis. We believe that the bioengineering principle demonstrated here will make important contributions to developing new capabilities for preclinical studies of adoptive cell therapies for cancer and other complex diseases.

---

Among the key strategies of cancer immunotherapy is to use antigen-directed cytotoxicity of T lymphocytes to target and destroy cancer cells, which is known as adoptive T cell therapy^7,8^. As one of the main modalities of this approach, chimeric antigen receptor (CAR) T cell therapy uses autologous or allogenic T cells genetically engineered to express synthetic CARs that can bind to target molecules on the surface of cancer cells to induce antigen-specific cytotoxicity^9–11^. By using CD19-targeted CAR T cells, for example, this method demonstrated highly promising anti-cancer effects in patients with malignant B lymphocytes^12–14^, leading to the recent approval of CAR T therapies for treating several types of leukemia and advanced B-cell lymphoma^10,11,15–17^.

In contrast to the clinical success of CAR T therapy for hematologic malignancies, limited progress has been made in translating this technology into an effective treatment for solid tumors, which account for more than 90% of all cancer fatalities^18–21^. Unlike blood cancer cells that express common tumor-specific antigens, malignant cells in most solid tumors lack unique surface markers and are often enriched with antigens that are highly heterogenous and also found on normal cells at lower levels^18,22^. This makes it difficult to identify disease-specific target antigens required for the efficacy and safety of CAR T therapy. Another challenge arises from the local environment surrounding solid tumors, termed the tumor microenvironment (TME), that forms a physical barrier to tumor-directed CAR T cell trafficking and produces immunosuppressive signals to attenuate or inhibit anti-tumor function of CAR T cells^18,23–26^. Understanding and modulating these key aspects of cancer-immune interactions are the major focus of current efforts to develop more effective and safer CAR T therapies for solid malignancies^18,27–29^.

What is generally required for these types of studies is to use live tumor-bearing animals, most commonly mice, to model the complexity of TME in human solid tumors and examine the interaction of circulating CAR T cells with malignant cells in vivo^30–33^. As a common method, for example, xenograft models established by transplanting human tumor cell lines or patient-derived tumors into immunodeficient or humanized mice allow sustained engraftment of human cancer cells, providing a platform to assess anti-tumor activity of infused human CAR T cells in vivo^34–36^. Researchers have also developed syngeneic and transgenic models in which the behavior of murine CAR T cells in mouse tumors can be studied in vivo in the context of a fully competent immune system^27,37,38^, which has proven particularly instrumental for hypothesis-driven mechanistic investigation of cancer-immune interactions^39–41^.

These animal-based methods continue to evolve and serve as the mainstay of preclinical research in CAR T therapy, but they also face unique challenges due to the nature of CAR T cells as “living drugs”. Unlike traditional pharmacological agents, the therapeutic efficacy of administered CAR T cells relies on their ability to engage in highly coordinated activities, such as migration, activation, proliferation, and persistence, all of which are dynamically regulated by their interactions with various tumor-derived biological signals^42^. This complex nature of therapy poses challenges to the design of clinically relevant animal models and in-depth analysis of biological processes underlying therapeutic effects and toxicities to produce pharmacology and toxicology data^42–44^. Animal studies of CAR T cells have also come under increased scrutiny recently as concerns regarding their limited predictive capacity grow, especially due to the failure of recent clinical trials caused by severe immune-mediated toxicities of CAR T cells preclinically validated for the treatment of solid malignancies^45,46^. Given these ongoing challenges, a consensus is emerging that efforts should be made to develop new approaches that can complement animal studies by providing a means to generate preclinical data that are more relevant and translatable to human situations^47–50^.

Motivated by this emerging trend, here we introduce a bioengineering approach that combines the advanced capabilities of microengineered cell culture with the inherent complexity of solid tumors in vivo to model cancer-immune interactions in vitro. Our technology makes it possible to transplant living tumors from xenograft models or human patients into engineered vascular beds to generate vascularized human malignant solid tumor constructs that can be maintained for prolonged periods and perfused with human CAR T cells in a controlled manner. Using human lung adenocarcinoma and mesothelioma tumors as model systems, we demonstrate i) in vitro modeling and direct visualization of the entire process of CAR T trafficking and anti-tumor activities and ii) phenotypic and transcriptomic interrogation of CAR T cells in tumor tissues using flow cytometry and single-cell RNA sequencing. We also show how this technology may be leveraged for the development of new therapeutic approaches by presenting a druggable target discovered by ligand-receptor analysis of our lung adenocarcinoma model that can be pharmacologically modulated to increase tumor-directed CAR T cell trafficking. Finally, we demonstrate the feasibility of using our model to discover metabolic biomarkers of therapeutic efficacy that may be useful for continuous monitoring of this treatment strategy.

## Results

### Construction of microengineered platform for in vitro transplantation of solid tumors

Tumor transplantation is an established technique used for the study of tumorigenesis and metastasis in which malignant tumor tissues are surgically removed from donors and implanted into anatomically appropriate sites in recipient animals^31,33,51^ (**Fig. 1a**). Essential for the success of this method is engraftment of tumor transplants, which is achieved by their anastomosis with the host vasculature and subsequent vascular perfusion. Our approach is based on the same principle with the exception that tumor tissues are transplanted into microengineered recipient devices with externally accessible living stroma that contains a three-dimensional (3D) network of perfusable human blood vessels (**Fig. 1b**).

**Fig. 1.**
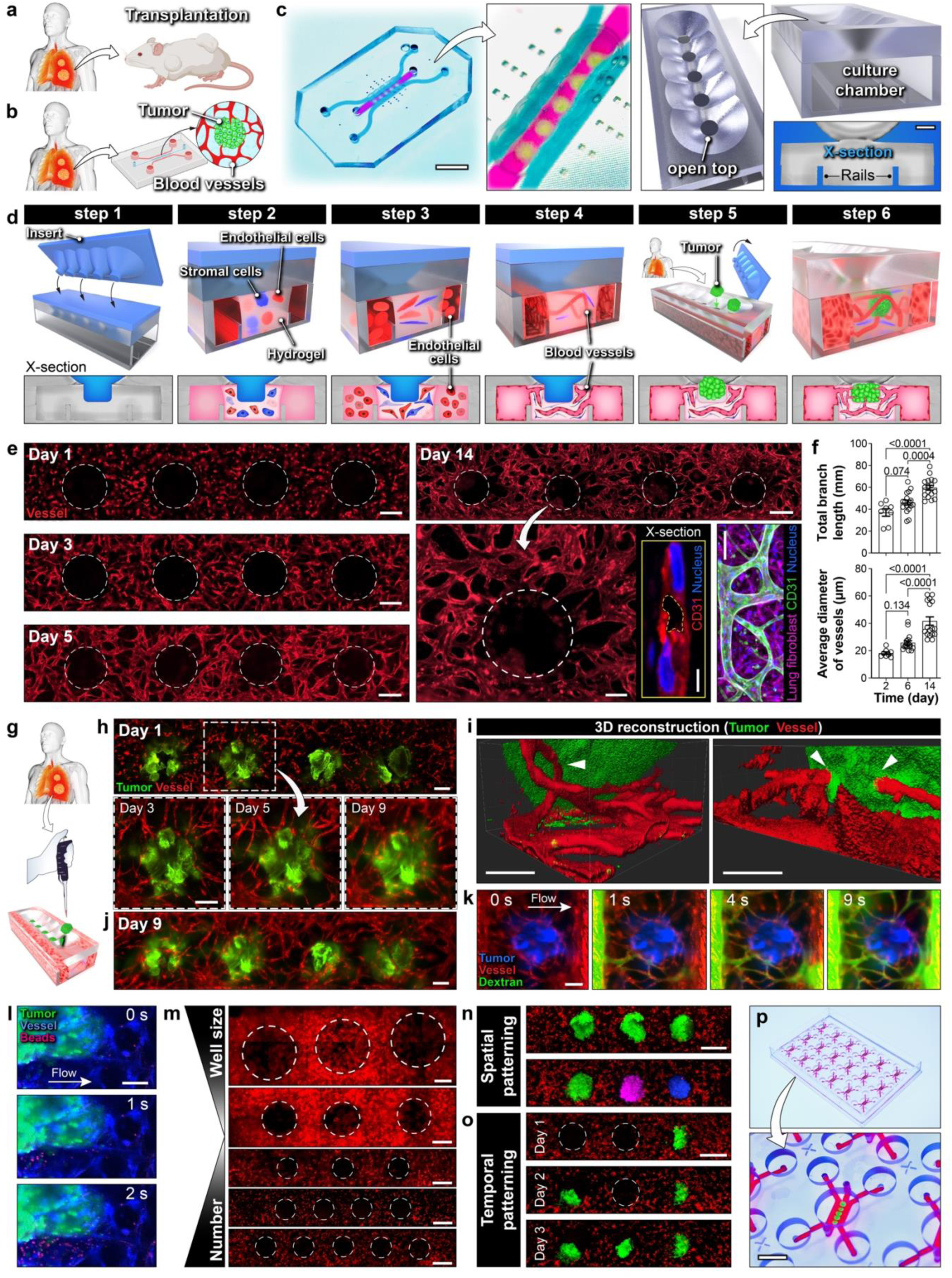
Microengineered platform for in vitro transplantation and prolonged maintenance of human solid tumors. **a,b,** Conceptual illustration of solid tumor transplantation in vivo (**a**) and in our microengineered device (**b**) designed to reconstitute the tumor-vascular interface. Created with Biorender.com. **c,** Photos and illustration of the open-top device with a compartmentalized chamber design. Scale bars, 5 mm (left) and 250 µm (bottom-right). **d,** Illustration of experimental procedure to construct vascularized and perfusable tumor constructs in the device. **e,** Representative fluorescence micrographs showing blood vessel development in the device over time. The dashed white circles show the location of closed culture wells. Scale bars, 250 µm (multi-well view), 100 μm (single-well view and image of human lung fibroblasts stained magenta), and 5 µm (inset showing the cross-section of a blood vessel). **f,** Quantification of total branch length (per engineered construct) (top) and the average diameters of microvessels (bottom) during the vascularization process in our microengineered platform. Data are presented as mean ± SEM (n = 8-17). **g-j,** Time-lapse fluorescence micrographs showing progressive vascularization of primary lung cancer explants. Scale bars, 200 μm. **i,** 3D reconstruction of a small part of a vascularized tumor showing the engineered microvessels wrapping around the tumor to form close contact with the tumor surface (left, arrowhead), and their ability to penetrate the tumor construct (right, arrowheads). Scale bars, 100 µm. **k,l,** Time-lapse fluorescence micrographs showing directional vascular perfusion of tumor constructs cultured for 10 days using 70-kDa FITC Dextran (**k**) and 1-µm fluorescent microbeads (**l**). Scale bars, 200 μm (**k**) and 100 µm (**l**). **m,** Examples of different array designs with different numbers or sizes of culture wells (shown with dashed white circles). Red in the images shows fluorescence of blood vessels embedded in the culture scaffold. Scale bars, 500 µm. **n,o,** Demonstration of controlling the spatial distribution (**n**) or timing (**o**) of in vitro tumor transplantation into the culture-well array. Tumors shown in **n** were color-coded to represent different types/compositions. Scale bars, 500 µm. **p,** Photos of a device containing an array of 18 individually accessible and controllable transplantation units to scale up the production of vascularized solid tumor constructs. Scale bar, 5 mm.

To enable this in vitro transplantation, we created a 3D culture system in a microdevice that consists of individually addressable three parallel cell culture chambers defined by two microfabricated rails protruding from the bottom surface (**Fig. 1c**). Another key design feature is a longitudinal array of funnel-shaped holes in the ceiling of the device, which provide direct access to the culture chamber s from the external environment (**Fig. 1c, Supplementary Fig. 1**). This open-top design allows for i) the production of microengineered 3D tissue reminiscent of the vascularized stroma surrounding solid tumors and ii) on-demand transplantation of tumor explants into the vascularized stromal constructs.

The first step of this sequential process is to block the open-top access in the device ceiling using a micropatterned insert designed to fit into the funnel-shaped wells (**Step 1** in **Fig. 1d**). Next, an extracellular matrix (ECM) hydrogel precursor solution containing suspended human vascular endothelial and stromal cells is injected into the middle chamber using an independent fluidic access port and solidified to embed and grow the cells in 3D with culture media in the side chambers (**Step 2** in **Fig. 1d, Supplementary Fig. 2**). After 24 hours, the side chambers are seeded with endothelial cells (**Step 3** in **Fig. 1d**). Designed to mimic de novo blood vessel formation during vasculogenesis^52^, this culture configuration induces the formation of tubular structures by the endothelial cells in the hydrogel and their assembly into an interconnected 3D vascular network that anastomoses with the endothelial lining of the side chambers (**Step 4** in **Fig. 1d**). The microengineered vascular bed can be maintained in the device until tumor explants become available for in vitro transplantation. This procedure begins with the removal of the insert from the device ceiling, which is followed by injection of tumor tissues cut into an appropriate size into the funnel-shaped wells and then by injection of acellular ECM solution to cover and seal the wells (**Step 5** in **Fig. 1d**). Over the next few days after gelation, the tumor implants embedded in the microengineered stroma become vascularized by the surrounding blood vessels, which also makes the tumors accessible and perfusable from the side chambers (**Step 6** in **Fig. 1d**).

For proof-of-concept demonstration, we first generated a 3D stromal tissue by culturing primary human lung fibroblasts and primary human umbilical vein endothelial cells (HUVECs) in a fibrin hydrogel in the middle chamber (**Fig. 1e**). In this environment, the HUVECs became elongated to form thin sprouts initially but over a period of 5 days, these structures adjoined together to create a network of connected endothelial tubes throughout the scaffold (**Fig. 1e**). The self-assembled microvasculature continued to develop over longer periods of time (over 14 days) without a loss of structural integrity and vascular connectivity (**Figs. 1e, 1f**). This microengineered construct was then used for in vitro transplantation of lung tumors surgically removed from lung cancer patients (**Fig. 1g**). Time-lapse analysis showed directional growth of existing blood vessels in the surrounding area towards the tumor deposited into the open well within 3-5 days of transplantation and as a result, the entire implant was enveloped by the vasculature by day 9 (**Fig. 1h**). During this process, the initial size of transplanted tumors appeared to influence some of the characteristics of tumor vascularization, such as the vessel-covered area and the number of vessel junctions (**Supplementary Fig. 3**). Closer examination of the tumor-vascular interface revealed that some of the vessels can also penetrate the tumors (**Fig. 1i, Supplementary Movies 1, 2**). In any given device, vascularized tumors were observed across the entire array of open wells (**Fig. 1j**). Importantly, when a pressure difference was generated across the hydrogel, it was possible to flow media between the side chambers through the microengineered blood vessels, making it possible to perfuse the vascularized tumors and their local microenvironment in a controlled manner (**Figs. 1k, 1l, Supplementary Movies 3, 4**).

The multi-well design of this platform allows for implantation and simultaneous vascularization of multiple tumors within the same construct. This is useful for modeling microscopic clusters of tumor cells surrounded by vascularized stroma, known as tumor nests, which have been recognized as an important microarchitectural feature of various types of malignant solid tumors in vivo (**Supplementary Fig. 4**)^53,54^. Since the size and relative position of the wells are readily adjustable during device fabrication, this design also provides a means to pattern the spatial arrangement of transplanted tumors with a high degree of controllability (**Fig. 1m**). Individual accessibility of the open wells adds to this capability by permitting spatiotemporal variation of cell/tissue types deposited into the wells for studies that require, for example, the inclusion of different types of solid tumors or a combination of malignant and normal tissues within the same vascularized construct (e.g., in vitro modeling of cancer metastasis) (**Figs. 1n, 1o**). Finally, our platform can be further engineered to form and maintain vascularized tumor constructs in an array format for increased experimental throughput (**Fig. 1p**) and may also be used for in vitro vascularization of tumor spheroids or organoids (**Supplementary Fig. 5, Supplementary Movie 5**).

### In vitro modeling, direct visualization, and analysis of CAR T cell-tumor interactions

Next, we set out to show the capabilities of our platform for the study of how CAR T cells interact with microengineered tumor transplants. The focus of this work was to demonstrate the use of our system as an in vitro platform to model and probe three essential steps of CAR T cell trafficking, including i) their extravasation and infiltration into solid tumors, ii) recognition of tumor-associated antigens by infused CAR T cells, and iii) their survival and persistent anti-tumor function (**Fig. 2a**), which also represent three critical challenges of CAR T therapies for solid tumors.

**Fig. 2.**
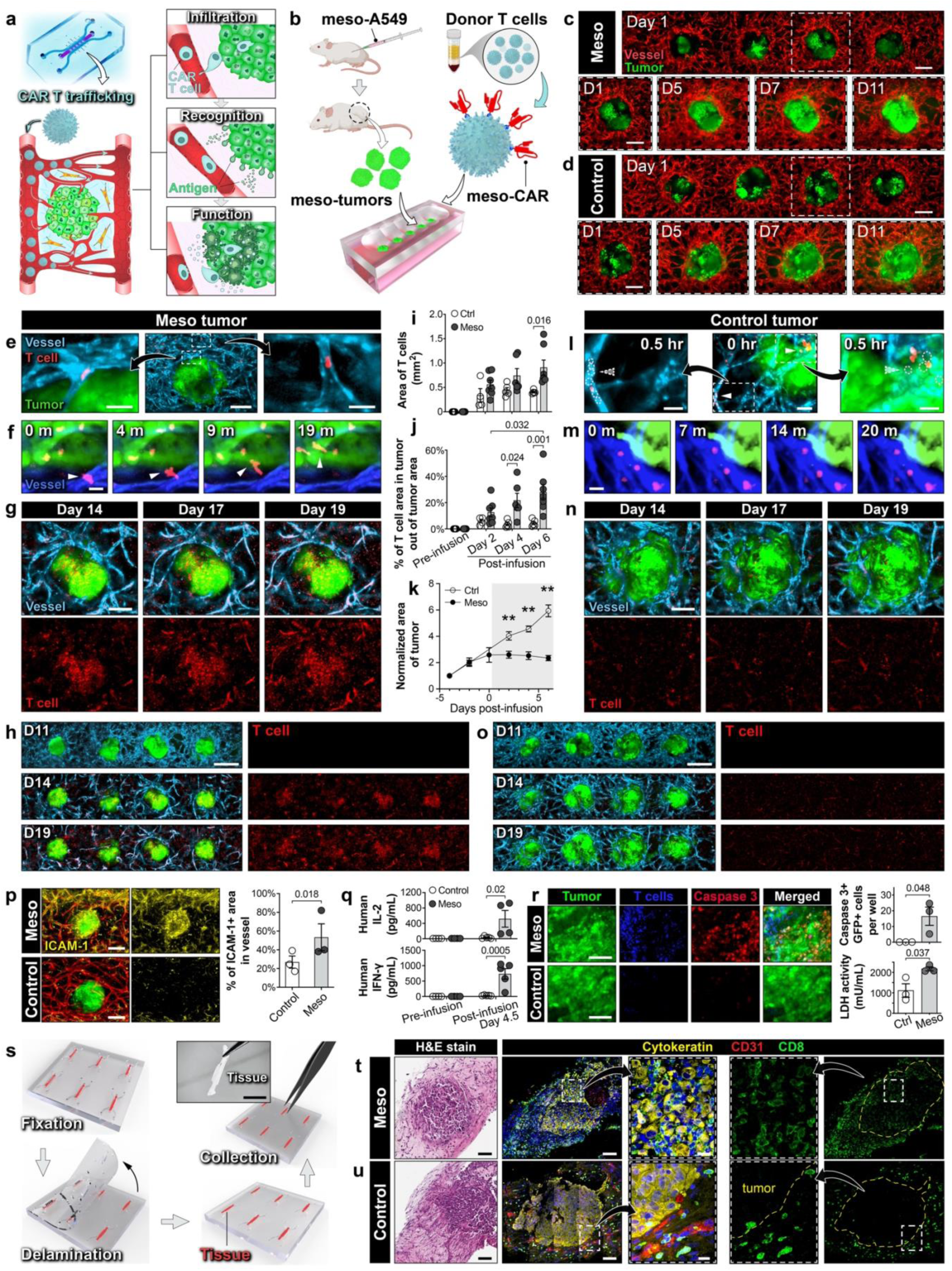
In vitro modeling of CAR T cell-tumor interactions. **a,** Illustration of three essential steps of tumor-directed CAR T cell trafficking modeled in the tumor-on-a-chip. **b,** Experimental workflow to model the interaction of mesothelin-targeted CAR T cells with mesothelin-overexpressing xenograft lung tumors (meso-tumors). Created with Biorender.com. **c,d**, Representative confocal images demonstrating rapid vascularization and growth of control (**c**) and meso-tumors (**d**) over 11 days. The lower panels in each figure show a close-up of tumor marked with a dashed square in the upper image. Scale bars, 250 µm. **e,** Micrographs of meso-CAR T cells adherent to blood vessels associated with meso-tumors on day 0 of CAR T infusion. Scale bars, 50 µm (left, right) and 250 µm (middle). **f,** Time-lapse images of CAR T cell extravasation and migration towards tumors. The images were taken 24 hours after CAR T cell infusion. Scale bars, 20 µm. **g,h,** Confocal images showing the accumulation of CAR T cells in meso-tumors over time. Scale bars, 200 µm (**g**) and 500 µm (**h**). **i,j,k,** Quantification of tissue area occupied by CAR T cells (**i**), the percentage of CAR T cells found within tumors (**j**), and tumor area normalized to that prior to CAR T cell infusion (**k**). Data are presented as mean ± SEM. \*\**P* < 0.01 (n ≥ 4). **l-o**, Micrographs of meso-CAR T cells infused into control tumors at various time points. Scale bars, 50 µm (left, right) and 100 µm (middle) (**l**), 20 µm (**m**), 200 µm (**n**), and 500 µm (**o**). **p,** Immunostaining of ICAM-1 expression (left) and quantification of tissue area stained positive for ICAM-1 (right). Scale bars, 250 µm. **q,** Measurement of secreted human IL-2 and IFN-ɣ. Data are presented as mean ± SEM (n ≥ 4). **r,** Visualization of apoptotic tumor cells at Day 5 post CAR T cell infusion (left) and quantification of caspase 3 expression and LDH release (right). Data are presented as mean ± SEM (n ≥ 3). Scale bars, 100 μm. **s,** Illustration of experimental procedure to harvest whole tumor constructs from culture chambers. Scale bar, 5 mm. **t,u,** Histological sections of CAR T cell-infused meso-(**t**) and control tumors (**u**). Scale bars, 100 μm and 20 µm (zoom-in view). The yellow dashed polygons in the rightmost images show the outline of tumors. CAR T cells derived from two healthy donors were tested in this study.

To demonstrate the proof-of-principle of this approach, we used a complementary pair of malignant human lung tumors and CAR T cells (**Fig. 2b**) – lung tumors formed in a cell line-derived xenograft (CDX) model using A549 human lung adenocarcinoma cells engineered to overexpress mesothelin (**Supplementary Fig. 6**), a tumor-associated antigen expressed at significantly higher levels in various types of cancer^55–57^, and healthy donor-derived T cells transduced to express human mesothelin-targeted CARs (referred to as meso-CAR T cells hereafter) (**Supplementary Fig. 6**), which are being tested in ongoing clinical trials for immunotherapy of solid tumors^58–61^. In this study, the meso-CAR T cells were derived from three healthy donors. When the mesothelin-overexpressing tumors (referred to as meso-tumors hereafter) were transplanted into a microengineered vascular bed, they were vascularized within 7 days and exhibited rapid growth over time, as illustrated by their enlargement over the course of 11 days (**Fig. 2c, Supplementary Fig. 7**). Similar patterns of tumor vascularization and growth were observed in the control group containing human lung tumor explants from a CDX model established using wild-type A549 cells that express very low levels of mesothelin (referred to as control tumors hereafter) (**Fig. 2d, Supplementary Figs. 6, 7**).

By day 13, a significant tumor mass was achieved in both meso-tumor and control groups, at which time a single dose of meso-CAR T cells was administered into the models. The CAR T cells infused into the side chamber of the meso-tumor-containing devices immediately flowed into the 3D vasculature in the hydrogel but during perfusion, many of these cells adhered to the blood vessels surrounding the tumors and began to spread and migrate along the vascular lumen, displaying the phenotype of activated T cells (**Fig. 2e, Supplementary Movie 6**). Some of these adherent cells formed aggregates initially but over the next 24 hours, they were observed to move around as individual cells, undergo diapedesis across the endothelium, and extravasate into the surrounding stroma, which was followed by directional migration towards the tumors (**Fig. 2f**, **Supplementary Movie 7**). On the next day, the tumors were seen with a large number of CAR T cell infiltrates, and their number continued to increase over time during 6-day culture post infusion (**Figs. 2g, 2i, 2j**). Notably, the projected area of these tumors remained relatively constant during this period, indicating arrested tumor growth (**Figs. 2h, 2k, Supplementary Fig. 7**).

These results were in contrast to what was observed in the control group. After a single dose of meso-CAR T cell infusion under identical conditions, the vast majority of the administered cells flowed through the tumor-associated vasculature without establishing firm attachment (**Fig. 2l, Supplementary Movie 6**). A small number of T cells were detected within the vascular network 24 hours post infusion but most of them showed round morphology and remained at their original locations without any noticeable motility (**Fig. 2m, Supplementary Movie 8**). In this group, the number of CAR T cells in the tumors did not change over time (**Figs, 2i, 2j, 2n**), and their effects on tumor growth were negligible as shown by a 6-fold increase in the tumor area by the end of 6-day culture following cell infusion, indicating a lack of cytotoxicity of meso-CAR T cells against control tumors (**Figs. 2k, 2o**). Of note, when the tumor constructs were infused with non-transduced human T cells, their tumor-directed trafficking and inhibitory effects on tumor growth were negligible in both the control and meso-tumor models (**Supplementary Fig. 8**).

To gain more insight into this significant difference, we first measured endothelial expression of intercellular adhesion molecule-1 (ICAM-1) prior to CAR T cell infusion. Compared to those associated with control tumors, the blood vessels present in the meso-tumors were seen with significantly more robust expression of ICAM-1 (**Fig. 2p**), supporting the observation of increased CAR T cell adhesion to the vasculature. Interestingly, analysis of 38 common chemokines using ELISA of device effluent showed no significant differences between the control and meso-tumor models, except for RANTES (CCL5), MCP-1 (CCL2), Ckβ 8-1 (CCL23), and CXCL16, all of which were detected in higher concentrations in the meso-tumor model (**Supplementary Fig. 9**). This result may indicate that these chemokines can contribute to increased CAR T cell recruitment in the meso-tumor model, but our data showing little to no trafficking of non-transduced T cells in the same group (**Supplementary Fig. 8**) suggest that CAR-mediated recognition of higher mesothelin expression by T cells may be a more important factor for understanding the enhanced activity of CAR T cells in this model.

After CAR T cell treatment, ELISA of device effluent showed significant production of interleukin-2 (IL-2) and interferon-gamma (IFN-γ), which are well-known markers of T cell activation and cytotoxic activity^62,63^, in the meso-tumor model, whereas the control group did not yield any measurable amounts (**Fig. 2q**). This result was corroborated by significant cell death in the meso-tumor group evidenced by substantially increased caspase-3 expression by tumor cells and LDH release into the vascular perfusate (**Fig. 2r**).

For further analysis, we developed a technique to peel apart the layers of the sealed culture device and harvest the vascularized tumor tissues intact after CAR T cell infusion and incubation (**Fig. 2s, Supplementary Fig. 10**). Importantly, this method permitted immunohistological examination of the entire tumor constructs, which provided direct evidence of extensive infiltration and accumulation of CD8+ T cells in the meso-tumors (**Fig. 2t**). The tumor-killing effects of these CAR T cell infiltrates were also evident from the fragmented morphology of tumor cells (inset, **Fig. 2t**). In contrast, accumulation of CD8+ cells in the control group was only observed along the tumor boundaries with very few cells visible in the tumor mass (**Fig. 2u**), approximating the spatial characteristics of T cell exclusion reported in solid tumors in vivo^54,64^.

Our models described here clearly show significant differences in immune responses of CAR T cells caused by their differential interactions with lung tumors expressing significantly different levels of mesothelin. These data verify the principle of the mesothelin-targeted approach simulated in this proof-of-concept study but they also demonstrate the feasibility of using our microengineered tumor transplants to emulate and visualize CAR T cell trafficking and persistence in solid tumors in a more physiologically relevant experimental setting.

### Spatially-defined analysis of CAR T cell phenotypes

Given that the ability of T cells to localize to tumor sites and carry out their anti-tumor function is reflected in their phenotypes^65^, we next investigated whether and how meso-CAR T cells infused into our models change their phenotype in the microengineered tumor constructs. Our analysis was conducted in a spatially defined manner to distinguish T cells within the tumors from those in the surrounding vascularized stroma to interrogate their characteristics separately. This was achieved by isolating only the tumors from the construct through the openings of the wells 6 days post infusion and dissociating them to retrieve CAR T cell infiltrates (**Fig. 3a**). Subsequently, the remaining tissues without the tumors were harvested from the devices and processed to release CAR T cells in the stroma (**Fig. 3a**).

**Fig. 3.**
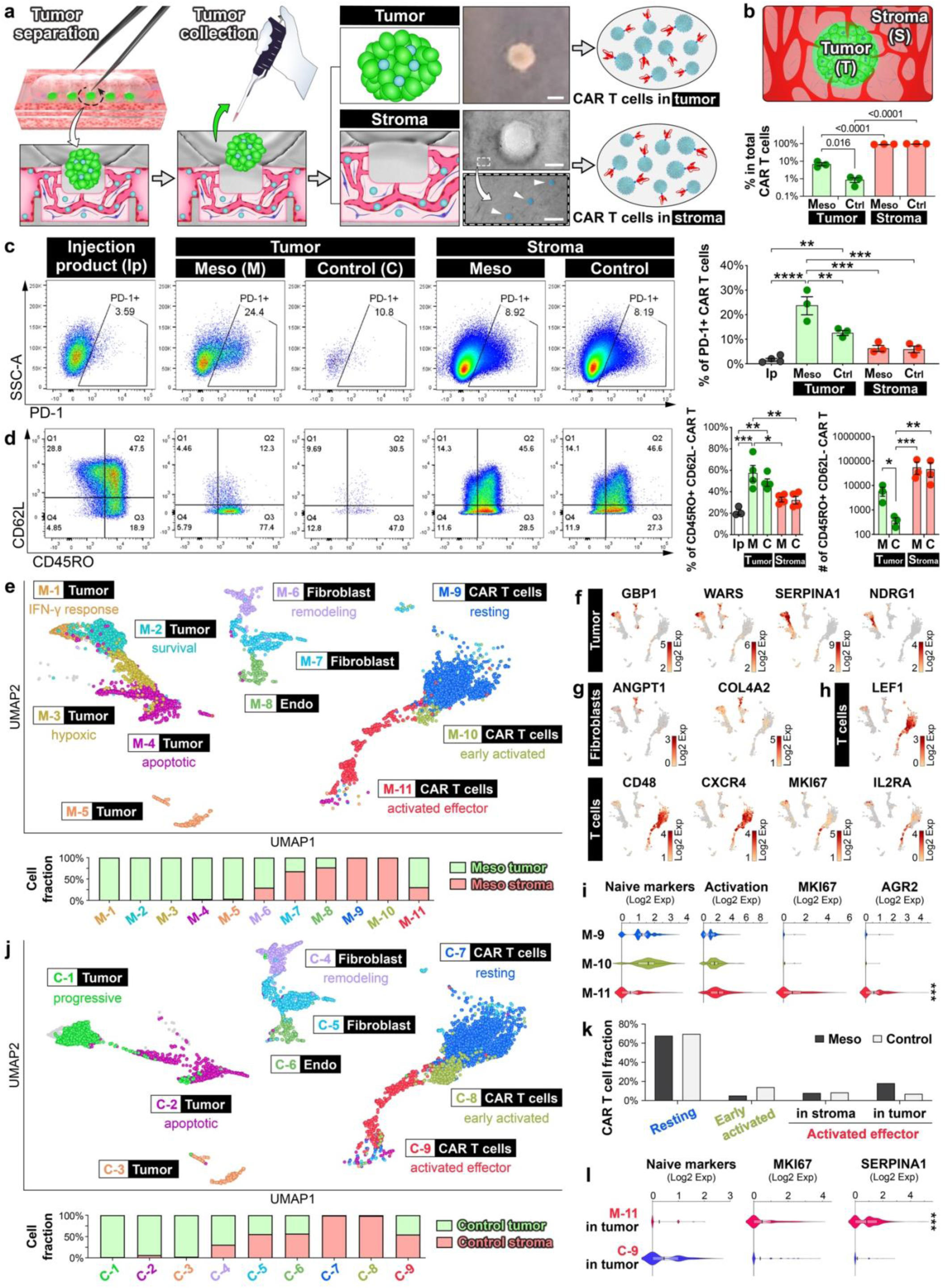
Flow cytometric analysis of CAR T cell phenotype and single-cell RNA sequencing analysis of microengineered tumor model. **a,** Illustration of workflow for separate analysis of CAR T cells harvested from tumors and those from the surrounding stroma. CAR T cells in the stroma are pseudo-colored blue and shown with white arrowheads in the phase contrast image. Scale bars, 250 µm (top, middle) and 25 µm (bottom). **b,** Quantification of CAR T cell proportions in the tumor and stromal compartments. Data are presented as mean ± SEM. *P < 0.05 and **P < 0.01 (n ≥ 3). **c-d,** Representative flow cytometry plots and quantification of surface marker expression by CAR T cells, including PD-1 (**c**), and CD45RO and CD62L (**d**). Data are presented as mean ± SEM. *P < 0.05, **P < 0.01, ***P < 0.001, and ****P < 0.0001 (n ≥ 3). **e,** UMAP projection of 11 color-coded clusters representing distinct cell subpopulations and their phenotypes in the meso-tumor model (top). Each cluster is marked with a box label indicating the number of the cluster (e.g., M-1) and the type of cells (e.g., Tumor). Underneath the box label is the annotation of cellular phenotype. The plot below the UMAP projection shows cluster-specific, relative proportions of cells quantified by their location (tumor vs. stroma). **f-h,** UMAP plots depicting the expression levels and spatial distribution of genes used to identify the subpopulations of tumor cells (**f**), fibroblasts (**g**), and CAR T cells (**h**). **i,** Violin plots showing the expression of select gene signatures by individual CAR T cell clusters. \*\*\**P* < 1e-7 vs. other clusters. **j,** UMAP projection of 9 color-coded clusters representing distinct cell subpopulations and their phenotypes in the control tumor model (top). Plot of cluster-specific, relative proportions of cells quantified by their location (bottom). **k,** Phenotype-specific comparison of the fraction of CAR T cells between meso-tumor and control groups. **l,** Violin plots of select markers and gene signatures by tumor-infiltrating activated effector CAR T cells from the meso-tumor (**M-11**) and control (**C-9**) models. \*\*\**P* < 1e-7. CAR T cells derived from one healthy donor were tested.

Using flow cytometry, we first measured the numbers of CAR T cells in different regions of the construct. The average total number of detected CAR T cells in the meso-tumor model (165,422) was similar to that in control tumors (124,223), but both groups showed substantially higher abundance of CAR T cells in the stroma in which over 90% of the total CAR T cells were found (**Fig. 3b**), illustrating tumor resistance to CAR T cell infiltration. Compared to control tumors, however, a significantly larger fraction (∼8 times) of CAR T cells was found within the tumor mass in the meso-tumor group (**Fig. 3b**). 23.9% of these cells expressed PD-1, an established marker of T cell activation that allows for identification of tumor-reactive T cells^66^, whereas the original population infused into the model contained a much smaller fraction of PD-1-positive T cells (1.8%) (**Fig. 3c**). Another significant difference in PD-1 expression was observed between the meso-tumor group and the control in which approximately 12.8% of the tumor-infiltrating CAR T cells were found to be PD-1-positive (**Fig. 3c**). The average number of these cells was 60, which was insignificant compared to that in the meso-tumor model (2,102). No significant difference was noted between these groups in the percentage of PD1-expressing CAR T cells in the stromal compartment (**Fig. 3c**). Analysis of CD69, which is another marker for T cell activation and tissue retention, showed similar patterns of compartment-specific differential expression between the groups (**Supplementary Fig. 11**).

Importantly, ∼60% of the CAR T cell infiltrates found within the meso-tumors 6 days post infusion was found to be CD45RO+ CD62L-, indicating their phenotype as effector memory T cells (T_EM_) (**Fig. 3d**). This was a substantial increase from the proportion of CAR T cells with the same phenotype detected in the original infused population (22%) (**Fig. 3d**). Similar increases in the percentage of T_EM_ were observed in the control tumors, but the total number of these cells was significantly smaller than that measured in the meso-tumor model (**Fig. 3d**). The intratumoral CAR T cell population in this group contained a large fraction of cells (∼40%) exhibiting the central memory T cell (T_CM_) phenotype (CD45RO+ CD62L+) (**Fig. 3d, Supplementary Fig. 11**). In both meso-tumor and control groups, the preferential differentiation of CAR T cells into T_EM_ was not observed in the stromal compartment (**Fig. 3d**). Our characterization also showed the phenotypic alterations of CAR T infiltrates in the meso-tumors towards CD103+ tumor-reactive tissue-resident memory T cells (T_RM_) (**Supplementary Fig. 11**), which have been shown to play important roles in anti-tumor activities of T cells and their persistence^67,68^. This cell population was negligible or accounted for minute fractions of CAR T cells in the stromal compartment surrounding the meso -tumors or in the control model (**Supplementary Fig. 11**).

Notably, the cancer-induced CAR T cell differentiation into T_EM_ and T_RM_ observed in the meso-tumor model is similar to what occurs in tumor-infiltrating lymphocytes (TILs) in mesothelioma and early-stage non-small cell lung cancer (NSCLC) patients^66,69^. Supporting our imaging data, the results of flow analysis show the activation and infiltration of the tumor antigen-encountered meso-CAR T cells infused into our model and their ability to undergo phenotypic changes required for their effector function and persistence^70,71^.

### Single-cell RNA sequencing analysis

To develop a more in-depth understanding of tumor-CAR T interactions in our models, we performed single cell RNA sequencing (scRNA-seq) of the entire CAR T cell-infused tumor constructs. For this work, cells were harvested from the microengineered tissues at Day 6 post infusion in a compartment-specific manner as described above.

Unsupervised clustering of the sequencing data from the meso-tumor model using uniform manifold approximation and projection (UMAP) yielded four clusters corresponding to three distinct groups of cells, including i) tumor cells identified by their expression of mesothelin (MSLN) and other genes indicative of their epithelial origin (e.g., AGR2) as well as their association with lung cancer (e.g., EGFR), ii) CAR T cells that express known T cell markers such as CD3E and IL7R, and iii) a mixed population of endothelial cells and fibroblasts distinguished by their expression of endothelial markers (e.g., PECAM1, VWF) and genes involved in ECM synthesis (e.g., COL1A1, TIMP1) and contractility (e.g., ACTA2, TAGLN) (**Supplementary Figs. 12a, 12b**). These different cell types were confirmed by gene ontology enrichment analysis (**Supplementary Fig. 13**). Spatial mapping of the identified clusters showed that the entire tumor cell population was associated only with tumor masses, whereas T cells, fibroblasts, and endothelial cells were found in both stromal and tumor compartments (**Supplementary Fig. 12a**).

Further analysis of cellular phenotypes in the meso-tumor model revealed a total of 11 different subpopulations (**Fig. 3e**). Specifically, the tumor cell clusters were composed of 5 subtypes. Among them are tumor cells that express hypoxia-inducible genes (e.g., NDRG1, EGLN3, HILPDA) (**M-3** in **Fig. 3e**; **Fig. 3f, Supplementary Figs. 12b, 14, 15**) or genes implicated in the survival of cancer cells, such as SERPINA1^72^ (**M-2** in **Fig. 3e**; **Fig. 3f, Supplementary Figs. 12b, 14, 15**). Importantly, immune attack by CAR T cells in this model was demonstrated by a tumor cell cluster that was uniquely identified by increased expression of WARS and GBP1 known to play a critical role in cellular responses to interferon-γ and other cytokines^73^ and by GO analysis that indicated tumor necrosis factor-mediated signaling (**M-1** in **Fig. 3e**; **Fig. 3f, Supplementary Figs. 12-15**). UMAP analysis also showed two subpopulations of fibroblasts with one cluster spatially more associated with tumor and characterized by the markers of perivascular cells that regulate vascular remodeling during cancer angiogenesis (e.g., ANGPT1, COL4A2)^74^ (**M-6** in **Fig. 3e**; **Fig. 3g, Supplementary Figs. 12-16**).

The CAR T cell group contained three clusters, all of which were enriched with CD8+ T cells (**Figs. 3e, 3h, Supplementary Figs. 12-14,17**). These subpopulations, however, exhibited distinct transcriptomic signatures reflecting different activation states and phenotypes. Specifically, the results showed resting meso-CAR T cells residing predominantly in the stroma that displayed markers of central memory CD8+ T cells (e.g., LEF1, SELL, TCF7) (**M-9** in **Fig. 3e**; **Figs. 3h, 3i, Supplementary Figs. 12-14,17-19**). These cells also expressed genes that are inducible by inflammatory cytokines (e.g., IL32, CD48, ZFP36L2) (**Fig. 3h, Supplementary Figs. 12-14,17-19**). The neighboring cluster showed an abundance of early activated cells that also expressed genes associated with cell motility and chemotaxis (e.g., CXCR4, RGS1)^75^ (**M-10** in **Fig. 3e**; **Figs. 3h, 3i, Supplementary Figs. 12-14,17-19**). Given that these cells were located in the stromal region, the cluster **M-10** likely represents CAR T cells undergoing infiltration and directional migration towards tumors.

The majority of activated CAR T cells found in Cluster **M-11** were localized to tumor masses and expressed high levels of cell-cycle genes (e.g., TOP2A, CENPF, MKI67) (**Figs. 3e, 3h, 3i, Supplementary Figs. 12-14,17-19**), illustrating their identity as tumor-infiltrating CAR T cells undergoing proliferation presumably due to antigen stimulation and activation^29^. The phenotype of this population as effector T cells was further verified by effector gene signatures and by GO analysis that indicated its capacity for interferon-gamma signaling and production (**Supplementary Figs. 13,18**). Compared to those in the other CAR T cell clusters, these cells also expressed significantly higher levels of gene markers associated with ER stress response such as AGR2 that can be induced by the hostile microenvironment of solid tumors during CAR T cell infiltration^76^ (**Fig. 3i**).

By comparison, the phenotypic variability of tumor cells in the control model was greatly reduced, and a large fraction of the cells were characterized by patterns of gene expression known to promote cancer progression (e.g., MUC5AC, CEACAM6) (**C-1** in **Fig. 3j, Supplementary Fig. 20**). In the case of CAR T cells, similar clustering was observed but the relative abundance of the subpopulations was different from that in the meso-tumor group. For instance, the early activated cell type (**C-8** in **Fig. 3j**) accounted for a substantially larger proportion of the CAR T cell population, whereas the activated effector CAR T cells (**C-9** in **Fig. 3j**) were present in much lower relative abundance (**Fig. 3k**). The tumor-infiltrating effector CAR T cells in this group also showed higher levels of naïve T cell markers and lower expression of MKI67 and SERPINA1 (**Fig. 3l**), indicating their reduced effector function, proliferative capacity, and persistence.

Further comparison using gene sets associated with T cell differentiation into effector memory (T_em_) or terminal effector T cells (T_eff_)^77^ showed that when compared to the control group, CAR T cells isolated from the meso-tumors expressed higher levels of genes known to be upregulated in T_em_ (e.g., SOCS1, IL2RA, CCL3) and lower levels of genes downregulated in T_em_ relative to T_eff_ (e.g., GNLY, GZMH, KLRB1) (**Supplementary Figs. 21, 22**), suggesting their skewed differentiation towards T_em_. Importantly, our marker-based manual classification and phenotypic characterization of meso-tumor-derived CAR T cells described here were consistent with the results of automated, unbiased cell type annotation of our CAR T cell sequencing data performed using SingleR in conjunction with published scRNA-seq of human TILs isolated from treatment-naïve NSCLC patients as a reference dataset^78^ (**Supplementary Fig. 21**).

Finally, we performed single-cell trajectory analysis using the sequencing data from the meso-tumor model to examine how the expression levels of top-ranked genes indicative of different cellular phenotypes change over pseudotime. This analysis allowed us to probe the dynamics of how the transcriptional profiles of CAR T and tumor cells in this model are regulated during their interactions (**Extended Data Fig. 1**).

### Assessment of armored CAR T cells

Among the recent trends in cancer immunotherapy is to further engineer CAR T cells through additional genetic modifications to express other proteins capable of boosting CAR T cell functions or recognizing biological signals derived from the TME^29,79^. Using this approach, researchers have developed a new generation of CAR T cells with enhanced anti-tumor activities, termed “armored” CAR T cells. In particular, one of the promising strategies under investigation is to target the chemokine system of malignant tumors by generating T cells that express not only CARs against tumor cell markers but also surface receptors for cancer-associated chemokines^29^. Animal studies have demonstrated improved tumor-targeting capacity of these cells for solid malignancies such as neuroblastoma^80^. Motivated by this body of work, we next explored whether we could use our system to model and assess the activity of such armored CAR T cells. Specifically, we sought to establish a model of malignant pleural mesothelioma in human lungs and test human mesothelin-targeted CAR T cells engineered to express CCR2 (**Fig. 4a**), a receptor for C-C motif chemokine (CCL2) known to be significantly elevated in mesothelioma patients^81^.

**Fig. 4.**
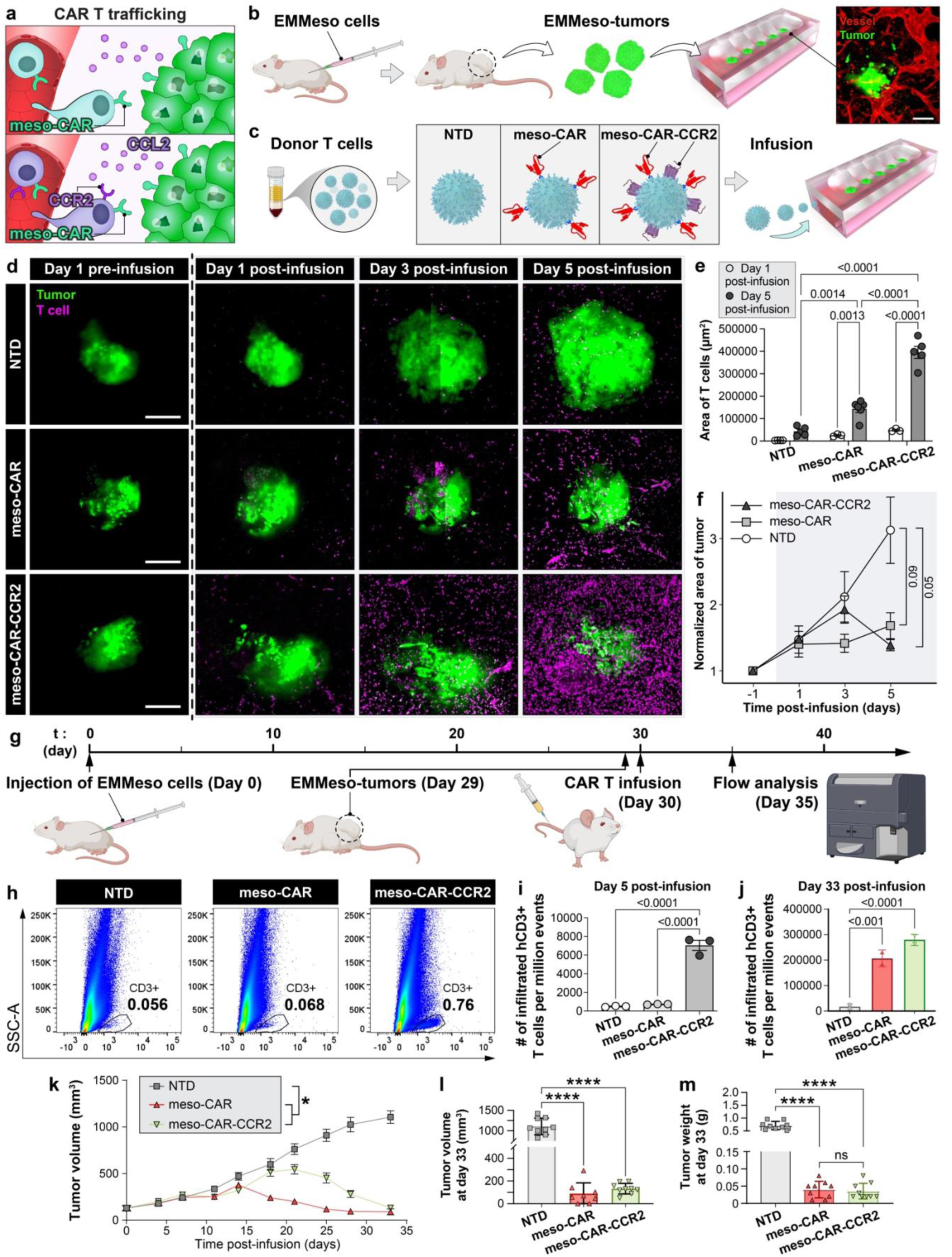
In vitro and in vivo analysis of human meso-CAR T cells armored with CCR2. **a,** Conceptual diagram illustrating (top) T cells that only express mesothelin-targeting CARs (meso-CAR, green receptor) and (bottom) those engineered to co-express meso-CARs and CCR2 (purple receptor) to recognize tumor-secreted CCL2 (purple circles). **b,c,** Experimental workflow to generate vascularized mesothelin-expressing xenograft mesothelioma tumors (EMMeso-tumors) in the device (**b**) and three types of human T cells used in our study (**c**). Scale bar, 250 µm. **d**, Representative tiled images of EMMeso-tumors prior to and after T cell infusion. Scale bars, 250 µm. The three types of T cells were derived from one healthy donor. **e**,**f**, Quantification and comparison of T cell-occupied hydrogel area (**e**) and tumor area (**f**) at different time points. Data are presented as mean ± SEM (n = 3-6). Shaded in grey in **f** is a period of T cell infusion. **g**, Experimental workflow to generate a mouse model of EMMeso-tumors for in vivo analysis of meso-CAR-CCR2 cells. Created with Biorender.com. A set of NTD, meso-CAR, and meso-CAR-CCR2 cells were derived from the same donor. T cells from two healthy donors were tested in this study. **h**-**j**, Representative flow cytometry plots (**h**) and quantification of tumor-infiltrated human CD3+ T cells at (**i**) Day 5 and (**j**) Day 33 post intravenous injection of T cells. Data are presented as mean ± SEM (n = 3). Each data point in **j** represents the average of 4-5 mice as shown in **l** and **m**. **k**, Quantification of tumor volume over time. *P < 0.05 (n = 8-9). **l**,**m**, Quantification of tumor volume (**l**) and tumor weight (**m**) at Day 33 post T cell injection. Data are presented as mean ± SEM (n = 8-9). ****P < 0.0001.

In this study, we first generated human mesothelioma tumor xenografts in NSG mice by injecting a human mesothelioma cell line called EMMeso that expresses mesothelin on the surface and secretes CCL2^34^. The EMMeso-tumors formed in the mice were then harvested and transplanted into our device containing primary human lung microvascular endothelial cell-derived blood vessels (**Fig. 4b**). Following vascularization, the tumor constructs received single-dose infusion of one of the following three types of human T cells – non-transduced T cells (NTD), meso-CAR T cells (meso-CAR), or meso-CAR T cells armored with constitutive expression of CCR2 (meso-CAR-CCR2) (**Fig. 4c, Supplementary Fig. 23**).

At day 1 post-infusion, the perfused constructs were seen with T cells identified by CD3 expression in all three groups, but quantification of their abundance in the gel and inside tumor masses did not yield any significant differences between the groups (**Figs. 4d, 4e**). Over the next 4 days, no notable changes were observed in the NTD model, whereas the other two groups showed a common trend of increasing T cell recruitment over time (**Figs. 4d, 4e**). Compared to the meso-CAR group, however, the mesothelioma constructs infused with the armored CAR T cells exhibited more extensive recruitment and retention of T cells in the tumor compartment and its vicinity at day 5 (**Figs. 4d, 4e**). Notably, these differential patterns of T cell trafficking were correlated with the growth of transplanted tumors over the course of experiments. The NTD group showed uncontrolled, rapid tumor growth, resulting in a more than 3-fold increase in the projected tumor area within 5 days of infusion (**Fig. 4f**). The meso-CAR-CCR2 model responded to the infused cells, but this occurred in a delayed manner – the tumors continued to grow at similar rates until even 3 days after CAR T cell infusion, after which they began to shrink and became roughly twice as small as their counterparts in the NTD control group by day 5 (**Fig. 4f**). The anti-tumor capacity of infused T cells was evident in the meso-CAR group as well, and it appeared that their effects were manifested as arrested tumor growth without significant tumor shrinkage (**Fig. 4f**).

These results demonstrate that mesothelin-targeted CARs indeed contribute to tumor-directed T cell recruitment in our model and that this increased T cell trafficking may be further promoted by adding CCR2 expression. To investigate the relevance of these findings to in vivo situations, we then conducted an animal study in which the same types of human T cells were tested in a xenograft mouse model of malignant mesothelioma (**Fig. 4g**). To establish this in vivo model, we injected EMMeso cells into three groups of NSG mice, after which mesothelioma tumors were allowed to form and grow to a volume of 100 - 200 mm^3^ over a period of four weeks. At this point, the same number (5 million) of NTD, meso-CAR, or meso-CAR-CCR2 cells were administered into the tumor-bearing mice via intravenous tail vein injection.

When the infused tumors were harvested from the mice for flow cytometry at day 5 post-injection, which was when our in vitro analysis was performed, data showed low abundance of intratumoral human CD3+ T cells in all three groups, presumably due to the early time point of measurement, but the proportion of these cells in the meso-CAR-CCR2 group was more than 10-fold higher than that in the NTD and meso-CAR models (**Fig. 4h**). Consistent with this result, quantification of human CD3+ T cells in the EMMeso-tumors at day 5 post-infusion revealed significantly higher cell numbers in the meso-CAR-CCR2 model (7035 per million) compared to the other two groups (486 and 721 per million for NTD and meso-CAR, respectively) (**Fig. 4i**). This difference was similar to what was observed in our in vitro model at the same time point (**Fig. 4e**). When the same measurements were taken at much later times points, for instance at day 33 post-infusion, the meso-CAR-CCR2 group still showed the largest numbers of T cell infiltrates, but substantially increased CAR T cell trafficking was observed in the meso-CAR model in which the measured cell numbers became comparable to those in the meso-CAR-CCR2 group (**Fig. 4j**). In comparison, the NTD group was seen with much smaller cell numbers (**Fig. 4j**).

These results were in agreement with temporal profiles of tumor volume measured over the course of 33 days during in vivo experiments, which demonstrated the capacity of meso-CAR-CCR2 and meso-CAR T cells to control tumor growth (**Fig. 4k**). Interestingly, despite the fact that the extent of CAR T cell trafficking was the greatest in the meso-CAR-CCR2 group even at early time points (e.g., day 5 post-infusion), measurable reduction in the tumor volume in this model was observed only after day 20 (**Fig. 4k**). At day 33 post-infusion, the average tumor volume and weight in the meso-CAR-CCR2 and meso-CAR models were approximately 88% and 94% smaller than those in the NTD group (**Figs. 4l, 4m**).

These in vivo data corroborate our in vitro result that armoring human meso-CAR T cells with CCR2 may allow them to recognize and infiltrate malignant human mesothelioma tumors more rapidly. It was also a common observation of our in vitro and in vivo studies that this accelerated cell trafficking may not necessarily result in more rapid therapeutic effects and that it may still take time for the T cell infiltrates to exert sufficient antigen-directed cytotoxicity to arrest or reverse the growth of mesothelioma tumors, although the delayed anti-tumor effects occurred over much longer periods of time in vivo. The role of CCR2 shown by our results needs to be further investigated but taken altogether, these data help us better understand how our microphysiological tumor model may be used to generate in vitro data that can correlate with in vivo activities of engineered human T cells for preclinical assessment of their therapeutic efficacy.

### Transplantation and CAR T cell infusion of tumor explants from mesothelioma patients

Having demonstrated the capabilities of our platform using genetically engineered CDX lung tumors, we next asked whether this technology could be applied to the analysis of unmodified, native tumor explants isolated from cancer patients. To make this study directly relevant to our proof-of-concept demonstration, we selected mesothelioma as a model disease. Malignant mesothelioma is an aggressive form of cancer in which overexpression of mesothelin by tumor cells has been reported in over 80% of patients^82^. This important feature of the disease has led to the development of targeted therapeutic approaches using meso-CAR T cells, many of which are being evaluated in several ongoing clinical trials^56,60^.

Malignant tumor tissues used in this study were obtained from surgical resections of multiple patients with locally advanced mesothelioma and were processed for in vitro transplantation into our device (**Fig. 5a**). To improve the physiological relevance of our original model that contained HUVEC-derived blood vessels, we used primary human lung microvascular endothelial cells to form the perfusable vascular bed in the device (**Fig. 5a**). Following transplantation, the blood vessels in the surrounding tissue grew into the open wells and covered the primary mesothelioma explants over time, generating fully vascularized and perfusable tumor constructs within 10 days (**Fig. 5b**). The extent of vascularization of patient tumor explants during the first few days of transplantation was similar to when CDX tumors were transplanted into the same system, but at the end of the vascularization period for 10 days, our data indicated the formation of significantly larger and more connected blood vessels in the patient tumor model (**Fig. 5c**).

**Fig. 5.**
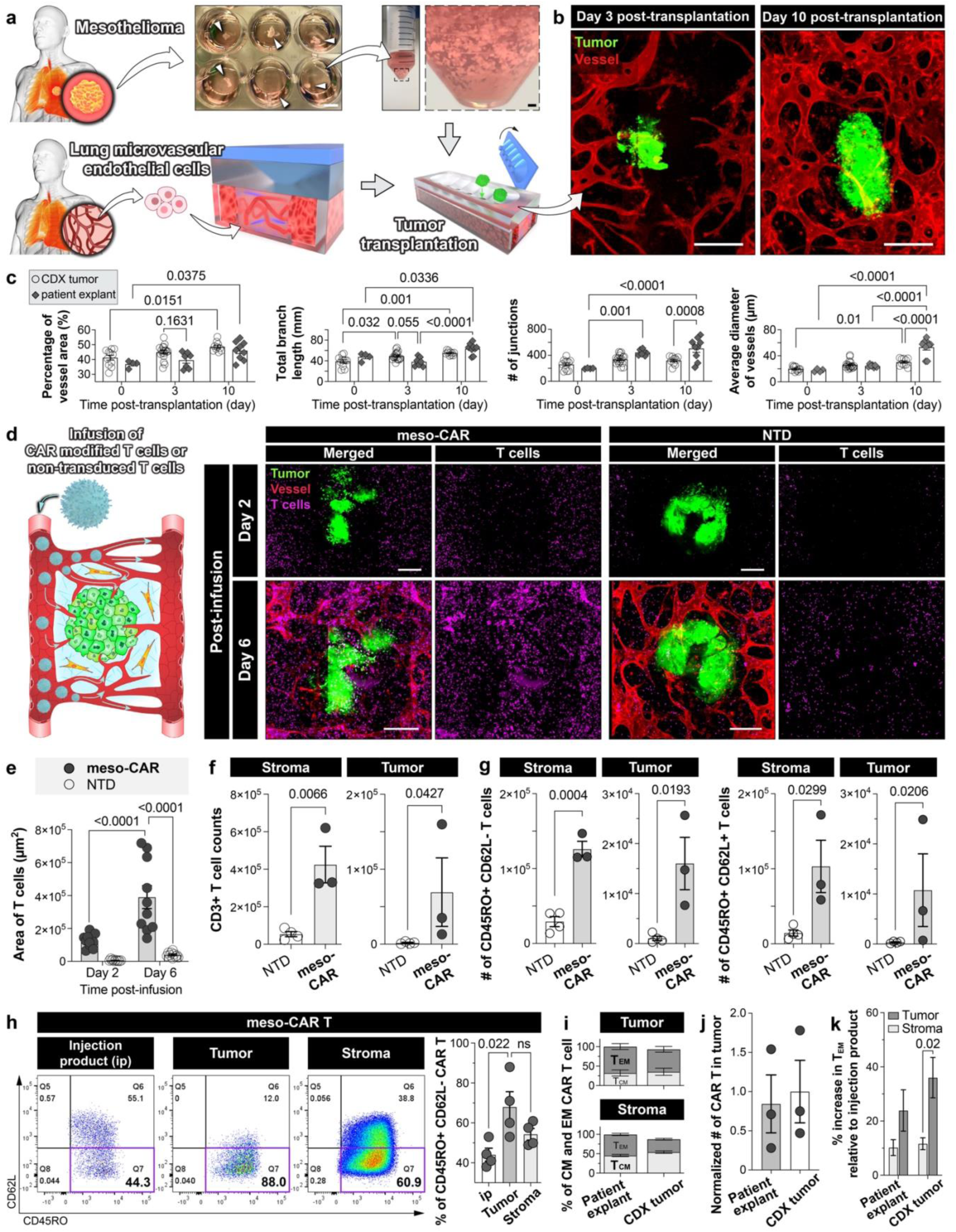
Transplantation and CAR T cell treatment of human mesothelioma explants. **a,** Workflow of in vitro transplantation of primary tumor explants surgically resected from mesothelioma patients and their vascularization using primary human pulmonary microvascular endothelial cells. The photos in the top row show tumor slices in a 24-well plate (arrowheads, left) and minced, suspended explants in a conical tube ready for transplantation (right). Scale bars, 5 mm (left) and 1 mm (right). **b,** Representative fluorescence images of tumor explants during vascularization. Scale bars, 250 µm. **c**, Quantification of the percentage of vessel-covered area, total branch length (per engineered construct), total number of vessel junctions (per engineered construct), and the average diameter of microvessels in the CDX tumor (open circle) and patient-derived explant (closed diamond) models during tumor vascularization in our device. Data are presented as mean ± SEM (n = 4-18). **d**, Representative tiled micrographs showing vascularized tumor constructs infused with non-transduced T cells (NTD) or meso-CAR T cells (meso-CAR). Scale bars, 250 µm. **e**, Quantification and comparison of T cell infiltration into the tumor compartment and surrounding stroma within the regions of interest. Data are presented as mean ± SEM. (n = 10-11). **f,g,** Tissue compartment-specific flow cytometric analysis of CD3+ T cells (**f**) and CD45RO+ CD62L-(effector memory) or CD45RO+ CD62L+ (central memory) CD8+ T cells (**g**) at Day 6 post-infusion. Data are presented as mean ± SEM. (n = 3-4). **h**, Representative flow cytometry plots of CD45RO and CD62L expression by meso-CAR T cells harvested at Day 6 post-infusion. Data are presented as mean ± SEM. (n = 4). CAR T cells derived from two healthy donors were tested in this study. **i-k**, Comparison of the relative ratio of effector memory to central memory CAR T cells (**i**), the number of tumor-infiltrating CAR T cells (**j**), and the increase in the proportion of effector memory CAR T cells post-infusion in the tumor and stroma compartments relative to injection product (**k**) between the patient explant- and CDX tumor-derived models. Data are presented as mean ± SEM (n = 4 in **i** and **k**, n = 3 in **j**).

When meso-CAR T cells were injected into the vascularized constructs, only small numbers of the cells were found in the model initially (**Figs. 5d, 5e**), which was in contrast to rapid trafficking and accumulation of CAR T cells observed in the CDX meso-tumor model within 5 hours (**Fig. 2**). During prolonged experiments, however, the recruitment of the infused cells into the tumor explants and their vicinity became more pronounced, as evidenced by significantly increased area of CAR T cell infiltration over time (**Figs. 5d, 5e**). In control experiments using non-transduced T cells (NTD) infused under identical conditions, the extent of T cell trafficking appeared to be similar to that in the meso-CAR T group initially, but it became evident over time that these cells were greatly limited in their capacity to interact with the tumors and their associated vasculature, which was illustrated by significantly lower levels of T cell adhesion and infiltration in the control group (**Figs. 5d, 5e**).

Verifying the imaging data, flow cytometry at Day 6 post-infusion showed significantly higher abundance of T cells in the stroma and tumors when the model was infused with meso-CAR T cells (**Fig. 5f**). These cells were also measured to be more capable of anti-tumor activity than their non-transduced counterparts, which was illustrated by substantially larger numbers of effector memory (CD45RO+ CD62L-) and central memory (CD45RO+ CD62L+) T cells in both tissue compartments of the meso-CAR T group (**Fig. 5g**). Our data indicated that approximately 70% of the CAR T cells retrieved from the tumor explants assumed the effector memory phenotype, and this was a significant increase from the proportion of the same cell type measured in the injection product (44%) (**Fig. 5h**), suggesting tumor-induced differentiation of the significant portion of the original population into effector memory T cells. In the stroma, CD45RO+ CD62L-CAR T cells accounted for roughly 54% of the total population, which was statistically insignificant from the fraction of the same cell type in the tumor compartment (**Fig. 5h**).

These results allowed us to compare the behavior of meso-CAR T cells in the patient-derived mesothelioma explants used in this study to that in the mesothelin-overexpressing CDX tumors used in the proof-of-concept demonstration of our platform (**Figs. 2-4**). First, these models showed similar compartment-specific ratios of effector memory T cells (T_EM_) to central memory T cells (T_CM_) (**Fig. 5i**). In particular, the skewed differentiation of tumor-infiltrating CAR T cells towards T_EM_ was observed in both systems and occurred to a similar extent. We also noted that the total number of tumor-infiltrating CAR T cells by the end of culture was comparable between the mesothelioma explants model and the CDX tumor model (**Fig. 5j**). A notable difference, however, was that in the CDX tumor model, the increase in the proportion of T_EM_ compared to the injection product was significantly higher in the tumor compartment than in the stroma, whereas this difference was insignificant in the patient explants model (**Fig. 5k**), which we think may be attributed to heterogenous architecture and cellularity of the patient tumor explants, as well as greater variability of data obtained from these tissues.

Although limited to testing meso-CAR T cells for mesothelin-expressing tumors, this comparison suggests that the CDX tumors may have the capacity to simulate their human in vivo counterparts for the purposes of studying tumor-directed trafficking of CAR T cells and their phenotypic changes necessary for anti-tumor activity. It should be noted, however, that the differences shown by this comparison are equally important in that they illustrate the difference between such CDX tumors engineered in a controlled environment to acquire uniformly high expression of target proteins and patient tumors that are intrinsically variable in tumor antigen expression and other biochemical properties.

### Discovery of therapeutic targets to promote CAR T cell trafficking

As part of ongoing efforts to advance CAR T therapy, increasing attention is being paid to combining CAR T cells with other treatment modalities (e.g., radiotherapy) or therapeutic agents designed to help improve their anti-tumor activities and safety profiles^79^. Research has shown, for example, that monoclonal antibodies targeting PD-1 and PD-L1 can enhance the persistence and anti-tumor function of CAR T cells in glioblastoma and other types of solid malignancies^18,26,83^, which is now under investigation in several clinical trials^26,79,83^. A growing body of evidence demonstrating the promise of this and other combinatorial strategies led us to wonder if our technology would offer any value for this line of investigation.

Specifically, we asked whether we could use our microengineered system to i) probe and interrogate complex biological interactions present in the microengineered tumor transplants infused with CAR T cells, ii) identify molecular pathways that play an important role in regulating the crosstalk between CAR T cells and the cellular constituents of the tumor constructs, and iii) explore the possibility of pharmacologically modulating them for the purposes of improving the trafficking and anti-tumor function of CAR T cells. For this work, we leveraged the scRNA-seq data acquired from the lung adenocarcinoma models used for our proof-of-concept demonstration (**Figs. 1-3**) and sought to further analyze the data using CellPhoneDB to predict enriched intercellular communication mediated by ligand-receptor complexes^84^ (**Fig. 6a**).

**Fig. 6.**
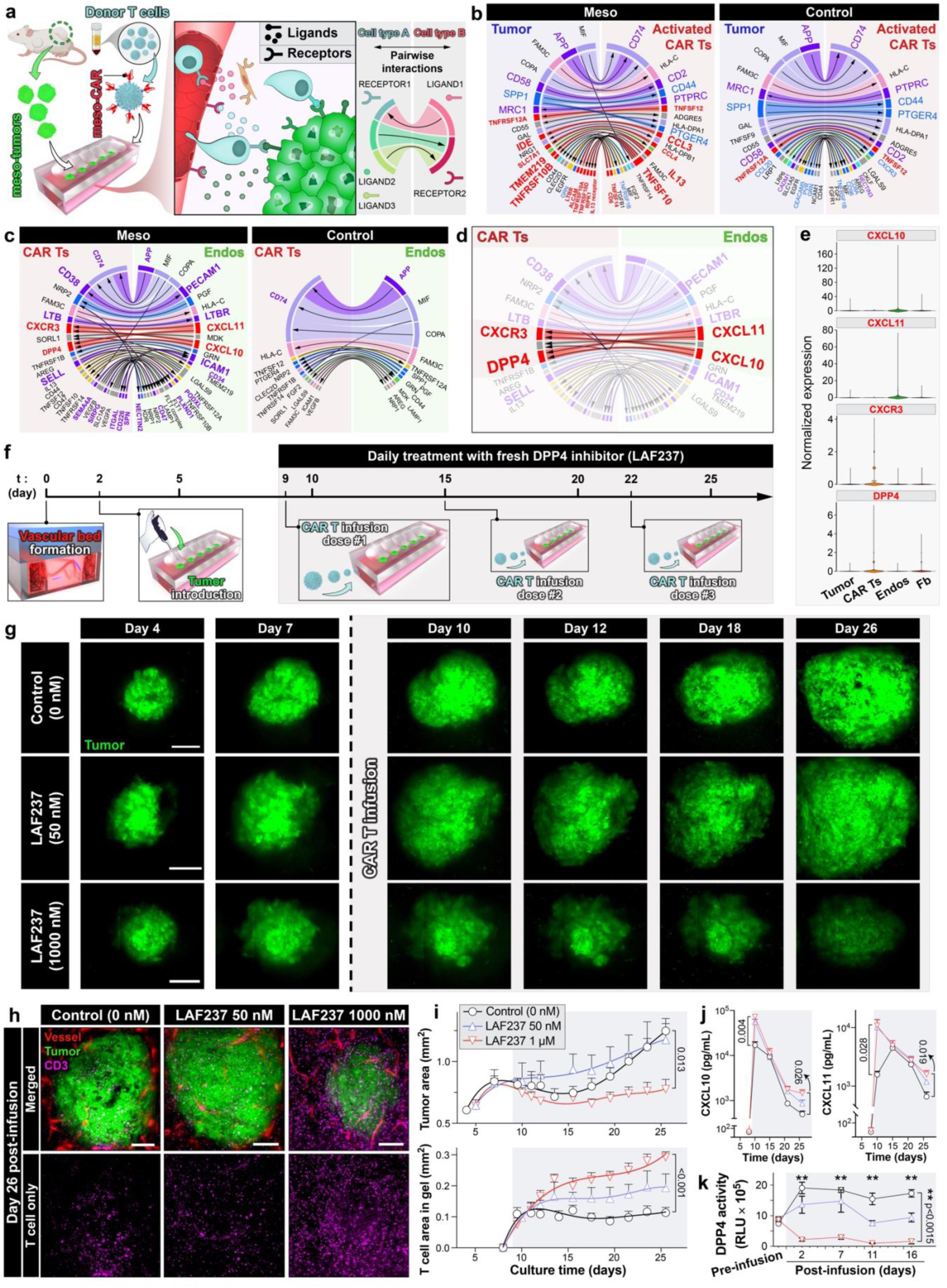
Identification of new therapeutic targets through analysis of ligand-receptor interactions in CAR T cell-treated tumor models. **a,** Conceptual diagram illustrating pairwise mapping of directional ligand-receptor interactions between different cell types in the engineered tumor model. **b,c,** Chord diagrams of cell pair-specific ligand-receptor interactions with adjusted p-values < 0.05 and total mean expression > 0.35. **d,** Visualization of interactions between CAR T cells (CAR Ts) and endothelial cells (Endos) mediated by CXCR3/DPP4-CXCL10/CXCL11. **e,** Violin plots comparing the expression of these mediators across all cell types. **f,** Experimental timeline for CAR T cell infusion and drug treatment. **g,** Representative confocal micrographs of single tumors treated with CAR T cells and different concentrations of LAF237. Blood vessels are not shown in these images. Scale bars, 250 μm. **h,** Representative images of CAR T cell-infused tumors at Day 26. Scale bars, 200 µm. **i,** Quantification and comparison of tumor area (top) and CAR T cell-occupied hydrogel area (bottom). Shaded in grey is a period of CAR T cell infusion and daily LAF237 treatment. Data are presented as mean ± SEM (n ≥ 3). **j,k,** Quantification of CXCL10 and CXCL11 levels (**j**) and the activity of DPP4 (**k**) in device effluent. Shaded in grey is a period of CAR T cell infusion and daily LAF237 treatment. Data are presented as mean ± SEM in (**j**) and as mean ± SD in (**k**) (n ≥ 3). CAR T cells derived from three healthy donors were tested in this study.

The results showed that the mesothelin-overexpressing and control tumor models each contained more than 150 directional ligand-receptor interactions with statistically significant (p < 0.05) cell-type specificity (**Extended Data Fig. 2a, Supplementary Fig. 24, Supplementary Table 1**), which permitted pairwise analysis of the crosstalk between two different cell types. In case of the activated CAR T cell-tumor cell pair, for example, our data showed common interactions observed in both the meso-tumor and control groups, such as APP-CD74, PTPRC-MRC1, and CD58-CD2 (purple chords in **Fig. 6b**), all of which are known to be involved in mediating T cell adhesion to tumor cells^85–87^. In comparison to the control group, however, the meso-tumor model contained substantially greater numbers and extent of interactions involved in functional activation and cytotoxicity of “on-target” T cells that are mediated by binding of T cell-produced ligands to their receptors on cancer cells (e.g., CCL3-IDE, IL13-TMEM219, TNFSF10-TNFRSF10B) (red chords in **Fig. 6b, Supplementary Fig. 24**). The obtained data also permitted visualization and close examination of various directional signaling activities between the other cell-type pairs (**Extended Data Figs. 2b-2e**).

In defining the scope of further analysis, we chose to focus on the endothelial-CAR T cell crosstalk, which represents the first critical step of CAR T cell infiltration into the tumor microenvironment. Despite the established importance of the vascular endothelium as an essential regulator of immune cell recruitment^88–91^, the idea of targeting and modulating the biological activity of this specialized tissue has not been explored as extensively in CAR T therapy where current research and clinical practice focus heavily on the principle of targeting antigens expressed by malignant cancer cells^18,25^. Therefore, we aimed to investigate i) whether our analysis could generate any new biological insight into how the interaction of CAR T cells with the tumor-associated vasculature is regulated in our model and ii) whether this knowledge could be exploited for developing new strategies that could be combined with CAR T cells to enhance their therapeutic potential.

In comparison to the control group, our data in the meso-tumor group showed well-known, classical signaling, such as ITGAL-ICAM1, known to play a crucial role in endothelial adhesion of T cells during their recruitment (**Fig. 6c, Extended Data Fig 3**). Overall comparison of the predicted interactions made it clear that the endothelial-CAR T cell crosstalk was much more active in the meso-tumor model (**Fig. 6c**). Among the top-ranked ligand-receptor complexes in this group with even higher levels of significance than ICAM1-ITGAL were pairs of i) CD38 on CAR T cells and its ligand PECAM1 produced by endothelial cells (**Fig. 6c**, **Extended Data Fig 3**) and ii) lymphotoxin-beta receptor (LTBR) expressed by endothelial cells and its ligand lymphotoxin-beta (LTB) from CAR T cells, both of which were not significant in the control group (**Fig. 6c, Extended Data Fig 4**). Importantly, when the activity of these pathways was suppressed by treating the meso-tumor model with Daratumumab, a monoclonal antibody drug that can bind to CD38 ^92^, or Baminercept, which is a lymphotoxin β receptor IgG fusion protein ^93^, during and after CAR T cell infusion, we indeed observed significant reduction in tumor-directed trafficking of meso-CAR T cells and their effects on tumor growth (**Extended Data Figs. 3, 4**), validating the importance of these endothelial-CAR T cell signaling axes predicted by our ligand-receptor analysis.

Interestingly, the top-ranked and most notable interactions predicted in the meso-tumor model also included the crosstalk between endothelial CXCL10/11 and CXCR3 expressed by CAR T cells (**Figs. 6d, 6e**), which was absent in the control group. Binding of CXCL10/11 produced by IFN-γ-stimulated endothelium to CXCR3 expressed on T cells is a well-known interaction that induces chemotaxis of T cells and their endothelial trafficking^94,95^. It was interesting, however, that CXCL10/11 was also interacting with DPP4 produced by CAR T cells and fibroblasts in our model (**Figs. 6d, 6e**) – DPP4 is a receptor protein known to be secreted by T cells and certain types of fibroblasts that can bind to CXCL10/11 and cleave the N-terminal peptides of these chemokines, impairing their ability to mediate T cell recruitment and chemotaxis^96^. Based on this finding, we asked whether pharmacological inhibition of DPP4 could be used as a strategy to preserve the bioactivity of CXCL10/11 and thus promote the trafficking of meso-CAR T cells, which had not been investigated previously.

To test this idea, we modified the meso-tumor infusion model as follows. First, we decreased mesothelin expression by A549 cells from 88% to 44% by reducing the fraction of mesothelin-positive cells in the tumor constructs, in order to more closely approximate the pathophysiological levels of mesothelin expression in malignant lung tumors in vivo^97,98^. Second, we changed the regimen of CAR T cell infusion from a single dose of highly concentrated cells to three doses with lower cell density separated by 6-7 days to simulate the typical regimen of fractioned administration of engineered human TCR/CAR T cells for efficacy testing in preclinical and clinical settings (**Fig. 6f**)^35,99,100^. Lastly, the total culture period was extended to 26 days to allow for longer-term monitoring and assessment of CAR T cell activities. Importantly, we used LAF237 (Vildagliptin) as a pharmacological inhibitor of DPP4 in our experiments – LAF237 is a potent DPP4 inhibitor previously developed for the treatment of type 2 diabetes^101,102^. Given the short half-life of this drug, our model was administered with a single daily dose of LAF237 for the entire duration of experiments after CAR T cell infusion (from days 9 to 26) (**Fig. 6f**)^101^.

Prior to the introduction of CAR T cells, the lung tumors transplanted into the vascularized bed continued to grow in size over time (**Fig. 6g, Supplementary Fig. 25**). Following the perfusion of the tumor constructs with meso-CAR T cells, this trend appeared to stop as illustrated by the slightly reduced tumor area between days 12 and 18 but they began to grow again after the second infusion and subsequently underwent rapid enlargement (top row in **Fig. 6g**, **Fig. 6i, Supplementary Fig. 25, Supplementary Movie 9**). Similar growth patterns were observed in CAR T cell-infused tumors treated with 50 nM of LAF237, a dose representing the lower end of clinically-relevant plasma concentrations of this drug^103,104^ (middle row in **Fig. 6g**, **Fig. 6i, Supplementary Fig. 25, Supplementary Movie 10**). When a higher dose (1000 nM) of LAF237 was used, however, the tumors began to shrink shortly after the first infusion of CAR T cells, after which their size remained relatively constant (bottom row in **Fig. 6g**, **Fig. 6i, Supplementary Fig. 25, Supplementary Movie 11**). Of note, at 50 and 1000 nM, LAF237 alone without CAR T cell infusion did not have any significant effects on tumor growth (**Supplementary Fig. 26**).

Importantly, immunofluorescence imaging of tumor constructs at Day 26 revealed a marked difference in the distribution of infused T cells between these groups, which was highlighted by much more extensive recruitment and retention of T cells in the tumor compartment and its vicinity when the model was treated with 1000 nM of LAF237 (**Figs. 6h, 6i, Supplementary Fig. 25**). Measurement of intra-stromal and intra-tumoral T cell area over time showed that significant T cell trafficking and infiltration occurred in all three groups after the first infusion but in the absence of or at the low dose of LAF237, the abundance of CAR T cells in the constructs remained largely unchanged even after the second and third infusions (**Fig. 6i**). In contrast, the higher dose of the drug permitted a continuous increase in the T cell area (**Fig. 6i**), suggesting the ability of LAF237 to potentiate therapeutic effects of additional doses of CAR T cells after the initial infusion.

Of note, the increased CAR T cell trafficking and controlled tumor growth observed at 1000 nM of LAF237 was abrogated when the model was treated with anti-CXCR3 antibody (**Supplementary Fig. 27**). Considering that CXCR3 is the unique receptor for CXCL10 and CXCL11 for T cells and that CXCL10 and CXCL11 are the only two ligands for CXCR3 found to be significant in our model, this result indicates that the beneficial effects of LAF237 shown here are indeed mediated through CXCR3-CXCL10/11 signaling, supporting our proposed mechanism of drug treatment.

Analysis of device effluent using ELISA indicated that the concentration of CXCL10 and CXCL11 spiked due to the first infusion of CAR T cells and then gradually decreased over time in all three groups (**Fig. 6j**). However, the levels of these chemokines in the drug-treated constructs were higher, especially at the beginning and towards the end of post-infusion culture, compared to those measured in the untreated control model (**Fig. 6j**), supporting the observation of increased CAR T cell recruitment in these groups. Recognizing that ELISA can detect both the cleaved and intact forms of CXCL10 and CXCL11 and may therefore not provide accurate measurement of their contribution to CAR T cell trafficking, we also performed ultraviolet-visible (UV-vis) and Raman spectroscopy of the effluent samples using engineered gold nanoparticles functionalized with valine-specific aptamers that can recognize and bind to dipeptides cleaved from the intact chemokine molecules (**Supplementary Fig. 28a**). Focusing on CXCL10, this analysis showed reduced truncation of the signaling chemokine when the CAR T cell-infused model received 1000 nM of LAF237 as compared to the control group without drug treatment (**Supplementary Figs. 28a-28c**), which was confirmed by western blot analysis (**Supplementary Figs. 28d, 28e**).

When we evaluated DPP4 activity in the same effluent samples, data from the untreated control group indicated that the level of DPP4 activity greatly increased by the first CAR T cell infusion and remained elevated throughout culture (**Fig. 6k**). When the model was administered with LAF237, however, the measured activity of DPP4 was lowered significantly in a dose-dependent manner, resulting in 35.4% (50 nM) and 89% (1000 nM) reduction in average activity during post-infusion culture when compared to the control group (**Fig. 6k**).

These results altogether demonstrate the potential of LAF237 as a pharmacological agent to enhance anti-tumor capacity of meso-CAR T cells in our model, which may provide a basis for developing potentially promising combinatorial approaches for CAR T cell therapies for malignant lung adenocarcinomas. As validated by our data, the proposed method relies on the ability of LAF237 to modulate the endothelial-CAR T cell crosstalk by suppressing the enzymatic activity of DPP4 to preserve endothelial CXCL10/11 and their interactions with CXCR3 on T cells.

### Discovery of therapy biomarkers through metabolomic analysis of tumor-on-a-chip

Having demonstrated the therapeutic potential of inhibiting DPP4 in meso-CAR T treatment, we then explored the possibility of developing biomarkers that indicate the efficacy of this method. In light of increasing evidence pointing to the importance of metabolism in cancer immunotherapy^105,106^, our goal was to identify metabolic signatures that can be correlated with the progression and outcome of meso-CAR T therapy coupled with the use of LAF237 in our model. To this end, we conducted untargeted, global metabolomic profiling of our model using vascular perfusate collected from the side chambers (**Extended Data Fig. 5a**). Specific conditions considered in this study included media only, pre-infusion, CAR T cell infusion only (**ctrl**), CAR T cell infusion with 50 nM LAF237 (**low**), and CAR T cell infusion with 1000 nM LAF237 (**high**) – the CAR T cell-infused groups were examined at defined time points (days 2, 7, 11, and 16 post-infusion) (**Fig. 7a, Supplementary Table 2**).

**Fig. 7.**
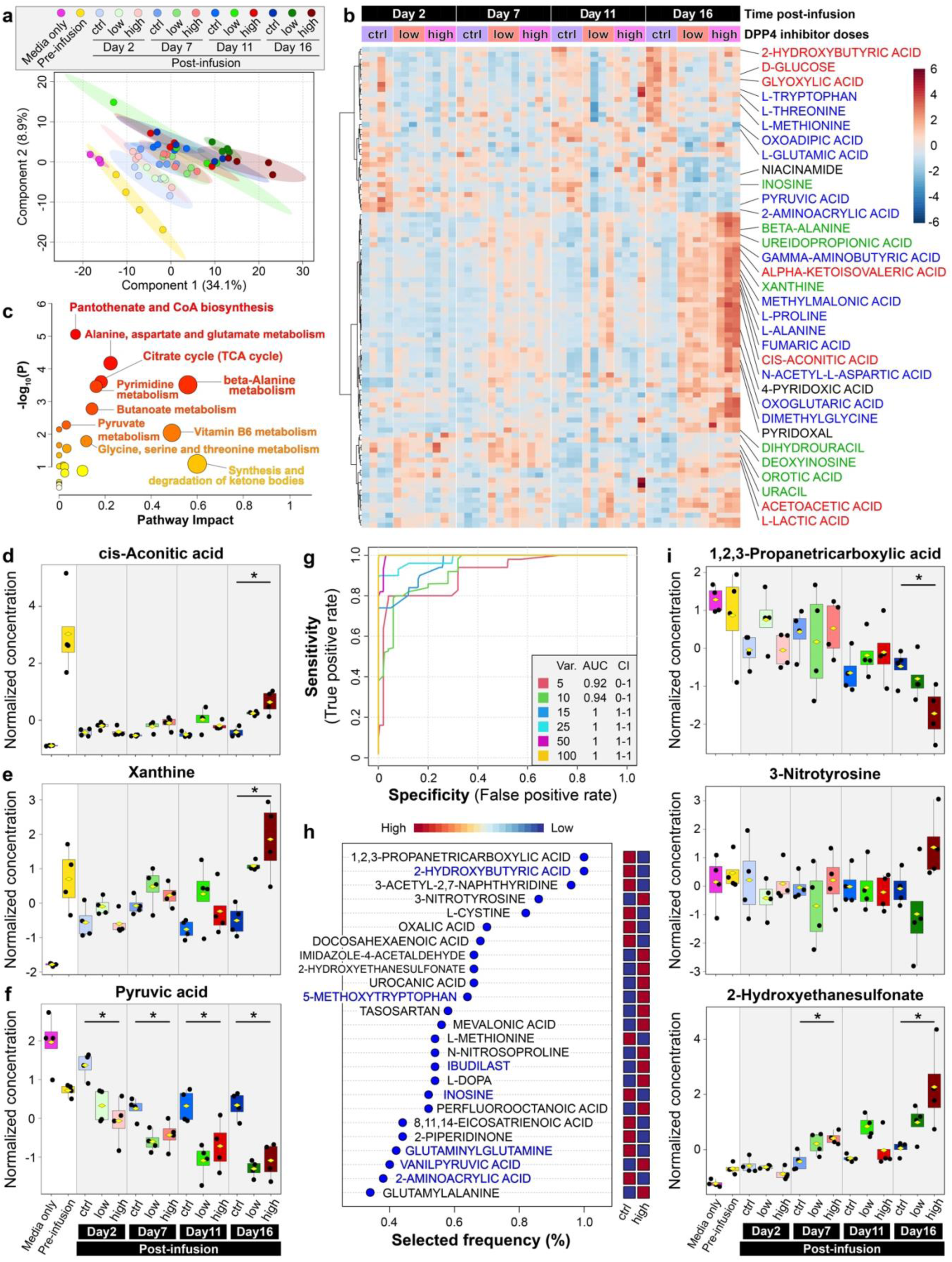
Metabolomic analysis of lung tumors treated with CAR T cells and LAF237. **a**, Plot of PLS-DA scores. N = 4 for each group. **b**, Heatmap of metabolites whose levels of expression were significantly altered by LAF treatment. N = 4 for each group. The source of metabolites is color-coded in their labels – blue, amino acid metabolism; red, carbohydrate metabolism; green, nucleotide metabolism. The color gradient of the scale bar indicates the relative abundance of metabolites, with red and blue indicating higher and lower concentrations, respectively. p < 0.05 was considered significant. **c**, Plot of significantly changed metabolic pathways identified by pathway impact analysis of significantly upregulated metabolites in the high-dose LAF237 group. **d-f**, Comparison of normalized concentrations of select metabolites across all conditions. **g**, ROC curves for biomarker prediction models to identify metabolites that can differentiate more efficacious CAR T cell treatment with high-dose LAF237 from control CAR T cell treatment without LAF237 at day 16 post infusion. Different numbers of constituent metabolite features per model are indicated by different colors. **h**, List of top metabolites yielded by the 25-feature prediction model for day 16 post infusion shown in **g** and ranked according to their predictive accuracy. The metabolites labelled blue also appear in the list of top metabolites predicted for all time points shown in **Extended Data Fig. 5d**. **i**, Comparison of normalized concentrations of metabolites selected from **h**. Boxplots show minimum, 25th percentile, mean, 75th percentile, and maximum. N = 4 for each group. Note that post-hoc pairwise comparison was calculated between the high-dose (high) and control (ctrl) groups at all time points with *p < 0.05.

This analysis revealed 205 differentially regulated metabolites with statistical significance (False Discovery Rate (FDR) < 0.05) across all conditions (**Supplementary Fig. 29**). Further analysis by partial least squares discriminant analysis (PLS-DA) identified a fraction of these differentially regulated metabolites that can be used to distinguish individual conditions with different time points and doses of LAF237 (**Fig. 7a; Supplementary Fig. 30**). Importantly, through subsequent multivariate analysis, we identified 96 metabolites whose concentrations in the vascular perfusate were significantly altered by LAF237 treatment (**Fig. 7b**). The hierarchical clustering-based heatmap showed that 65 of these metabolites were present in higher abundance in the drug-treated groups compared to control (CAR T cell infusion only), whereas the other 31 metabolites were downregulated due to LAF237 (**Fig. 7b**). According to metabolic pathway impact analysis, the upregulated metabolites had significant effects on biological processes involved in beta-alanine metabolism, alanine, aspartate and glutamate metabolism, citrate cycle, pyrimidine metabolism, and pantothenate and CoA biosynthesis (**Fig. 7c, Supplementary Fig. 31, Supplementary Table 3**), suggesting significant changes in central carbon metabolism due to DPP4 inhibition.

To highlight a few key findings, certain products of carbohydrate metabolism, such as cis-aconitic acid in the TCA cycle, were detected at significantly elevated levels at day 16 post-infusion when the model was treated with a higher concentration (1000 nM) (**Fig. 7d**). At the same time point, the higher dose treatment was correlated with increased metabolism of amino acids and nucleic acids as demonstrated by significantly greater production of intermediate metabolites of these pathways, including xanthine, L-proline, fumaric acid, and beta-alanine (**Fig. 7e, Extended Data Fig. 5b**). Our data also revealed a negative correlation between the production of pyruvic acid and drug treatment (**Fig. 7f**).

These results prompted us to ask whether the differentially regulated metabolites found in our analysis would have any value as molecular indicators of the efficacy of inhibiting DPP4 activity during meso-CAR T therapy. For systematic investigation of this question, we performed biomarker analysis to identify metabolite-based signatures that can be used to quantitatively distinguish the CAR T cell-infused group receiving an efficacious dose (1000 nM) of LAF237 from that without LAF237 treatment. In this analysis, we generated the receiver operating characteristic (ROC) curves with increasing numbers of constituent model features for the last time point of measurement (day 16 post-infusion) to predict combinatorial biomarkers of efficacy (**Fig. 7g, Supplementary Fig. 32**). For the top five metabolites identified by this analysis (**Fig. 7h**), our model predicted that higher efficacy of CAR T therapy due to 1000 nM LAF237 is correlated with significantly lower levels of 1,2,3-propanetricarboxylic acid, 2-hydroxybutyric acid, 3-acetyl-2,7-naphthyridine, L-cystine and increased production of 3-nitrotyrosine and 2-hydroxyethanesulfonate at (**Fig. 7i, Supplementary Fig. 33**).

When our analysis was extended to all time points measured in our experiments (days 2, 7, 11, and 16 post-infusion), the model predicted pyruvic acid, ibudilast, vanilpyruvic acid, zymonic acid, and 2-hydroxybutyric acid as the five most significant biomarkers (**Extended Data Figs. 5c, 5d, Supplementary Fig. 32**). In case of pyruvic acid and 2-hydroxybutyric acid, for example, their concentrations in the CAR T cell-infused group with 1000 nM LAF237 was substantially lower than that in control at days 11 and 16 post-infusion, whereas the same condition/group was seen with a significant increase in the level of ibudilast at days 7 and 16 post-infusion (**Fig. 7f, Extended Data Fig. 5e**). Notably, the two sets of predictions made for day 16 post-infusion only (**Fig. 7h**) and all time points (**Extended Data Fig. 5d**) shared several metabolites that were upregulated (5-methoxytryptophan, ibudilast, vanilpyruvic acid) or downregulated (2-hydroxybutyric acid, inosine, glutaminylglutamine, 2-aminoacrylic acid) due to administration of 1000 nM LAF237 during CAR T cell infusion (**Supplementary Fig. 33**).

## Discussion

As our understanding of malignant solid tumors and their complex ecosystem continues to evolve, significant efforts are being made to use this knowledge to design new CARs and related cellular engineering strategies to enhance therapeutic potential of engineered T cells against solid malignancies^18,79,83^. With increasing efforts towards rapid, successful translation of these new generations of CAR T cells into effective clinical therapies, one of the key challenges in the field is how to preclinically assess, predict, and optimize their ability to find and engage their targets protected by the complex tumor microenvironment and produce intended cytotoxicity in human-relevant conditions, which is an essential step for validating or improving the efficacy and safety of CAR T cells before clinical trials. Our work is significant in that the demonstrated technology may provide a potentially powerful alternative to conventional preclinical models for these types of studies and also allow us to achieve greater depth and breadth of human-relevant preclinical data.

Developing preclinical models for the study of adoptive cell therapy is a new area of investigation in cancer immunotherapy where engineering design principles and techniques have made great contributions to important research advances in recent years^107–113^. Particularly relevant to our current study is a growing body of work focused on developing organotypic 3D culture models in microfabricated devices to investigate cancer-immune interactions. For example, researchers have demonstrated a microengineered 3D culture system containing a micropatterned collagen hydrogel construct laden with hepatocellular carcinoma cells grown as dispersed single cells or 3D aggregates for in vitro analysis of human T cells engineered to express tumor-specific T cell receptors^114^. By altering the environment of the cultured tumor cells in a precisely controlled manner, this model showed the adverse effects of hypoxia commonly present in the tumor microenvironment in vivo on tumor-killing capacity of the engineered T cells, which was not observed in conventional 2D well-based assays using the same types of cells. The microengineered system also permitted in vitro modeling and investigation of adoptive T cell therapy in clinical scenarios involving the use of immunosuppressants (e.g., after organ transplantation), revealing the ability of the engineered T cells to retain their anti-tumor activities during stable immunosuppression achieved by low doses of mTOR inhibitors. In a more recent study, a multilayered microfluidic device was developed as an in vitro platform to culture a monolayer of endothelial cells with breast cancer spheroids or patient-derived breast tumor organoids grown in a 3D environment to model the endothelial-tumor interface^115^. This microphysiological system was used to examine responses of the engineered breast tumor tissues to CAR T cells engineered to target a tumor-associated antigen called receptor tyrosine kinase-like orphan receptor and measure the kinetics of proinflammatory cytokine release during CAR T cell infusion and interaction with the breast tumor constructs for up to 8 days. This specialized model also provided an in vitro platform to test a tyrosine kinase inhibitor as pharmacological intervention of cytokine release syndrome during CAR T therapy.

Our work adds to this emerging body of research by demonstrating the feasibility of integrating a living human tissue explants retaining the complexity of in vivo malignancies with a microengineered system to generate in vitro-in vivo hybrid tumor constructs that can be maintained and directly observed for prolonged periods while also being perfused, stimulated, and sampled in a controlled manner. We believe that this approach represents an important advance as it allows us to achieve higher levels of physiological realism in modeling human solid tumors, especially their interface with blood vessels and vascular supply of blood-borne cells in the surrounding environment, which is essential for the study of cancer-immune interactions. Another distinct aspect of our work is the demonstration of data-driven studies aimed at exploring the breadth, depth, and utility of in vitro data generated by our model. With rapidly increasing interest in microphysiological models of CAR T therapy, whether they can actually generate useful preclinical data is becoming an increasingly important question. We believe that our work bears significance for ongoing and future efforts to address this question in that it provides a collection of data that may help us better understand the data production capacity of microengineered tumor-on-a-chip systems and the potential value of biological data from such models for preclinical development and assessment of CAR T therapy for solid tumors.

Although building our system is not trivial and requires rather complex experimental work, we think that our methods and general workflow are translatable to other laboratories including those without engineering expertise. Specifically, the open-top device can be fabricated using simple soft lithographic techniques and made-to-order, custom-designed 3D printed molds obtained from commercial manufacturers without the need for capital equipment. Obviously, engineering the tumor constructs in this device is not an easy task and may indeed be perceived as a significant barrier to adopting our technology. Since our method is based on spontaneous organization of the starting cell populations into a vascularized tissue, our experience has been that efforts required to gain sufficient proficiency in the demonstrated 3D culture techniques are comparable to those needed to establish conventional organoid cultures for long-term experiments. What can be particularly challenging is to optimize experimental parameters for co-culture (e.g., cell ratio, seeding density, media formulation, hydrogel concentration), but as long as such optimized culture protocols can be made available (see **Methods**), we are optimistic about the likelihood of success in establishing the same or similar models in other laboratories.

While allowing us to demonstrate the advanced capabilities of our technology, the presented data also help us identify areas of further investigation. Among them is to refine and enhance the physiological realism of our microengineered system. In particular, it is questionable how closely the vascularized stroma of our model recapitulates its counterpart in vivo. In the human lung adenocarcinoma model described here, this compartment contains normal lung fibroblasts, whereas the native TME is interspersed with cancer-associated fibroblasts (CAFs) known to actively interact with cancer cells using soluble and insoluble cues to enhance their malignant phenotype^116–119^. Interestingly, our scRNA-seq analysis indicated that despite their origin from healthy lungs, the fibroblasts grown in our tumor models overexpress CD44 and TWIST1, which are expressed in higher abundance in CAFs, and use them to communicate with tumor cells as exemplified by an interacting ligand-receptor pair of SPP1-CD44 (**Supplementary Fig. 16, Extended Data Fig. 2c**), which has been shown to mediate the ability of the TME to promote cancer progression^120^. Our data also showed significant upregulation of TGFB1, HGF, and FGF7 in the fibroblast population adjacent to tumor explants (**Figs. 3h, 3m, Supplementary Fig. 16**). These observations raise the possibility that the normal fibroblasts in our system may undergo phenotypic changes during long-term culture with tumor explants to acquire certain features of CAFs^121^. Nevertheless, efforts should be made to incorporate CAFs in the stroma and examine i) how they affect the behavior of cancer cells and more importantly ii) whether and how their presence in the microengineered TME influences the trafficking and anti-tumor function of infused CAR T cells. Considering that cancer development and progression can alter the phenotype and functional capacity of endothelial cells^91,122^, similar studies are necessary to investigate whether and how CAR T-tumor interactions observed in the current study using normal endothelial cells change when the model contains blood vessels derived from tumor-associated endothelial cells.

Improving the biological complexity and physiological relevance of our model will also involve the inclusion of other cellular constituents of the native TME. For instance, tumor-associated macrophages (TAMs) are often pro-tumorigenic and have the ability to suppress anti-tumor activities of the immune system by producing immunosuppressive cytokines and growth factors^123^. It is possible that data generated by our current model without this tumor-protective population may overestimate therapeutic potential of CAR T cells against solid tumors. On a related note, the formation of tertiary lymphoid structures (TLSs), which are ectopically formed, organized aggregates of lymphocytes and antigen-presenting cells with specialized postcapillary blood vessels called high endothelial venules, has been described in several types of solid tumors^124–126^. By promoting lymphocyte trafficking, TLSs have been correlated with long-lasting anti-tumor immunity that can result in improved clinical outcomes in lung cancer patients^127,128^.

The ability to recreate these additional features of the native TME will likely increase the fidelity and predictive capacity of our model. Alternatively, it would also be interesting to compare data from our current model without these additional components or its modified versions containing select immune cell types to those generated by CAR T cell infusion into in vivo animal models that contain the same types of malignant tumors or tumor organoid transplants. Such comparison may provide a means to probe and better understand how the resident immune cell populations in solid tumors affect the behavior of CAR T cells, allowing us to identify specific cell types and their biological signals that suppress CAR T cell trafficking and activities. These types of studies may in turn provide useful information and guidance for emerging efforts to use antigen-directed cytotoxicity of CAR T cells to target and deplete TAMs and other types of resident immune cells in the TME as a strategy to delay tumor progression^129,130^.

Demonstrating the potential of the microengineered CAR T therapy model for applications in drug development is another key accomplishment of our work that warrants further investigation. Through scRNA-seq-based interrogation of intercellular interactions in this model, we identified DPP4 as a druggable target to promote meso-CAR T cell trafficking into mesothelin-expressing human lung tumor constructs. Our proof-of-principle experiments using a pharmacological inhibitor of DPP4 (LAF237) showed significantly increased trafficking and tumor infiltration of meso-CAR T cells and arrested tumor growth due to drug treatment, verifying the therapeutic potential of our proposed approach. Although IC_50_ of LAF237 is typically in the nanomolar range^101^, the demonstrated beneficial effects required a much higher concentration (1000 nM), which is consistent with previous reports of micromolar concentrations of LAF237 required for effective inhibition of DPP4^101,131^. This may be attributed to the short elimination half-life of LAF237 (2-3 hours)^131^ and our dosing regimen in which the drug is administered into the model only once a day. Better understanding this and other aspects of LAF237 treatment will be an important goal for future efforts to explore the potential of the proposed combinatorial therapeutic strategy.

On a related note, LAF237, also known as Vildagliptin, is an anti-hyperglycemic drug that increases the activity of incretin hormones by inhibiting DPP4^102,132^. Given that vildagliptin is already in clinical use for the treatment of type 2 diabetes, our results may provide a rationale for exploring the possibility of repurposing this drug as a new strategy to augment the efficacy of CAR T therapies for solid tumors. Obviously, further development of this therapeutic approach will require efforts to acquire more extensive efficacy and toxicity data and to investigate its translatability to other types of CAR T cells, adoptively transferred T cells, and solid tumors beyond meso-CAR T treatment of human lung adenocarcinoma tumors tested in this study. Considering that CXCR3-CXCL10/11-DPP4 signaling is conserved across many different types of cancer^94,133^, we have reason to think that this method may be applicable to other clinical conditions of solid malignancies.

We note that studies have shown increased lymphocyte recruitment into mouse melanoma and hepatocellular carcinoma due to DPP4 inhibition^134,135^. Based on the ligand-receptor interaction analysis of scRNA-seq data, our investigation of DPP4 was motivated and conducted independently of this previous work and provides new evidence verifying the efficacy of this principle in the context of treating malignant human solid tumors using human CAR T cells. Given the significant interspecies difference in the activity of T cells and DPP4^18,136^, we believe that the human-relevant data generated in our study add significant value to the existing knowledge. From a broader perspective, these results may contribute to ongoing efforts to address the rapidly increasing need for new strategies to augment and maximize the therapeutic efficacy of CAR T treatment for solid tumors^15,18,29,137^. Unlike CD19-targeted CAR T cells used for blood cancers, CAR T cells designed for treating solid tumors undergo limited homeostatic expansion after infusion due to the physical and molecular barriers created by the immunosuppressive TME that prevent their recognition of target antigens^138^. Clinical and animal studies have also shown the distribution of infused adoptively transferred T cells not only in tumor sites but also in lymphoid and other organs such as the bone marrow, lymph nodes, spleen, liver, and lungs^139^. These become problematic by limiting the number of tumor-infiltrating CAR T cells necessary for effective treatment^138^. Since increasing the dose of CAR T cells to boost anti-tumor activity can lead to lethal toxicities^138^, enhancing their ability to efficiently home to target tumors is emerging as one of the most promising approaches towards more efficacious and safer CAR T therapy of solid tumors^140,141^. Our work shows how these types of studies could be designed and carried out in vitro to identify new therapeutic targets and evaluate the effects of their pharmacological modulators.

As an extension of this study, we discovered a set of metabolites whose levels were correlated with the efficacy of LAF237 treatment during meso-CAR T therapy in our model. The significance of this work can be discussed in the context of metabolic alterations in the immunosuppressive tumor microenvironment and clinical challenges associated with monitoring the progression of CAR T therapy^142,143^. To address our limited understanding of tumor immune evasion by metabolic reprogramming, emerging studies have shed light on the molecular mechanisms of metabolic regulation of CAR T effector function and persistence by individual metabolites^144,145^. For example, studies have shown significant correlations between CAR T cell function in patients and fumarate hydratase levels in the microenvironment^144^, and that inosine can serve as an alternative carbon source for CAR T cell function when glucose supply is restricted^145^. These results illustrate not only the potency of metabolic regulation of immune functions but also the possibility of monitoring metabolite changes to infer CAR T cell performance in vivo. Although analysis of tumor biopsies provides accurate clinical measurement of the CAR T activities, the invasiveness and complexity of this procedure make it impractical for routine clinical monitoring^143,146^. Therefore, the identification of correlative and specific metabolite biomarkers may complement current methods for routine clinical assessment of therapy progression, including qPCR and flow cytometry of blood samples^147,148^, to provide additional value in metabolic measurement and monitoring of CAR T therapy and related treatment^149^. The differentially regulated metabolites identified in this study have not been described previously in the context of immunotherapy and may provide candidates for the development of clinical biomarkers for meso-CAR T therapy of lung tumors coupled with LAF237 treatment. Clinical translation of these types of findings will require in vivo validation, as well as further consideration of their allometric scaling and practical value in routine clinical settings. Our work focused on metabolomic analysis but it is not difficult to envision that the same approach would be applicable to probing lipidomic and proteomic profiles to diversify the development of predictive, specific biomarkers.

Further development of our work will also require more rigorous efforts to examine and validate the physiological relevance of our model and its findings. The in vitro-in vivo comparison of meso-CAR-CCR2 T cell activities presented in this paper provides a good example of how such studies can be conducted. This work allowed us to demonstrate the benefit of CCR2 for tumor-directed trafficking of meso-CAR T cells and verify this result by showing similar results in matching mouse experiments. This approach was also instrumental for showing delayed therapeutic effects of the armored CAR T cells. Given the known and suspected species-specific differences in immunological profiles of malignant tumors and their local environment^150^, however, efforts should be made to explore how data from human subject research can be made more readily available for model validation work, which will require alliances and coordinated efforts between clinical investigators and model developers. On a related note, these types of studies tend to focus on in vitro-in vivo similarities but considering that in vitro models as reductionist representations of their in vivo counterparts are constructed with select cell populations in simplified culture environments, the discrepancies between in vitro and in vivo results may be as valuable and informative for studying the role of specific cellular components or environmental factors in cancer-immune interactions in vivo.

Finally, the utility and potential of our technology are not limited to investigating CAR T therapy and could readily be extended to in vitro modeling and preclinical assessment of cancer immunotherapies using other types of cells, such as TCR T cells, CAR NK cells, and CAR macrophages^95,151–153^. Based on previous findings that tumor responses to immune checkpoint blockade (ICB) therapies require sufficient tumor-infiltrating lymphocytes^154,155^, increasing efforts are being made to develop more advanced treatment strategies that combine different modalities of cancer immunotherapy (e.g., CAR T-ICB therapy)^156^. Our platform may also provide a useful tool for these types of studies by enabling combinatorial screening of different cells/drug compounds to identify conditions that promote synergistic effects^32,154,157,158^.

## Methods

### Device fabrication and assembly

The open-top microdevice used for in vitro transplantation of tumor explants (**Fig. 1**, **Supplementary Fig. 1**) consists of three device layers of micropatterned poly(dimethylsiloxane) (PDMS), which were fabricated separately using conventional soft lithographic techniques. The casting molds necessary for PDMS-based replica molding of the culture chambers, open-top ceiling, and removable inserts of our device were produced by stereolithographic 3D printing (Protolabs, USA). For some experiments, the mold for the culture chamber was fabricated in a negative photoresist (SU-8 2100, MicroChem, USA) using photolithographic techniques to create positive relief structures on a prime-grade 4-inch silicon wafer (Wafer World Inc., USA) per manufacturer recommended protocols as described before^159^. To fabricate the micropatterned PDMS device layers, Sylgard 184 silicone elastomer base (Dow Corning, USA) and the curing agent were mixed at a ratio of 10:1 (w/w), cast over the photolithographically prepared or 3D printed molds, degassed under vacuum in a desiccating chamber (United Scientific Supplies, USA) for 30 minutes, and cured at 65°C for 2 hours. Cured PDMS layers were then cut and released from the molds and punched using 1-mm biopsy punches (Miltex, Integra, USA) to generate fluidic access ports. Holes were made through an additional blank PDMS slab and used as media reservoirs. Following fabrication of the device layers, the three-lane culture chamber was aligned and bonded to the open-top ceiling layer as shown in **Supplementary Fig. 1** using uncured PDMS as an adhesive material^159^. The assembled open-top microdevice array was baked at 65°C for 2 hours to fully cure the PDMS glue and was stored at room temperature until use. When necessary, device bonding was tested by microscopic inspection of the chambers filled with food coloring dye to detect fluid leakage between the bonded layers.

### Cell culture

Human non-small cell lung cancer cell line A549 cells (CCL-185, ATCC) were used as model tumor cells for studying meso-CAR T therapy. Since wild-type A549 cells only have minimal expression of mesothelin, lentiviral transduction of A549 cells was applied for stable expression of high levels of human mesothelin (**Supplementary Fig. 6**) as previously described^35,160^. The transduced cell line was referred to as meso-A549 in this study. A human mesothelioma cell line derived from a patient tumor, EMP (parental), was transduced with a lentivirus to stably express human mesothelin and thus named EMMeso as previously described^34^. The EMMeso tumor cells were used for the study of meso-CAR-CCR2 T cell therapy presented in **Figure 4**. Wild-type A549 cells, meso-A549 cells, and EMMeso cells were also transduced to stably express green fluorescent protein (GFP). A549 and EMMeso cells, primary normal human lung fibroblasts (hLF; CC-2512, Lonza), primary human umbilical vein endothelial cells (HUVEC; cAP-0001RFP and cAP-0001GFP, Angio-Proteomie), and primary human lung microvascular endothelial cells (HMVEC-L; CC-2527, Lonza) were cultured and maintained in RPMI 1640 media supplemented with 10% fetal bovine serum (FBS), hLF media supplemented with growth factors included in the FGM-2 BulletKit (CC-3132, Lonza), endothelial cell media supplemented with growth factors included in the EGM-2 BulletKit (CC-3162, Lonza), and endothelial cell media supplemented with growth factors included in the EGM-2 MV BulletKit (CC-3202, Lonza), respectively. These cells were used between passages 2 and 3 for all experiments.

### Tissue engineering of blood vessels

The assembled open-top microdevice array and additional device layers containing the removable inserts and media reservoirs were first sterilized by exposing to high-power ultraviolet (UV) light (Electro-lite ELC-500) for 30 minutes. Subsequently, the insert layer was fit into the wells of the open-top ceiling, after which the culture chamber was filled and incubated with 2 mg/ml (0.2% w/v in 10 mM Tris-HCl buffer, pH 8.5) of sterile dopamine hydrochloride solution at room temperature for 2 hours to form a surface coating of poly(dopamine) (PDA) on PDMS for enhanced adhesion to hydrogel scaffolds^159^, after which the solution was gently aspirated from the device. The PDA-coated microdevices were kept dry and sterile until use.

To construct a vascular bed for in vitro transplantation of solid tumors, 15 μl of fibrin gel pre-polymer solution was injected through the inlet access port into the middle lane of the culture chamber. The pre-polymer solution contained fibrinogen at a final concentration of 9 mg/ml (F8630, Sigma), thrombin at 1 U/ml (T7513, Sigma), aprotinin at 0.25 U/ml (A1153, Sigma), endothelial cells at 2.5-5 × 10^6^ cells/ml for both HUVECs and HMVEC-L, and fibroblasts at 4 × 10^6^ cells/ml. Injection of this mixture was performed manually in a biosafety cabinet using a 20-μl pipette within 5 seconds for each device to prevent unwanted premature gelation of fibrinogen. During this step, close attention was paid to the advancement of the liquid meniscus along the length of the middle lane. This method of visual inspection using the naked eye was useful for detecting unwanted fluid spillage into the side lanes and allowed us to verify the confinement of the mixture solution between the microfabricated rails. The key to success of this procedure was to apply and maintain constant, gentle thumb pressure to the pipette plunger during injection. The microdevice was then transferred to a cell culture incubator to induce fibrin gelation at 37°C for 10 minutes. Upon gelation, the complete endothelial cell growth media (EGM-2 or EGM-2 MV depending on the experiment) were introduced to the side channels of the culture chamber through media reservoirs. Next day, the side channels were filled with fresh EGM-2 or EGM-2 MV media containing a suspension of endothelial cells at 1 × 10^7^ cells/ml to form endothelial lining on the channel walls and the exposed surfaces of the hydrogel scaffold. The endothelial cell growth media in the reservoirs were changed every other day during the subsequent culture period. During culture, the fibrin construct was monitored daily using phase-contrast microscopy (and fluorescence microscopy when HUVECs were used) for visual confirmation of the development of interconnected tubular structures due to vasculogenesis, as well as for optical detection of any deleterious events, especially a loss of structural integrity of the hydrogel scaffold often caused by gel detachment from channel walls or spontaneous degradation. Our culture protocol was optimized to prevent such events while permitting sustained vascular development and long-term maintenance. When necessary, vascular perfusability was assessed by flowing media containing 1-µm fluorescence microbeads or 70-kDa FITC-dextran from one side channel to the other.

### In vivo xenograft tumor model and evaluation of CAR T therapy

All animal experiment protocols were approved and conducted in accordance with the Institutional Animal Care and Use Committee (IACUC) at the University of Pennsylvania. Our study used six-to ten-week-old female NOD/scid/IL2rγ^−/−^ (NSG) mice bred in the University of Pennsylvania Stem Cell and Xenograft Core as described^34,160^. The mice were housed under specific pathogen-free conditions in microisolator cages and given ad-libitum access to sterilized food and acidified water. An equal number of 1 × 10^6^ wild-type A549 cells and meso-A549 cells in PBS solution were injected into the left and right flanks of NSG mice, respectively. To generate EMMeso tumors in mice, a total of 2 × 10^6^ EMMeso tumor cells were subcutaneously injected in the flanks of NSG mice in PBS. For all experiments, tumor volume was monitored and measured over time using calipers for the entire duration of the experiments. Tumor volumes were calculated using the formula 0.5 × (length) × (width)^2^. After tumors were established (100-200 mm^3^ usually after 3-4 weeks), the mice carrying the tumors were sacrificed, and the xenograft tumors were harvested and precision-cut sliced using a Compresstome vibrating microtome (Precisionary Instruments LLC) with aseptic techniques. The sliced xenograft tumors were maintained in RPMI 1640 supplemented with 10% FBS at 37°C with 5% CO_2_ and were used for in vitro transplantation to the microengineered devices on the same day. The in vivo xenograft experiments were repeated more than eleven times in independent fashion.

For in vivo evaluation of CAR T therapy, mice were randomly assigned to one of the following three treatment groups after the formation of tumors: (i) 5 × 10^6^ non-transduced donor T cells, (ii) 5 × 10^6^ meso-CAR positive T cells, and (iii) 5 × 10^6^ meso-CAR positive T cells transduced with CCR2 lentivirus. Each group contained 3-10 mice. T cells were suspended in 100 µl of PBS and injected into the tail vein. The mice infused with T cells were sacrificed at Day 5 or Day 33 post-injection, and the tumors were harvested, micro-dissected, and digested in a solution of BD Horizon™ Dri Tumor & Tissue Dissociation Reagent (BD Biosciences) at 37°C for 30 minutes with frequent agitation. The digested tumors were then filtered through 70 μm nylon mesh cell strainers and washed twice in 1% BSA/DPBS/2 mM EDTA (without Ca++/Mg++), and red blood cells were lysed if needed (BD Pharm Lyse; BD Biosciences). Cells were strained again through a 30 μm nylon mesh cell strainer to obtain single-cell suspensions.

### Acquisition of human patient tumors

Operative tumor biopsy samples were obtained from non-small cell lung cancer (NSCLC) and malignant pleural mesothelioma (MPM) patients undergoing potentially curative surgery (n = 2) and pleurectomy (n = 3), respectively. The study was approved by the Institutional Review Board at the University of Pennsylvania and informed consent was obtained from all patients. The patient tumor explants were sliced and maintained as described above within 2 hours of resection, without freezing or overnight “rest”.

### Transplantation of tumor constructs

To prepare for in vitro transplantation, the sliced tumor explants from both patients and xenografts were manually minced into small pieces using razor blades (Hi-Stainless, Feather). The minced explant pieces were then filtered to select for diameters between 100 and 400 μm and suspended in complete endothelial cell growth media for the next step. To implant patient and xenograft tumor explant pieces, the inserts layer was removed first to expose the open-top wells (**Fig. 1**). A 10 μl of EGM-2 or EGM-2 MV media suspension containing around 10 tumor explant pieces was then added to the wells using a micropipette (Eppendorf). The injected tumor tissues were allowed to settle into the open-top wells overnight, after which a mixture of fibrinogen and thrombin at the same concentrations was added to seal the open-top wells. To examine the viability of patient tumor explants, we used Live/Dead™ Viability/Cytotoxicity Kit (L3224, ThermoFisher Scientific). The labeled cells in the tumor explants were examined using a laser scanning confocal microscope (LSM 800, Zeiss). To retrieve tumor tissues for compartment-specific analysis, we first cut around the acellular fibrin gel seal and gently removed it from the open-top wells using tweezers with sharp tips (Pixnor). The tumors in the wells were then separated from the surrounding stroma using fine-point tweezers, aspirated, and pooled together for subsequent analysis. The tumor compartment was defined as the original space of the open-top wells that later contained the tumor constructs and had the same diameter as the open-top wells (mostly 500 µm unless otherwise noted). The stroma compartment was defined as the whole of the gel construct excluding the tumor compartment.

### Isolation of tumor constructs and CAR T cells from culture chambers

The isolation of whole tumor constructs from culture chambers was made possible by our device bonding method described above. Specifically, when the device was assembled by using uncured PDMS as a glue, the PDMS layer comprising the open-top ceiling could be peeled apart from the underlying culture chamber by applying pulling force to one of its four corners and sustaining it until the entire layer was detached. This delamination was not possible when the layers were bonded using air or oxygen plasma commonly used for assembly of microfluidic devices, which results in irreversible bonding.

Prior to device delamination, the acellular fibrin gel seal was first carefully removed to expose the embedded tumor in the open-top wells. Subsequently, a manual procedure was performed to lift one of the sharp corners of the open-top layer and peel it gently and slowly (roughly 2-3 mm/s), during which great care was taken to ensure that the tumor construct remained attached to the microfabricated rails and the bottom channel surface without a loss of its structural integrity. We found that the speed at which the open-top layer was peeled apart was the key determinant of the success of this procedure – rushed delamination of the device led to physical damage of the tumor construct and in some cases, tearing of the PDMS layer. The average success rate of this procedure performed as described, which was assessed by histological examination of the shape and integrity of harvested tissues, was approximately 90% (**Supplementary Fig. 10**).

For tissue compartment-specific analysis shown in **Figures 3** and **5**, we used a different procedure for cell and tissue isolation. Briefly, tumors were exposed by removing the fibrin plug sealing the open-top wells, after which they were separated from the surrounding stroma using fine-point tweezers and gently aspirated from the wells for further analysis – what was retrieved at the end of this step was counted as the tumor compartment in our analysis. Next, the hydrogel construct containing CAR T cells and stromal cells remaining in the device, which was counted as the stromal compartment in our analysis, was infused with a tissue digestion buffer (TTDR, BD Horizon) and left in a cell culture incubator for 30 minutes to disintegrate the stromal tissue. When the tissue was digested into fragmented pieces, they were aspirated out of the device together with all of the liquid content. Subsequently, more digestion buffer was added to the tissue suspension up to a total volume of 10 ml, and the mixture was incubated for another 30 minutes to obtain single cell suspensions.

### Evaluation of vascular features and perfusability

Quantitative analysis of vascular development was conducted by measuring the percentage of vascular area, total branch length, number of junctions, and the average diameter of vessels in the self-assembled vascular network. Specifically, the AngioTool 0.5a plugin in ImageJ was used to identify vessels in a given image after applying a recursive Gaussian filter, followed by multiscale Hessian analysis and skeletonization. The vessel intensity parameter was adjusted to 15 or 20, with the background noise removed and other options in default settings, to make sure that the overlay accurately represented the vascular structures. The Vessel Analysis Tool and Mexican Hat Filter plugin were used for vessel diameter measurement. Images were converted to 8-bit grayscale in Fiji, then applied with the Mexican Hat Filter to enhance vessel structures and with the Geometry to Distance Map function (threshold = 255) to minimize noise. Ten random vessel segments were selected per image with the Diameter Measurements feature to calculate the average diameter of blood vessels.

To test the perfusability of the microengineered vascular network in our model, we used 70-kDa FITC-dextran (46945-100MG-F, Sigma-Aldrich) and fluorescently labeled 1-μm microbeads (FluoSpheres; F-8821 and F-8823, ThermoFisher) as flow tracers for visualization. To generate flow through the vasculature, the cell culture media in the media reservoirs were aspirated, after which either a FITC-dextran solution (50 μg/ml in PBS) or a suspension of microbeads (1:10,000 dilution in PBS) was injected into one of the side channels to generate pressure gradient across the vascularized hydrogel scaffold. Vascular perfusion was then visualized and monitored using a laser scanning confocal microscope (LSM 800, Zeiss) with a 10×/0.45 objective (C-Apochromat, water immersion, Zeiss). Time-lapse and Z-stack confocal images were acquired for 2 minutes and processed using the ZEN software (Zeiss).

### Generation and modification of meso-CAR T cells and on-chip evaluation of their activity

The plasmid design for meso-CAR and CCR2b, lentivirus packaging, as well as T cell activation, expansion, and cryopreservation have been previously described^160^. Briefly, the single-chain Fv domain of the anti-mesothelin antibody (scFv SS1), originally provided by Dr. Ira Pastan (National Cancer Institute/NIH, Bethesda, MD), was subcloned into the lentiviral vector pELNS that was driven by the EF1 α (eukaryotic translation elongation factor 1 alpha) promoter ^35^. All CAR constructs contained a CD8 hinge and transmembrane domain, 4-1BB costimulatory domain, CD3ζ signaling domain, and fluorescent reporter GFP or mCherry to evaluate transduction efficiency (i.e., SS1BBz mesoCAR as detailed in **Supplementary Fig. 6**). Primary human T cells were obtained from a total of eight healthy volunteer donors at the Human Immunology Core at the University of Pennsylvania following leukapheresis by negative selection using RosetteSep kits (Stem Cell Technologies). All specimens were collected under a University Institutional Review Board-approved protocol, and written informed consent was obtained from each donor. T cells were cultured in RPMI 1640 supplemented with 10% FCS (R10) and stimulated with magnetic microbeads coated with anti-CD3/anti-CD28 at a 1:3 cell to bead ratio without the addition of exogenous IL-2. Approximately 24 hours after activation, T cells were transduced with the lentiviral vectors encoding SS1BBz mesoCAR at a MOI of ∼5. T cells were counted and fed with complete RPMI media (R10) every 2 days and once appeared to become quiescent, as determined by both decreased growth kinetics and cell size, they were cryopreserved for long-term storage.

To model meso-CAR T therapy in our vascularized model, the cryopreserved T cell injection product that contained non-transduced or meso-CAR T cells (with or without CCR2 expression) were thawed and recovered overnight at a density of 1 × 10^6^ cells/cm^2^/ml in complete RPMI media (R10) prior to their infusion. Next day, the recovered T cells were first labelled with a fluorescent dye at a final concentration of 1 µM (CellTracker Deep Red, ThermoFisher) and resuspended in EGM-2 or EGM-2 MV media. The non-transduced and meso-CAR T cell suspension were then injected into the vascularized tumor models through one of the side microchannels at ∼150 μl of an equal cell density for each individual microdevice. The injected T cells were perfused through the vascular network and the flow of T cells was maintained overnight by continuous tilting on a rocker inside a cell culture incubator, resulting in a volumetric flow rate of 675.52 μl/min and an average wall shear stress of 6.59 dyn/cm^2^. Next day, the microdevices with infused T cells were washed with fresh EGM-2 or EGM-2 MV media to remove non-attached T cells. The remaining non-transduced or meso-CAR T cells were cultured for about 6 days. The microdevices with infused T cells were imaged every two days using a laser scanning confocal microscope (LSM 800, Zeiss) to monitor the dynamics of T cell trafficking and tumor growth. To capture specific events such as extravasation and infiltration into tumor masses, time-lapse and Z-stack confocal images were acquired for 25 to 45 minutes and processed using ZEN software (Zeiss). The time-series confocal images were processed in ImageJ (National Institutes of Health) and analyzed various endpoints described in the paper.

### Histology and immunofluorescence staining

The vascularized tumor tissues were individually washed with PBS and fixed in 10% normal buffered formalin in situ overnight at 4°C. The corresponding microdevices were individually delaminated by carefully peeling off the open-top ceiling layers. The fixed tumor constructs were carefully released from the culture chambers, transferred to pre-labeled tissue cassettes, and submerged in ethanol for dehydration and subsequent paraffin embedding. Thin tissue sections with a thickness of 5 μm were cut from the paraffin-embedded tissue blocks.

For staining with Hematoxylin and Eosin (H&E), the slides containing paraffin sections were deparaffinized and rehydrated by immersing the slides sequentially into 3× Xylene, 2× 100% ethanol, 95-90-80-70% ethanol, and distilled water. The slides were then immersed in 10 mM citric acid buffer (pH 6.0) and incubated in a microwave oven for 15 minutes. After gentle rinse, the slides with tissue sections were blocked with a protein blocking agent and immersed in Hematoxylin followed by rinsing with deionized water. The slides were further immersed in Eosin for 30 seconds and dehydrated in 95% ethanol-100% ethanol-xylene solutions. Stained tissue sections were covered with coverslip slides using Permount^TM^ and stored until imaging.

For immunofluorescence staining, paraffin on the tissue section slides was cleared with xylene, and the slides were rehydrated through descending concentrations of ethanol. The slides were then treated with 3% H_2_O_2_/methanol for 30 minutes, followed by pretreatment with Antigen Unmasking solution (Vector Labs H3300) in a pressure cooker (Biocare Medical). After cooling, the slides were blocked in Sudan Black (199664-25G, Sigma-Aldrich) for 30 minutes at RT, rinsed in 0.1M Tris Buffer, and blocked with 2% fetal bovine serum for 15 minutes. After removing the block solution, the slides were incubated overnight at 4°C with CD8 antibody (RB-9009-PO, Thermo) at a 1:500 dilution, rinsed, and then incubated with anti-rabbit polymer secondary prediluted (K4003, DAKO) for 30 minutes at RT. Following rinsing, the slides were incubated with the TSA biotin complex (NEL7490B001, Perkin Elmer) at 1:50 for 10 minutes at RT and incubated with Alexa 488 Streptavidin secondary (A21370, Life Technologies) at 1:200 dilution for 30 minutes at RT. After rinsing, the slides were treated in preheated 5% SDS (CS-5585-28, Denville Scientific) for 7 minutes at 55°C, rinsed, and blocked again with 2% fetal bovine serum before incubating with CD31 antibody (M0828, Dako) at 1:200 dilution for 1 hour at RT. After rinsing, the slides were incubated with Alexa Fluor 594 goat anti-mouse secondary antibody (A11032, Invitrogen) for 30 minutes, rinsed, and incubated with Cytokeratin antibody (Z0662, Dako) at 1:1000 dilution for 1 hour at RT. Subsequently, the slides were rinsed, incubated with Alexa Fluor 647 goat anti-rabbit secondary antibody (A21245, Invitrogen) for 30 minutes, counterstained with DAPI, and rinsed again before coverslipping with Prolong Gold (P36930, Life Technologies). After drying, the slides were scanned at 20X using an Aperio IF slide scanner (Leica Biosystems) and imaged at 60X using a laser scanning confocal microscope (LSM800, Zeiss).

### Flow cytometry

To evaluate the expression of CAR T cell surface markers by flow cytometry, the tumor constructs were carefully separated from the stroma as described above and then removed and pooled together for subsequent processing. The sample suspension was digested using a tumor digestion buffer (TTDR, BD Horizon) according to the manufacturer’s instructions and incubated on a rocker in a cell culture incubator for one hour with vigorous pipetting every 10 minutes. The same digestion buffer was added to the microdevices without the tumor constructs to digest the stroma gels, which were separately collected and pooled together to obtain single cell suspensions. All cell suspensions were passed through 70 µm nylon mesh cell strainers to remove undigested clusters and multiplets.

All samples were stained and analyzed using standard flow cytometry. Single-cell suspensions from all samples were stained with Live/Dead cell stain (Invitrogen) for 10 minutes at 4°C. For staining of human cell surface marker proteins, the cells were incubated at 4°C for 30 minutes in staining buffer (2% FBS in PBS) with the following fluorescently-labeled antibodies based on the manufacturer’s recommendations. The antibodies used include CD3 (clone HIT3a, 300310), CD45RO (clone UCHL1, 304246), CD103 (clone Ber-ACT8, 350221), PD-1 (clone EH12.2H7, 329924), CD62L (clone DREG-56, 304830), CD69 (clone FN50, 310926) purchased from Biolegend; CD8 (clone RPA-T8, 563795) purchased from BD Horizon. For experiments conducted to evaluate CAR T therapy using the in vivo EMMeso tumor model, 1 × 10^6^ cells from the digested single-cell suspensions were placed in standard fluorescence-activated cell sorting (FACS) tubes and stained for human CD45-BV421 (clone HI30) and/or CD3-FITC

(clone HIT3a) or CD3-PE/Cy7 (clone UCHT1) antibodies (all from Biolegend). Labeled cells were washed and resuspended in FACS buffer for flow cytometric analysis using LSRFortessa Cell Analyzer (BD Biosciences). Subsequent computer analysis was done using FlowJo software (v10.2). Negative gating was based on a “fluorescence minus one” (FMO) strategy.

### Single-cell reverse transcription, library preparation, and sequencing

To prepare for single-cell RNA sequencing analysis, 6 to 12 tissue samples from each array microdevice were pooled together. Similar to the preparation for flow cytometry, the tumor explants were carefully separated from the stroma and removed from the culture chambers. Single cell suspensions were obtained using the same tissue digestion buffer used above (TTDR, BD Horizon). The isolated cells were filtered, manually counted, and prepared at the desired cell concentration of 800-1000 cells/µl. Finally, 20,000 cells were collected and washed with PBS, and then loaded onto a Chromium Single Cell Chip (10x Genomics) according to the manufacturer’s instructions for co-encapsulation with barcoded beads at a target capture rate of 10,000 individual cells per sample.

The capture of RNA transcripts in the barcoded cells and reverse transcription of cDNA were performed using the manufacturer’s standard protocols. The Agilent TapeStation High Sensitivity D5000 ScreenTape was used for QC of generated cDNA. The barcoded cDNA was converted into single-cell RNA-seq libraries for sequencing using the Chromium Single Cell 3’ Reagent Kit v3 (10x Genomics) according to the manufacturer’s instructions. The Agilent TapeStation High Sensitivity D1000 ScreenTape was used for QC of prepared libraries for sizing (bp) and concentration. The final libraries from four samples (i.e., control tumor, control stroma, meso-tumor, and meso-stroma) were pooled together and sequenced by Illumina NovaSeq sequencer with a SP v1.5 FlowCell (100 cycles). The pooled libraries were sequenced to a target depth of at least 20,000 mean reads per cell.

### Computational analysis of single-cell RNA sequencing data

#### Data pre-processing and quality control

The raw FASTQ files containing sequence reads were pre-processed in the CellRanger pipeline (10X Genomics, v3.1.0). Reads produced from the gene expression profiling were aligned to the human GRCh38 reference genome, which was additionally customized to include the reference sequences of GFP that tagged the tumor cells and of RFP and SS1 scFv that were contained in the mesoCAR construct. On a side note, data showed negligible abundance of murine cells, which were excluded from our analysis. The feature-barcode matrices were generated using default parameters for each sample, which record the number of unique molecular identifiers for each gene within each cell barcode. The quality of cells was assessed based on the proportion of mitochondrial gene counts and the number of genes detected per cell. Low-quality cells were filtered out if the proportion of mitochondrial gene counts was higher than 20%. In addition, only cells with number of features/genes larger than 200 but less than 5000 were kept for subsequent analysis to avoid capture of possible doublet or multiplet.

#### Single-cell data integration and clustering analysis

The R package Seurat (v3.2.0)^161^ was used to filter out low-quality cells, normalize the raw counts data to account for sequencing depth, scale and identify highly variable features, cluster upon dimensionality reduction, and integrate samples. Briefly, the gene-count matrix was first normalized by cell-specific size factor, log-transformed, and then scaled to unit variance and zero mean. Samples from the same experiment were integrated using the merge function for cell type identification and direct comparisons. Principal components were calculated using the data decomposition technique of Latent Semantic Indexing (LSI) and used to reduce the dataset into two dimensions. Unsupervised hierarchical clustering of cells was generated and visualized using Uniform Manifold Approximation and Projection (UMAP) in Seurat. Differential-expression tests for all cells were performed using FindAllMarkers function. Genes with log_2_-fold changes >0.25, expression in at least 25% of cells in tested groups, and adjusted p <0.05 were regarded as significantly differentially expressed genes (DEGs). Clusters were labelled based on the expression of the top DEGs as well as the canonical markers for each cell type.

#### Gene Ontology (GO) analysis

For each cluster, GO analysis was conducted by comparing top 100 genes that were highly differentially expressed and all the other genes in the dataset. GO analysis was performed using PANTHER Overrepresentation test (Released 20200728) (PANTHER version 16.0 Released 2020-12-01) with default parameters (Fisher’s exact test; cut off at False Discovery Rate p < 0.05). The results were visualized with heatmap using Morpheus (https://software.broadinstitute.org/morpheus).

#### Comparison of CAR T cell clusters with external human single-cell gene signatures

To assess the physiological relevance of CAR T cell phenotypes observed in our microengineered tumor constructs, we compared our CAR T cell cluster annotations against a published human single-cell RNA-seq dataset containing T cells isolated from tumors, adjacent normal tissues, and peripheral blood of lung cancer patients^78^. The single-cell profiles of 12,346 T cells from 14 patients were obtained under GEO accession number GSE99254. The R package SingleR (v1.4.1)^162^ was used for comparison following standard procedures. The raw count matrices were downloaded from the publication and then normalized and clustered in Seurat following standard workflow. The same cell annotations from the reference publication were maintained on a cell name/barcode basis. Note that both the reference and our subject dataset were subsetted for CD4 and CD8 T cells, which were separately compared. SingleR calculates correlations between subject and reference cells using variable genes in the reference dataset to assign cellular identity to each cell in our subject dataset. The SingleR comparison outputs an UMAP clustering of our CAR T cells with the predicted T cell annotation from the reference dataset, and a heatmap of prediction scores of each cell within the subject dataset towards the corresponding cell type from the reference dataset.

#### Single-cell trajectory analysis

Monocle 3 (v1.0.0)^163,164^ was used for the analysis of pseudotime transition of cell transcriptional states. In this analysis, cells were ordered in pseudotime as a measure of progress through biological processes based on their transcriptional similarities. The aggregated Seurat object after dimensional reduction as constructed above was converted to a cell_data_set object in Monocle. Single cells of each cell type were visualized in the UMAP space with similar clustering to that in Seurat. CAR T cell clusters and tumor cell clusters were subsetted for pseudotime trajectory analysis, separately. The principal graphs were generated along each trajectory to represent possible paths that cells can take in response to emerging cues in the tumor microenvironment. Cell-wise pseudotime was calculated based on its position along the principal graphs after manual selection of root-node for each trajectory. The root nodes for CAR T cell trajectory were selected from principal points with highest expression of T cell naiveness markers (i.e., SELL, TCF7, and LEF1). For tumor cell trajectories, root nodes were assigned to the earliest principal points for pseudotime computation. Moran’s I test was performed to identify genes with expression that are trajectory-dependent using the graph_test function. The smoothed gene marker expression along pseudotime was generated by the plot_genes_in_pseudotime function with a natural spline used to fit the gene expression along pseudotime. Top ranked genes with monotone increase or decrease of expression and biphasic trend of expression along the pseudotime were selected for comparison.

#### Analysis of inter-lineage interactions

We used CellPhoneDB 2.0^84,165^ to identify enriched ligand-receptor interactions from our scRNA-seq datasets involved in soluble factor- or direct binding-mediated signaling of inter-lineage interactions in our microengineered tumor tissues. CellPhoneDB is a manually curated public repository of ligands, receptors and their interactions, integrated with a statistical framework for inferring cell-cell communication networks from single-cell transcriptomic data. The normalized count matrix with metadata column consisting of annotations of cell identity and other default parameters were provided as input. The interaction analysis was limited to the control tumor and meso-tumor models, separately. We estimated the potential interaction between two cell types mediated by a specific ligand-receptor pair by determining the mean of the average expression of the receptor in one cell type and the interacting ligand in the other cell type. To examine the statistical significance of such estimated interaction, random permutations were applied on the cell type cluster labels of individual cells for 1000 times. The p value of the likelihood of cell type specificity was estimated by the number of permutations that had interaction mean larger than the real mean value. The cutoff was set with the mean expression greater than 0.1 and p value smaller than 0.05. To precisely identify the specific interactions only between cells, the output matrix of significantly interacting ligand-receptor pairs was filtered to remove all direct integrin-ECM interactions. From those remaining, we prepared chord diagrams of interactions between each pair of interacting cell types and generated an overarching web using the Circlize package in R^166^. In the overarching web, each chord represents the bundle of all significant interactions between a particular pair of cell types. The chord width represents the total number of ligand-receptor pairs in each pair of interacting cell types. For closer assessment, all the directional interactions between ligand-receptor pairs within each interacting cell pair were also visualized in chord diagrams. The ligand-receptor pairs with total mean < 0.35 were neglected for visualization. In the directional chord diagrams, each chord represents the maximum of all significant means from all paired clusters for each interacting gene pair within each pair of interacting cell types. In this case, the chord width indicates the level of total mean of interaction between each specific ligand-receptor pair.

### Assessment of DPP4 activity

The DPP4 inhibitor - LAF237 (Vildagliptin, PharmaForm LLC) - was used to inhibit the enzymatic activity of DPP4. LAF237 was used at final concentrations of 50 nM and 1,000 nM. The DPP4 inhibitor was diluted in EGM-2 media to the target final concentrations and administered to the tumor-chip after the infusion of meso-CAR T cells. Fresh EGM-2 media with different concentrations of DPP4 inhibitors were separately prepared and changed daily until the end of culture. For evaluation of DPP4 activity, the spent cell culture media from individual microdevices were collected daily prior to routine media change. DPP4 activity was measured with the DPPIV-Glo Protease Assay (Promega) per manufacturer’s instructions.

### Inhibition of CXCR3

The anti-human CXCR3 antibody (clone 49801, MAB160, R&D Systems) was used to inhibit the CXCL10/11-CXCR3 signaling pathway in our model. The anti-hCXCR3 and its IgG_1_ isotype control antibody (clone 11711, MAB002, R&D Systems) were used at final concentrations of 10 µg/mL. The CAR T cells suspended in EGM-2 MV media were incubated with the anti-hCXCR3 and isotype control antibodies, as well as LAF237, at final concentrations for one hour at room temperature prior to initial infusion to the microdevices. Fresh EGM-2 MV media with final concentrations of anti-hCXCR3, isotype control, and LAF237 were separately prepared and changed daily until the end of culture.

### Pharmacological modulation of CD38-PECAM1 and LTB-LTBR

The human CD38 inhibitor - Daratumumab (HY-P9915, MedChemExpress) and the lymphotoxin β receptor IgG fusion protein - Baminercept (HY-P99459, MedChemExpress) were used to inhibit the interactions between CD38 and its ligand PECAM1 and between lymphotoxin β and its receptor, respectively. Daratumumab and Baminercept were used at final concentrations of 0.5 µg/ml and 10 µg/ml, respectively. Their human IgG_1_ isotype control antibody (HY-P99001, MedChemExpress) was used at a final concentration of 10 µg/ml. Daratumumab, Baminercept, and isotype control antibody were diluted in endothelial media to the target final concentrations and administered to the devices with meso-CAR T cells. Fresh endothelial media with different concentrations of Daratumumab, Baminercept, and isotype control antibody were separately prepared and changed daily until the end of culture.

### ELISA, LDH, and chemokine assays

To analyze and quantify the secretion of IL-2 and IFNγ from the activated T cells, the cell culture media from individual microdevices were collected at specified time points. Human IL-2 DuoSet ELISA kit (DY202-05, R&D Systems) and human IFN-gamma DuoSet ELISA kit (DY285B-05, R&D Systems) were used to measure the concentrations of IL-2 and IFNγ, respectively. Each assay was performed and analyzed following the manufacturer’s instructions. Briefly, 100 μl of cell culture samples or standards was added per well upon plate preparation and incubated for 2 hours at RT. Subsequently, the wells were washed four times with 200 μl of manufacturer-provided wash buffer and incubated with 100 μl of respective detection antibodies for another 2 hours at RT. After washing, 100 μl of Streptavidin-HRP was added per well and incubated for 20 minutes at RT in the dark. After washing, 100 μl of substrate solution was added per well and incubated for another 20 minutes at RT in the dark. Finally, 50 μl of stop solution was added per well, and the plate was measured for optical densities of each well using the absorbance mode of a microplate reader (Infinite M200, Tecan).

For measurement of CXCL10 and CXCL11 dynamics in the engineered tumor constructs, the cell culture media from individual microdevices were collected at specified time points and analyzed using human CXCL10/IP-10 Quantikine ELISA kit (DIP100, R&D Systems) and human CXCL11/I-TAC Quantikine ELISA kit (DCX110, R&D Systems) following the manufacturer’s instructions. Briefly, 100 μl of cell culture samples or standards in respective assay diluent was added per well and incubated for 2 hours at RT. The wells were washed four times with 200 μl of manufacturer-provided wash buffer and incubated with 200 μl of respective conjugates for another 2 hours at RT. After washing, 200 μl of substrate solution was added per well and incubated for 30 minutes at RT in the dark. Finally, 50 μl of stop solution was added per well, and the plate was measured in a microplate reader (Infinite M200, Tecan).

To analyze and quantify the cytotoxicity of CAR T cells, the cell culture media from at least three individual microdevices per group were collected on day 5 post infusion of CAR T cells and analyzed using a lactate dehydrogenase (LDH) assay kit (ab102526, abcam) following manufacturer’s instructions. 50 μl of cell culture samples or standards diluted in assay buffer and another 50 μl of reaction mix were added per well in sequence and mixed thoroughly. The output optical densities were measured immediately using a microplate reader (Infinite M200, Tecan) in a kinetic mode, every 3 minutes, for a total of 1 hour.

To measure chemokines, cell culture media from individual microdevices were collected one day prior to T cell infusion and analyzed using the human chemokine array G1 kit (AAH-CHE-G1-8, Ray Biotech) following the manufacturer’s instructions. Samples were analyzed without dilution, and the slides were imaged with a laser scanning confocal microscope (LSM 800, Zeiss).

### Measurement of intact and truncated forms of CXCL10

#### Preparation of aptamer-modified gold nanoparticle

Colloidal gold nanoparticles (GNP) capped with citrate were synthesized by seed-mediated growth according to the previous methods^167^. In brief, gold nanoparticles were grown from citrate-capped seed particles. The seed particles were synthesized by reacting 100 ml of 0.01 wt % HAuCl4 solution and 3 ml of 1 wt % citrate solution at 99 °C for 30 minutes. 100 ml of 0.01 wt % HAuCl4 solution was treated with 4 ml of seed solution and 400 μl of 1 wt % citrate solution at 99 °C for 30 minutes. An aptamer that specifically binds to valine was obtained from BIONEER (South Korea) and modified with a thiol group to bind onto the surface of gold nanoparticles. For the preparation of the GNP-aptamer complex, 10 ml of aqueous solution including ca. 14 × 10^13^ aptamers was mixed with 990 ml of gold nanoparticles solution (ca. 9 × 10^10^ nanoparticles/ml), and the mixture was kept at 4 °C for 24 hours.

#### Absorbance and Raman measurements of GNP-aptamer after mixing with samples

The absorbance spectra of the GNP-aptamer complex were measured by Cary 5000 UV-Vis-NIR spectrophotometer after mixing with filtered samples collected from device effluent. For Raman measurements, the GNP-aptamer complex was collected by centrifugation (4000 rpm, 10 minutes) after mixing. Next, the collated GNP-aptamer was dropped onto a silicon wafer and dried on the surface. Raman spectra were measured by using a micro-Raman system with a spectrometer SR-303iA (Andor Technology), 785 nm laser module I0785SR0100B (Innovative Photonic Solution Inc.), and Olympus BX-53 M TRF microscope (Olympus). The integration time was set to 5 s.

#### Immuno-precipitation

Immunoprecipitation of CXCL10 from device effluent samples was performed following the manufacturer’s protocol. Briefly, 50 μl of Dynabeads™ Protein G magnetic beads (10007D, Thermo Fisher) were placed on a magnet to separate the beads from the solution. After removing supernatant, the beads were incubated with 200 μl of rabbit anti-CXCL10 monoclonal antibody (MA5-32674, Invitrogen) solution at a concentration of 20 μg/ml for 1 hour and washed on the magnet to remove unbound antibodies. Then, 550 μl of the collected samples was added to the bead-antibody complex and incubated for 2 hours at room temperature to ensure antigen binding. To elute the target antigen from the beads, the complex was incubated in 20 μl of elution buffer for 2 min and separated from the magnet. The pH of collected eluates was adjusted by adding 1 M Tris-HCl pH 7.5 solution for Western blotting analysis. To generate the truncated CXCL10 solution as a control, recombinant human CXCL10 (266-IP, R&D systems) was incubated at a final concentration of 1 μg/ml with recombinant DPP4 (D3446, Sigma) at a final concentration of 2 U/ml in a 100 mM Tris-HCL pH 8 solution for 2 h at 37°C.

#### Western blotting

Western blotting was performed according to a previously established protocol with modifications^168^. Briefly, 25 μl of protein samples were resolved by 20% SDS-PAGE and transferred onto polyvinylidene fluoride membranes. The membranes were blocked with 5% skim milk for 1 hour at room temperature, followed by overnight incubation at 4°C with primary antibodies against CXCL10 (MAB266-100, R&D systems). After three washes, the membranes were incubated with horseradish peroxidase-conjugated secondary antibodies for 1 hour at room temperature. Protein bands were visualized using ECL Western Blotting Substrate (Pierce) and quantified using Fiji/ImageJ software.

### Metabolomics analysis

#### Supernate metabolite extraction

The spent media samples were collected from individual microdevices at each time point and individually frozen for storage at -80°C. A 5 μl of media was added to 120 μl of 25:25:10 (v/v/v) acetonitrile:methanol:water solution at −20°C, vortexed for 10 seconds, and put on ice for at least 5 minutes. The resulting extract was centrifuged at 16,000 × g for 20 minutes at 4°C, and the supernatant was transferred to tubes for liquid chromatography-mass spectrometry (LC-MS) analysis. A procedure blank was generated identically without spent media, which was used later to remove background ions.

#### Metabolite measurement by LC-MS

Metabolites were analyzed using a Vanquish Horizon UHPLC System (Thermo Scientific) coupled to an Orbitrap Exploris 480 Mass Spectrometer (Thermo Scientific). Waters XBridge BEH Amide XP Column (particle size, 2.5 μm; 150 mm (length) × 2.1 mm (i.d.)) was used for hydrophilic interaction chromatography (HILIC) separation. Column temperature was kept at 25 °C. Mobile phases A = 20 mM ammonium acetate and 22.5 mM ammonium hydroxide in 95:5 (v/v) water:acetonitrile (pH 9.45) and B = 100% acetonitrile were used for both ESI positive and negative modes. The linear gradient eluted from 90% B (0.0–2.0 min), 90% B to 75% B (2.0–3.0 min), 75% B (3.0–7.0 min), 75% B to 70% B (7.0–8.0 min), 70% B (8.0–9.0 min), 70% B to 50% B (9.0–10.0 min), 50% B (10.0–12.0 min), 50% B to 25% B (12.0–13.0 min), 25% B (13.0–14.0 min), 25% B to 0.5% B (14.0–16.0 min), 0.5% B (16.0–20.5 min), then stayed at 90% B for 4.5 min. The flow rate was used at 0.15 mL/min. The sample injection volume was 5 μL. ESI source parameters were set as follows: spray voltage, 3200 V or −2800 V, in positive or negative modes, respectively; sheath gas, 35 arb; aux gas, 10 arb; sweep gas, 0.5 arb; ion transfer tube temperature, 300 °C; vaporizer temperature, 35 °C. LC–MS data acquisition was operated under full scan polarity switching mode for all samples. The full scan was set as: orbitrap resolution, 120,000 at m/z 200; AGC target, 1e7; maximum injection time, 200 ms; scan range, 60–1000 m/z.

#### Data analysis

LC-MS raw data files (.raw) were converted to mzXML format using ProteoWizard (version 3.0.20315) ^169^. El-MAVEN (version 0.12.0) was used to generate a peak table containing m/z, retention time, and intensity for the peaks ^170^. Default parameters were used for peak picking except for the following: mass domain resolution, 5 ppm; time domain resolution, 10 scans; minimum intensity, 10,000; and minimum peak width, 5 scans. The resulting peak table was exported as a .csv file. Peak annotation of untargeted metabolomics data was performed using NetID with default parameters ^170^. Statistical analyses were performed using MetaboAnalyst 5.0 ^171^. Cut-off for significant change was set to FDR < 0.05.

### Immunofluorescence analysis

For in situ immunofluorescence staining, cell-containing gel constructs in our microdevices were washed with PBS, fixed with 4% paraformaldehyde (Electron Microscopy Sciences) for 15 minutes at RT. The fixed cells were then permeabilized with 0.25% Triton X-100 in PBS for 15 minutes and blocked with 3% bovine serum albumin in PBS (Sigma Aldrich) for 1 hour at RT. After washing, the cells were incubated overnight at 4°C with primary antibodies against CD3 for T lymphocytes (pre-diluted, rabbit monoclonal 2GV6, 790-4341/05278422001, Roche), mesothelin for target antigen (1:50, mouse monoclonal G-1, sc-271540, Santa Cruz Biotechnology), cleaved caspase-3 for apoptotic cells (1:200, rabbit polyclonal Asp175, 9661S, Cell Signaling Technology), pan-cytokeratin for tumor cells (1:250, mouse monoclonal AE1/AE3+5D3, ab86734, Abcam), CD31 for endothelial cells (1:200, mouse monoclonal JC/70A, ab9498 or ab215911, Abcam), and Phalloidin for cellular actin filaments (1:100, A22287, ThermoFisher Scientific). After washing, the cells were then incubated with secondary antibodies (Goat anti-Rabbit IgG H&L (Alexa Fluor 488), 1:500, A32731, ThermoFisher Scientific; Goat anti-Mouse IgG H&L (Alexa Fluor 488), 1:500, A32723, ThermoFisher Scientific; Goat anti-Mouse IgG H&L (Alexa Fluor Plus 555), 1:500, A32727, ThermoFisher Scientific; Goat anti-Rabbit IgG H&L (Alexa Fluor Plus 555), 1:500, A32732, ThermoFisher Scientific; Goat anti-Mouse IgG H&L (Alexa Fluor 647), 1:500, A21236, ThermoFisher Scientific) overnight at 4°C or for 2 hours at RT. Finally, cell nuclei were counterstained with DAPI (D1306, ThermoFisher Scientific).

Fluorescence images of the cells were acquired using a laser scanning confocal microscope (LSM 800, Zeiss) and processed using ZEN software (Zeiss) and ImageJ (National Institutes of Health). 3D reconstruction of imaged vasculature was performed using Imaris (Bitplane). Quantification of vascular features was achieved using AngioTool2 and Vessel Analysis plugins in ImageJ.

### Statistical analysis

Sample size for each experiment was determined on the basis of a minimum of *n* = 3 independent microdevices for each experimental group. Data were analyzed with Student’s *t*-test and with one-way and two-way ANOVA followed by appropriate post-hoc testing for multigroup pairwise comparisons in GraphPad Prism v8.2, unless noted otherwise. Results were presented as mean ± standard error of the mean (SEM). Statistical significance of the analyzed data was attributed to values of *\*P* < 0.05, \*\**P* < 0.01, and \*\*\**P* < 0.001 as determined by respective analyses.

---

**Extended Data Fig. 1.**
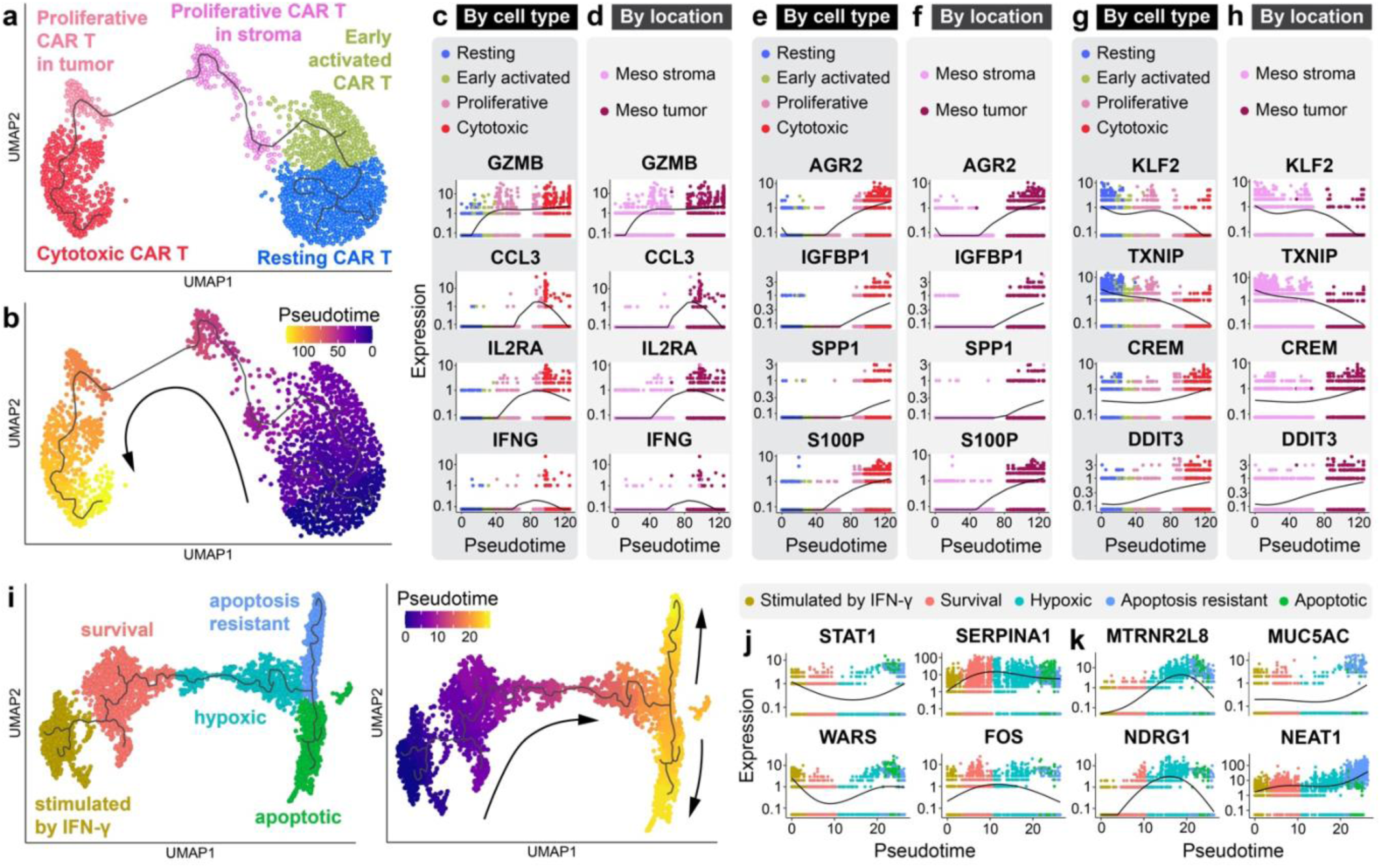
Single-cell trajectory analysis of CAR T and tumor cells in the meso-tumor model. **a,** UMAP plot showing CAR T cell subpopulations in the meso-tumor model re-clustered for trajectory analysis. **b,** Transition of CAR T cell phenotype along the pseudotime trajectory. **c-h**, Pseudotime dynamics of key marker expression plotted by cell type (**c,e,g**) and location (**d,f,h**). **c,d,** Biphasic regulation of gene markers associated with T cell activation (e.g., GZMB, CCL3, IL2RA, IFNG). These markers begin to increase with the emergence of early activated CAR T cells in the stroma and peak when the cells exhibit the phenotype of proliferative effector T cells within the tumors, which is followed by a plateau or gradual decrease with the progression of the cytotoxic activities of the CAR T cells while they are still in the tumors. **e,f,** The activated effector CAR T cells in the tumors show monotonically increasing expression of stress markers, such as AGR2, IGFBP1, S100P, and SPP1, demonstrating exposure and responses to increasing cellular stress in the CAR T cells as they gain effector function, enter the tumor environment, and engage cancer cells. **g,h,** Among transcription factors correlated with the phenotypic changes of CAR T cells are KLF2 and TXNIP. These markers exhibit almost monotonically decreasing expression, suggesting that their downregulation may be required for antigenic activation of CAR T cells due to their established roles in regulating T cell effector functions and glucose consumption^1,2^. Increased expression of CREM and DDIT3 is closely associated with the transition of CAR T cells towards effector lineages, consistent with their reported role as negative regulators of effector functions^3,4^. **i,** UMAP plots of tumor cell clusters in the meso-tumor model (left) and pseudotime trajectories (right). **j,k,** Dynamic expression of select markers by tumor cells over pseudotime. At the beginning of the pseudotime trajectory, the cells show the highest expression of genes activated by IFN-ɣ (e.g., STAT1, WARS), verifying their phenotype as tumor cells stimulated by effector CAR T cell-produced IFN-ɣ. These genes show a cycle of downregulation and subsequent recovery, which is reversed in SERPINA1, FOS, and other transcription factors associated with survival promotion. Another notable feature is the induction of genes implicated in tumor cell apoptosis or inhibition of cancer progression (e.g., MTRNR2L8, NDRG1)^5,6^ and their continuous increase prior to entering the apoptotic states, illustrating persistent deleterious effects of tumor-infiltrating CAR T cells. Transcriptomic signatures of tumor cells also include markers that promote tumor development and progression (e.g., MUC5AC, NEAT1), which may be interpreted as a mechanism to protect/recover stressed and apoptotic tumor cells affected by CAR T cells. Increased induction of these genes is indeed visible at the late stages of phenotypic transition as the tumor cells move towards the apoptotic state.

**Extended Data Fig. 2.**
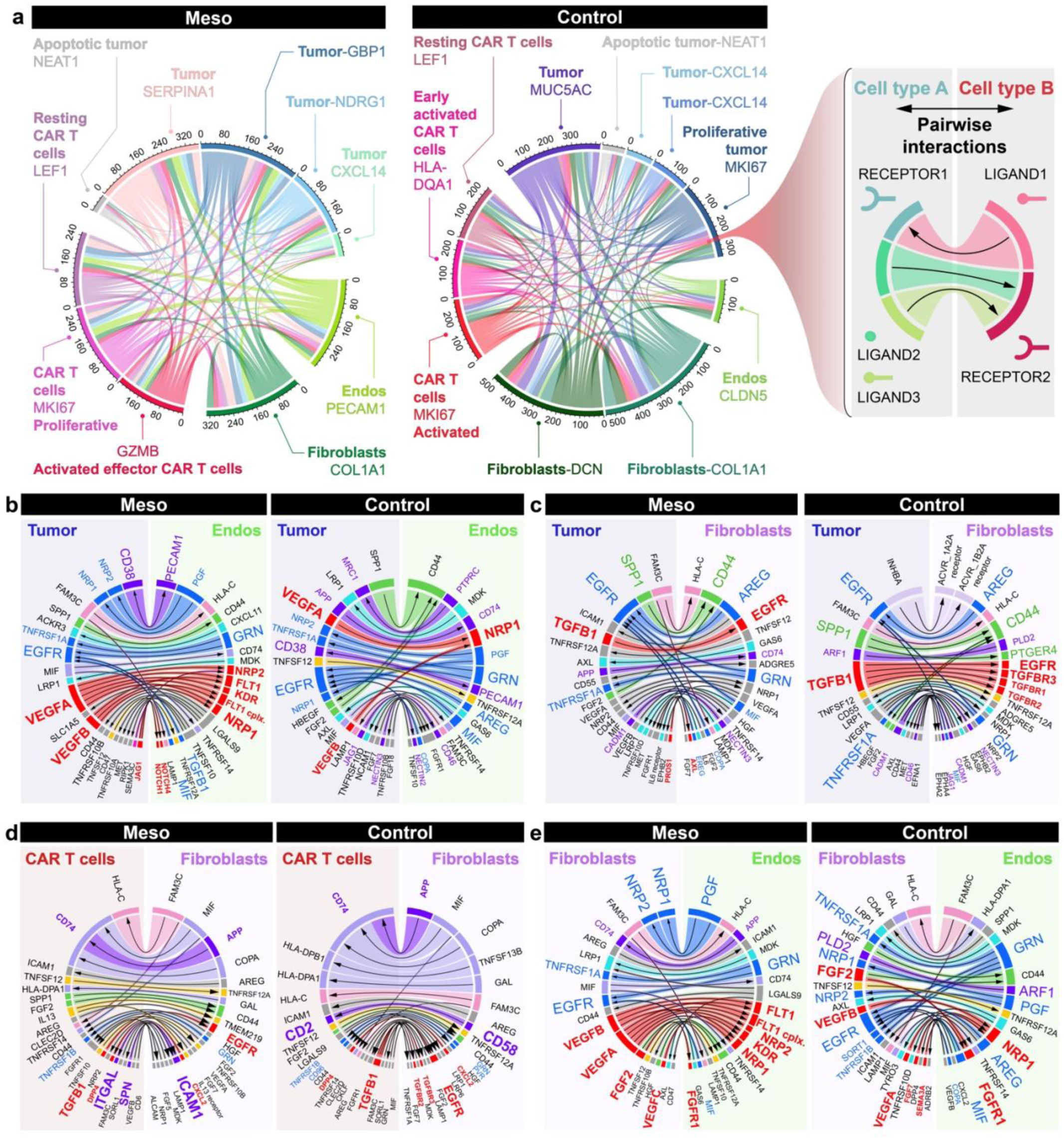
Analysis of ligand-receptor interactions in a model of human lung adenocarcinoma tumors infused with meso-CAR T cells. **a,** Chord diagrams showing an overview of intercellular communication and the number of identified interactions in the meso-tumor (left) and control (middle) groups. Each chord represents a bundle of paired and statistically significant ligand-receptor interactions between a particular pair of cell types (right). The width of the chord indicates the number of interacting ligand-receptor pairs. **b-e,** Chord diagrams of cell pair-specific ligand-receptor interactions with adjusted p-values < 0.05 and total mean expression > 0.35.

**Extended Data Fig. 3.**
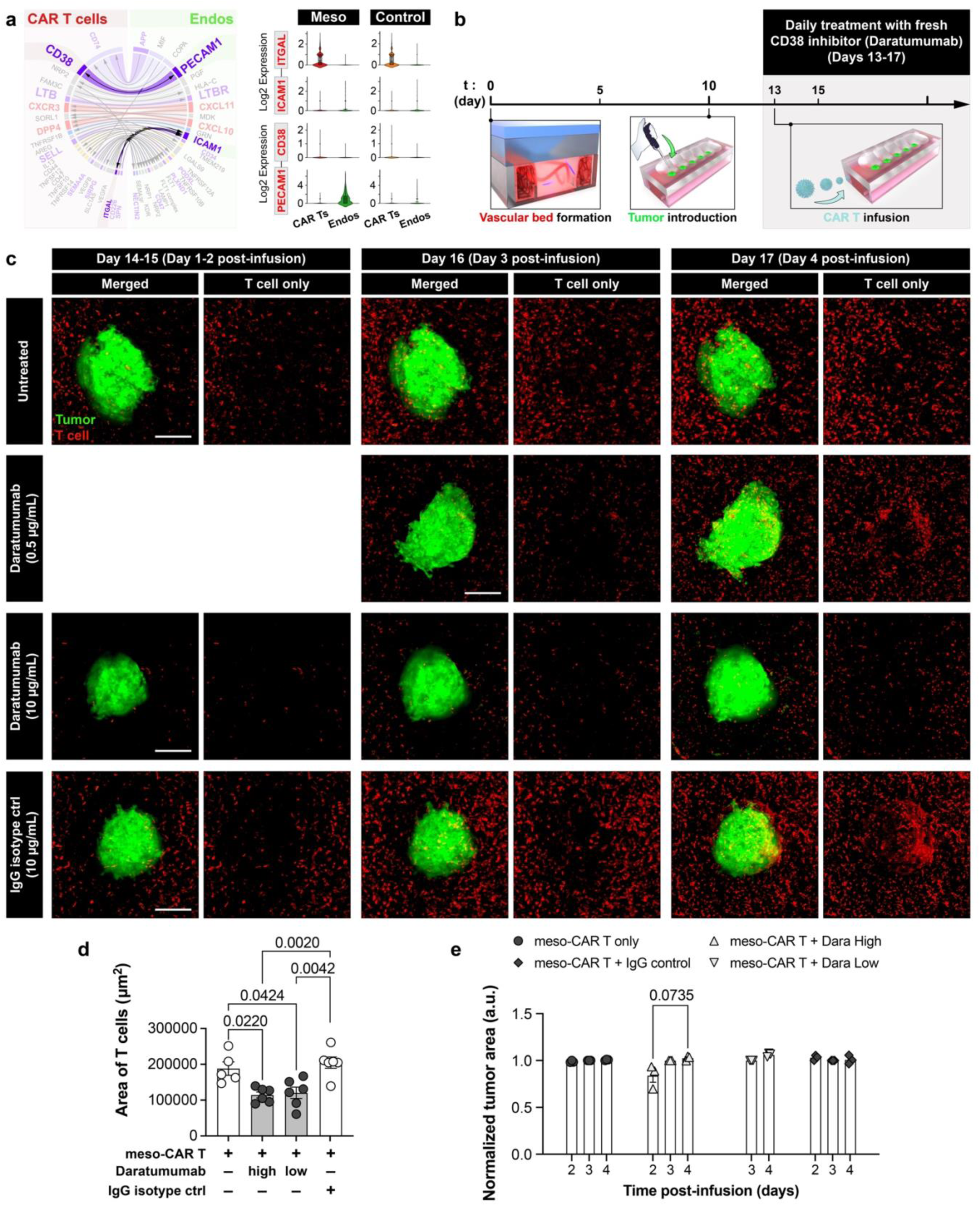
Pharmacological inhibition of CAR T-endothelial interactions mediated by CD38-PECAM1 signaling. **a,** Visualization of the interacting pairs of ITGAL-ICAM1 and CD38-PECAM1 in the meso-tumor model (left) and violin plots comparing the expression of interacting genes of interest mediating the crosstalk between CAR T cells and endothelial cells (right). **b,** Experimental timeline for CAR T cell infusion and drug treatment. **c,** Representative fluorescence micrographs of single meso-tumors infused with meso-CAR T cells without drug treatment (Untreated), treated with Daratumumab (0.5 or 10 μg/ml), or IgG isotype control antibody (10 μg/ml IgG). Blood vessels are not shown in these images. Scale bars, 250 μm. **d,e,** Quantification of T cell area (**d**) and normalized tumor area (**e**) over time. Data are presented as mean ± SEM (n ≥ 3). CAR T cells derived from one healthy donor were tested.

**Extended Data Fig. 4.**
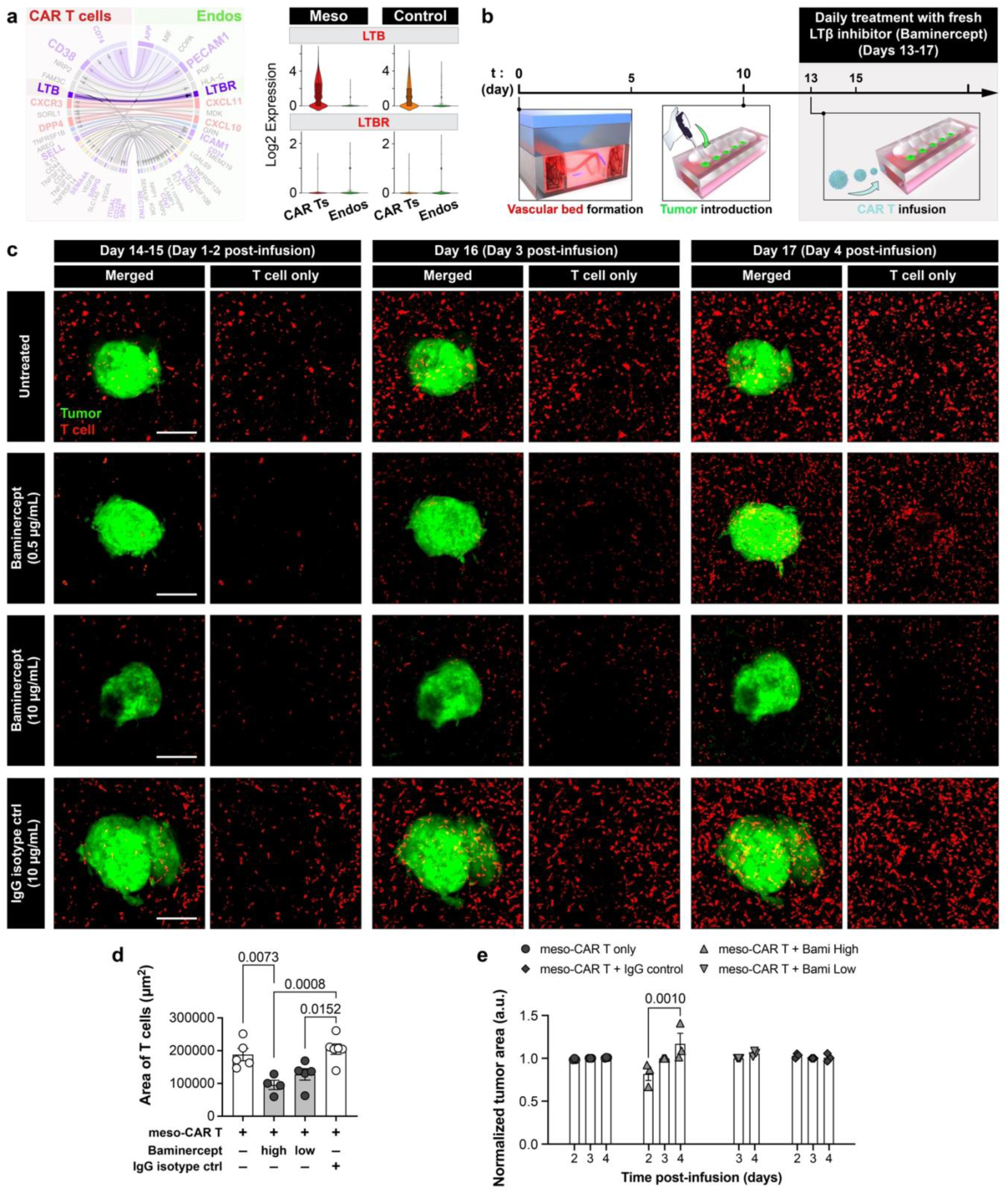
Pharmacological inhibition of CAR T-endothelial interactions mediated by LTB-LTBR signaling. **a,** Visualization of the interacting LTB-LTBR pair in the meso-tumor model (left) and violin plots comparing the expression of interacting genes of interest mediating the crosstalk between CAR T cells and endothelial cells (right). **b,** Experimental timeline for CAR T cell infusion and drug treatment. **c,** Representative fluorescence micrographs of single meso-tumors infused with meso-CAR T cells without drug treatment (Untreated), treated with Baminercept (0.5 or 10 μg/ml), or IgG isotope control antibody (10 μg/ml IgG). Blood vessels are not shown in these images. Scale bars, 250 μm. **d,e,** Quantification of (**d**) T cell area and (**e**) normalized tumor area over time. Data are presented as mean ± SEM (n ≥ 3). CAR T cells derived from one healthy donor were tested.

**Extended Data Fig. 5.**
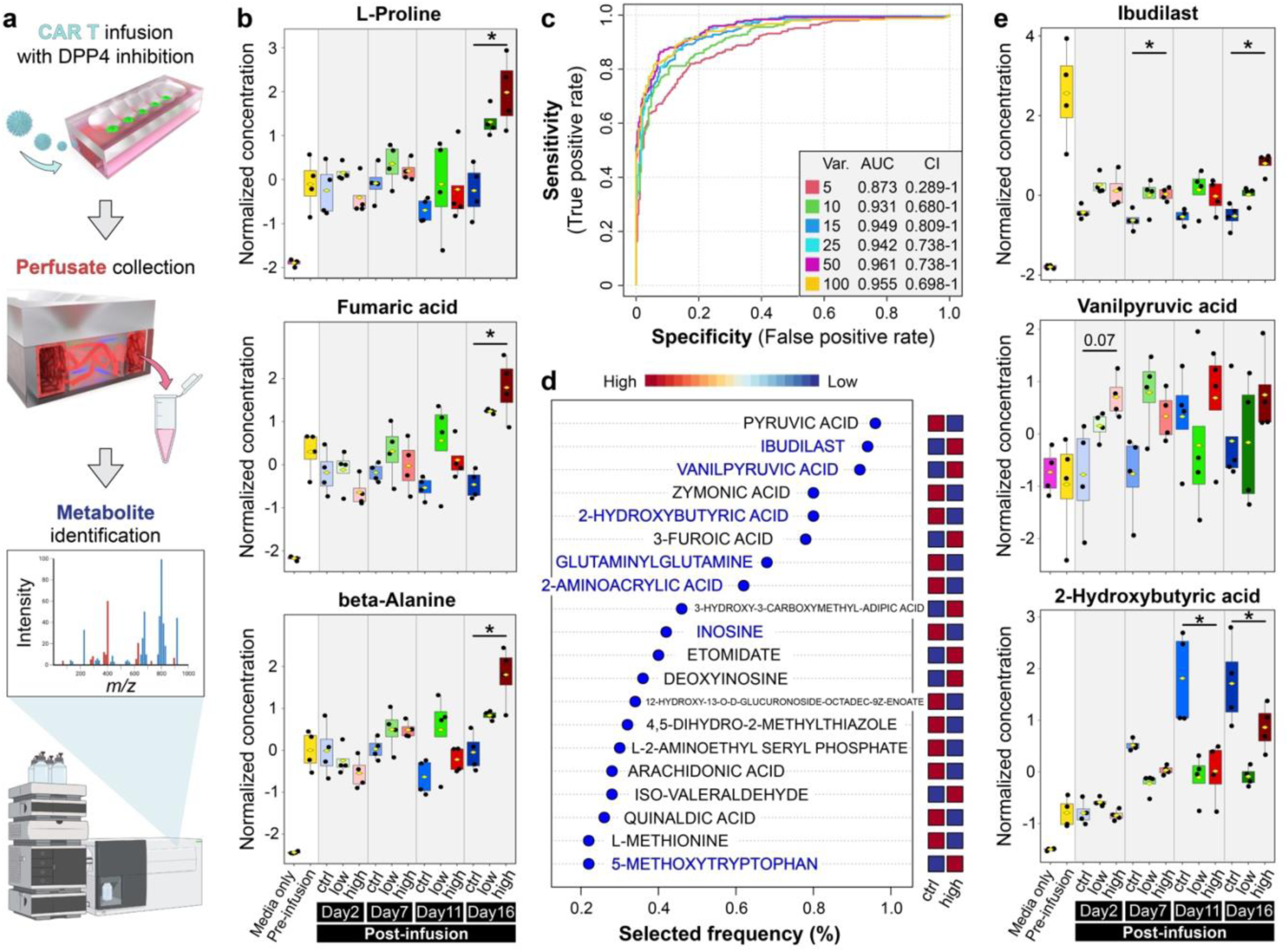
Identification of metabolomic signatures indicating the outcome of CAR T therapy with LAF237. **a,** Workflow of untargeted, global profiling of metabolites in vascular perfusate. Created with Biorender.com. **b,** Comparison of normalized concentrations of select metabolites across all conditions. **c,** ROC curves for biomarker prediction models to identify metabolites that can differentiate more efficacious CAR T cell treatment with high-dose LAF237 from control treatment at all time points post-infusion. Different numbers of constituent metabolite features per model are indicated by different colors. **d,** List of top metabolites yielded by the 15-feature prediction model shown in **c** and ranked according to their predictive accuracy. Common metabolites that appear here and in Fig. 7h are labelled blue. **e,** Comparison of normalized concentrations of metabolites selected from **d**. Boxplots show minimum, 25^th^ percentile, mean (yellow closed diamond), 75^th^ percentile, and maximum. N = 4 for each group. Note that post-hoc pairwise comparison was calculated between the high-dose (high) and control (ctrl) groups at all time points with *p < 0.05.

## Acknowledgements

We thank Mengxi Luo, Andrei Georgescu, Jeong Min Oh, Alexander C. Huang, Jennifer E. Wu, Jean-Christophe Beltra, Josephine R. Giles, Cécile Alanio, Derek Oldridge, Emma Niemeyer, Yun Chang Choi, and Cherie R. Kagan for their intellectual input and technical assistance. We also thank Renata Pellegrino Da Silva, Fernanda Mafra, Jacqueline M. Smiler, James P. Garifallou, and Michael V. Gonzalez from the Center for Applied Genomics at The Children’s Hospital of Philadelphia, and Jonathan Schug and Olga Smirnova from the Next-Generation Sequencing Core at the University of Pennsylvania for their technical assistance with single cell RNA-sequencing. We thank Tyler Skinner and Daniel Martinez from the Pathology Core Laboratory at The Children’s Hospital of Philadelphia for their technical assistance with immunohistochemistry. We thank Drs. Sunil Singhal, Melissa Culligan and Joseph Friedberg from the Hospital of the University of Pennsylvania and the Temple University Hospital for their assistance with mesothelioma patient explants. This work was supported by the Cancer Research Institute (DDH), the National Institutes of Health (NIH) (1DP2HL127720-01 for DDH; F32DK127843 for WDL; R01CA163591 and DP1DK113643 for JDR), the National Science Foundation (CMMI:15-48571) (DDH), the Ministry of Trade, Industry & Energy of the Republic of Korea (DDH), the GRDC Cooperative Hub through the National Research Foundation of Korea funded by the Ministry of Science and ICT (RS-2023-00259341) (TK, DDH), the Paul Allen Foundation (JDR), Ludwig Cancer Research (JDR), and the University of Pennsylvania. H.L. is a recipient of the Penn Institute for Regenerative Medicine Postdoctoral Fellowship and the Penn Center for Engineering MechanoBiology Pilot Grant Award. D.D.H. is a recipient of the NIH Director’s New Innovator Award and the Cancer Research Institute Technology Impact Award.

## Author contributions

H.L. designed the research, performed the experiments, analyzed the data, created the figures, and wrote the manuscript. E.N.-O., X.S., M.L., M.C.M., J.J.B., S.K., E.K.M., and S.M.A. provided materials, performed the experiments, and analyzed the data. Z.C. and E.J.W. provided assistance in the design of research and the analysis of the single-cell RNA sequencing data. W.D.L. and J.D.R. performed the experiments and provided assistance in the analysis of the metabolomics data. X.D., J.C., Y.L., A.W., Z.O., J.P., J.Y.P., A.L., and H.H. provided assistance in the experiments. S.A.A., Y.S., D.M.R., S.K., G.S.W., W.G., and T.K. provided assistance in the experiments and analyzed the data. Z.X. and E.P. provided materials and analyzed the data. D.D.H. designed the research, analyzed the data, and wrote the manuscript.

## Data availability

The scRNA-seq datasets generated and analyzed during the current study is available at the NCBI Gene Expression Omnibus, under accession number GSE240121. All other relevant data supporting the key findings of this study are available within the article and its Supplementary Information files. The raw images are too large for public deposit and are available from the corresponding author upon reasonable request.

## References

1. Muri, J. et al. The thioredoxin-1 system is essential for fueling DNA synthesis during T-cell metabolic reprogramming and proliferation. Nat Commun 9, 1851 (2018).

2. Weinreich, M. A. et al. KLF2 Transcription-Factor Deficiency in T Cells Results in Unrestrained Cytokine Production and Upregulation of Bystander Chemokine Receptors. Immunity 31, 122–130 (2009).

3. Hayashi, K., Jutabha, P., Endou, H., Sagara, H. & Anzai, N. LAT1 Is a Critical Transporter of Essential Amino Acids for Immune Reactions in Activated Human T Cells. The Journal of Immunology 191, 4080–4085 (2013).

4. Tenbrock, K., Juang, Y.-T., Tolnay, M. & Tsokos, G. C. The Cyclic Adenosine 5′-Monophosphate Response Element Modulator Suppresses IL-2 Production in Stimulated T Cells by a Chromatin-Dependent Mechanism. The Journal of Immunology 170, 2971–2976 (2003).

5. Rajah, R., Valentinis, B. & Cohen, P. Insulin-like Growth Factor (IGF)-binding Protein-3 Induces Apoptosis and Mediates the Effects of Transforming Growth Factor-β1 on Programmed Cell Death through a p53- and IGF-independent Mechanism. Journal of Biological Chemistry 272, 12181–12188 (1997).

6. Stein, S. et al. NDRG1 Is Necessary for p53-dependent Apoptosis. Journal of Biological Chemistry 279, 48930–48940 (2004).

7. Rosenberg, S. A. & Restifo, N. P. Adoptive cell transfer as personalized immunotherapy for human cancer. Science (1979) 348, 62–68 (2015).

8. Rosenberg, S. A., Restifo, N. P., Yang, J. C., Morgan, R. A. & Dudley, M. E. Adoptive cell transfer: a clinical path to effective cancer immunotherapy. Nature Reviews Cancer 2008 8:4 8, 299–308 (2008).

9. Sadelain, M., Rivière, I. & Riddell, S. Therapeutic T cell engineering. Nature 545, 423–431 (2017).

10. June, C. H. & Sadelain, M. Chimeric Antigen Receptor Therapy. New England Journal of Medicine 379, 64–73 (2018).

11. June, C. H., O’Connor, R. S., Kawalekar, O. U., Ghassemi, S. & Milone, M. C. CAR T cell immunotherapy for human cancer. Science (1979) 359, 1361–1365 (2018).

12. Porter, D. L., Levine, B. L., Kalos, M., Bagg, A. & June, C. H. Chimeric Antigen Receptor–Modified T Cells in Chronic Lymphoid Leukemia. New England Journal of Medicine 365, 725–733 (2011).

13. Maude, S. L. et al. Chimeric Antigen Receptor T Cells for Sustained Remissions in Leukemia. New England Journal of Medicine 371, 1507–1517 (2014).

14. Lee, D. W. et al. T cells expressing CD19 chimeric antigen receptors for acute lymphoblastic leukaemia in children and young adults: A phase 1 dose-escalation trial. The Lancet 385, 517–528 (2015).

15. Sterner, R. C. & Sterner, R. M. CAR-T cell therapy: current limitations and potential strategies. Blood Cancer J 11, (2021).

16. Neelapu, S. S. et al. Axicabtagene Ciloleucel CAR T-Cell Therapy in Refractory Large B-Cell Lymphoma. New England Journal of Medicine 377, 2531–2544 (2017).

17. Schuster, S. J. et al. Chimeric Antigen Receptor T Cells in Refractory B-Cell Lymphomas. New England Journal of Medicine 377, 2545–2554 (2017).

18. Newick, K., O’Brien, S., Moon, E. & Albelda, S. M. CAR T Cell Therapy for Solid Tumors. Annu Rev Med 68, 139–152 (2017).

19. Louis, C. U. et al. Antitumor activity and long-term fate of chimeric antigen receptor-positive T cells in patients with neuroblastoma. Blood 118, 6050–6056 (2011).

20. Shah, N. N. & Fry, T. J. Mechanisms of resistance to CAR T cell therapy. Nat Rev Clin Oncol 16, 372–385 (2019).

21. Schmidts, A. & Maus, M. V. Making CAR T cells a solid option for solid tumors. Front Immunol 9, 1–10 (2018).

22. Lim, W. A. & June, C. H. The Principles of Engineering Immune Cells to Treat Cancer. Cell 168, 724–740 (2017).

23. Joyce, J. A. & Fearon, D. T. T cell exclusion, immune privilege, and the tumor microenvironment. Science (1979) 348, 74–80 (2015).

24. Ramakrishna, S., Barsan, V. & Mackall, C. Prospects and challenges for use of CAR T cell therapies in solid tumors. Expert Opin Biol Ther 20, 503–516 (2020).

25. Larson, R. C. & Maus, M. V. Recent advances and discoveries in the mechanisms and functions of CAR T cells. Nat Rev Cancer (2021) doi:10.1038/s41568-020-00323-z.

26. Bagley, S. J., Desai, A. S., Linette, G. P., June, C. H. & O’Rourke, D. M. CAR T-cell therapy for glioblastoma: Recent clinical advances and future challenges. Neuro Oncol 20, 1429–1438 (2018).

27. Junttila, M. R. & De Sauvage, F. J. Influence of tumour micro-environment heterogeneity on therapeutic response. Nature 501, 346–354 (2013).

28. Sackstein, R., Schatton, T. & Barthel, S. R. T-lymphocyte homing: An underappreciated yet critical hurdle for successful cancer immunotherapy. Laboratory Investigation 97, 669–697 (2017).

29. Martinez, M. & Moon, E. K. CAR T Cells for Solid Tumors: New Strategies for Finding, Infiltrating, and Surviving in the Tumor Microenvironment. Front Immunol 10, 128 (2019).

30. Zitvogel, L., Pitt, J. M., Daillère, R., Smyth, M. J. & Kroemer, G. Mouse models in oncoimmunology. Nature Reviews Cancer 2016 16:12 16, 759–773 (2016).

31. Gengenbacher, N., Singhal, M. & Augustin, H. G. Preclinical mouse solid tumour models: status quo, challenges and perspectives. Nat Rev Cancer 17, 751–765 (2017).

32. Srivastava, S. & Riddell, S. R. Chimeric Antigen Receptor T Cell Therapy: Challenges to Bench-to-Bedside Efficacy. The Journal of Immunology 200, 459–468 (2018).

33. Day, C. P., Merlino, G. & Van Dyke, T. Preclinical Mouse Cancer Models: A Maze of Opportunities and Challenges. Cell 163, 39–53 (2015).

34. Moon, E. K. et al. Multifactorial T-cell Hypofunction That Is Reversible Can Limit the Efficacy of Chimeric Antigen Receptor–Transduced Human T cells in Solid Tumors. Clinical Cancer Research 20, 4262–4273 (2014).

35. Carpenito, C. et al. Control of large, established tumor xenografts with genetically retargeted human T cells containing CD28 and CD137 domains. Proceedings of the National Academy of Sciences 106, 3360–3365 (2009).

36. Chuprin, J. et al. Humanized mouse models for immuno-oncology research. Nature Reviews Clinical Oncology 2023 1–15 (2023) doi:10.1038/s41571-022-00721-2.

37. Ma, X. et al. Interleukin-23 engineering improves CAR T cell function in solid tumors. Nat Biotechnol 38, (2020).

38. Avanzi, M. P. et al. Engineered Tumor-Targeted T Cells Mediate Enhanced Anti-Tumor Efficacy Both Directly and through Activation of the Endogenous Immune System. Cell Rep 23, 2130–2141 (2018).

39. Wang, L. C. S. et al. Targeting fibroblast activation protein in tumor stroma with chimeric antigen receptor T cells can inhibit tumor growth and augment host immunity without severe toxicity. Cancer Immunol Res 2, 154–166 (2014).

40. Klampatsa, A. et al. Analysis and Augmentation of the Immunologic Bystander Effects of CAR T Cell Therapy in a Syngeneic Mouse Cancer Model. Mol Ther Oncolytics 18, 360–371 (2020).

41. Evgin, L. et al. Oncolytic virus–mediated expansion of dual-specific CAR T cells improves efficacy against solid tumors in mice. Sci Transl Med 14, 2231 (2022).

42. Duncan, B. B., Dunbar, C. E. & Ishii, K. Applying a clinical lens to animal models of CAR-T cell therapies. Mol Ther Methods Clin Dev 27, 17–31 (2022).

43. Elsallab, M., Bravery, C. A., Kurtz, A. & Abou-El-Enein, M. Mitigating Deficiencies in Evidence during Regulatory Assessments of Advanced Therapies: A Comparative Study with Other Biologicals. Mol Ther Methods Clin Dev 18, 269–279 (2020).

44. Abou-el-Enein, M. et al. Evidence generation and reproducibility in cell and gene therapy research: A call to action. Mol Ther Methods Clin Dev 22, 11–14 (2021).

45. McKean, M. et al. Safety and early efficacy results from a phase 1, multicenter trial of PSMA-targeted armored CAR T cells in patients with advanced mCRPC. Journal of Clinical Oncology 40, 94–94 (2022).

46. Kloss, C. C. et al. Dominant-Negative TGF-β Receptor Enhances PSMA-Targeted Human CAR T Cell Proliferation And Augments Prostate Cancer Eradication. Molecular Therapy 26, 1855–1866 (2018).

47. Mak, I. W. Y., Evaniew, N. & Ghert, M. Lost in translation: animal models and clinical trials in cancer treatment. Am J Transl Res 6, 114 (2014).

48. Hegde, P. S. & Chen, D. S. Top 10 Challenges in Cancer Immunotherapy. Immunity 52, 17–35 (2020).

49. Elmore, L. W. et al. Blueprint for cancer research: Critical gaps and opportunities. CA Cancer J Clin 71, 107–139 (2021).

50. Vanmeerbeek, I., Naulaerts, S. & Garg, A. D. Reverse translation: the key to increasing the clinical success of immunotherapy? Genes Immun 24, 217–219 (2023).

51. Hidalgo, M. et al. Patient-derived Xenograft models: An emerging platform for translational cancer research. Cancer Discov 4, 998–1013 (2014).

52. Risau, W. & Flamme, I. Vasculogenesis. Annu Rev Cell Dev Biol 11, 73–91 (1995).

53. Egeblad, M., Nakasone, E. S. & Werb, Z. Tumors as Organs: Complex Tissues that Interface with the Entire Organism. Dev Cell 18, 884–901 (2010).

54. Li, L. et al. Laminin γ2–mediating T cell exclusion attenuates response to anti–PD-1 therapy. Sci Adv 7, eabc8346 (2021).

55. Ho, M. et al. Mesothelin expression in human lung cancer. Clinical Cancer Research 13, 1571–1575 (2007).

56. Hassan, R. et al. Mesothelin immunotherapy for cancer: Ready for prime time? Journal of Clinical Oncology 34, 4171–4179 (2016).

57. Pastan, I. & Hassan, R. Discovery of mesothelin and exploiting it as a target for immunotherapy. Cancer Res 74, 2907–2912 (2014).

58. Klampatsa, A., Dimou, V. & Albelda, S. M. Mesothelin-targeted CAR-T cell therapy for solid tumors. Expert Opin Biol Ther 21, 473–486 (2021).

59. Tanyi, J. et al. Phase I study of autologous T cells bearing fully-humanized chimeric antigen receptors targeting mesothelin in mesothelin-expressing cancers (314). Gynecol Oncol 166, S164–S165 (2022).

60. MacKay, M. et al. The therapeutic landscape for cells engineered with chimeric antigen receptors. Nat Biotechnol 38, 233–244 (2020).

61. Saez-Ibañez, A. R. et al. Landscape of cancer cell therapies: trends and real-world data. Nat Rev Drug Discov 21, 631–632 (2022).

62. Ross, S. H. & Cantrell, D. A. Signaling and Function of Interleukin-2 in T Lymphocytes. Annu Rev Immunol 36, 411–433 (2018).

63. Liu, Y. et al. IL-2 regulates tumor-reactive CD8+ T cell exhaustion by activating the aryl hydrocarbon receptor. Nat Immunol 22, 358–369 (2021).

64. Mariathasan, S. et al. TGFβ attenuates tumour response to PD-L1 blockade by contributing to exclusion of T cells. Nature 554, 544–548 (2018).

65. Philip, M. & Schietinger, A. CD8+ T cell differentiation and dysfunction in cancer. Nat Rev Immunol 22, 209–223 (2022).

66. O’Brien, S. M. et al. Function of Human Tumor-Infiltrating Lymphocytes in Early-Stage Non–Small Cell Lung Cancer. Cancer Immunol Res 7, 896–909 (2019).

67. Hombrink, P. et al. Programs for the persistence, vigilance and control of human CD8 + lung-resident memory T cells. Nat Immunol 17, 1467–1478 (2016).

68. Ganesan, A. P. et al. Tissue-resident memory features are linked to the magnitude of cytotoxic T cell responses in human lung cancer. Nat Immunol 18, 940–950 (2017).

69. Klampatsa, A. et al. Phenotypic and functional analysis of malignant mesothelioma tumor-infiltrating lymphocytes. Oncoimmunology 8, 1–12 (2019).

70. Sallusto, F., Geginat, J. & Lanzavecchia, A. Central memory and effector memory T cell subsets: Function, generation, and maintenance. Annu Rev Immunol 22, 745–763 (2004).

71. Kaech, S. M. & Cui, W. Transcriptional control of effector and memory CD8+ T cell differentiation. Nat Rev Immunol 12, 749–761 (2012).

72. Janciauskiene, S. et al. Clinical Significance of SERPINA1 Gene and Its Encoded Alpha1-Antitrypsin Protein in NSCLC. Cancers (Basel) 11, (2019).

73. Hall, J. C. et al. Precise probes of type II interferon activity define the origin of interferon signatures in target tissues in rheumatic diseases. Proc Natl Acad Sci U S A 109, 17609–17614 (2012).

74. Manalo, D. J. et al. Transcriptional regulation of vascular endothelial cell responses to hypoxia by HIF-1. Blood 105, 659–669 (2005).

75. Boroughs, A. C. et al. A Distinct Transcriptional Program in Human CAR T Cells Bearing the 4-1BB Signaling Domain Revealed by scRNA-Seq. Molecular Therapy 28, 1–16 (2020).

76. Dumartin, L. et al. ER stress protein AGR2 precedes and is involved in the regulation of pancreatic cancer initiation. Oncogene 36, 3094–3103 (2017).

77. Szabo, P. A. et al. Single-cell transcriptomics of human T cells reveals tissue and activation signatures in health and disease. Nat Commun 10, (2019).

78. Guo, X. et al. Global characterization of T cells in non-small-cell lung cancer by single-cell sequencing. Nat Med 24, 978–985 (2018).

79. Uslu, U., Castelli, S. & June, C. H. CAR T cell combination therapies to treat cancer. Cancer Cell 42, 1319–1325 (2024).

80. Craddock, J. A. et al. Enhanced Tumor Trafficking of GD2 Chimeric Antigen Receptor T Cells by Expression of the Chemokine Receptor CCR2b. Journal of Immunotherapy 33, 780–788 (2010).

81. Gueugnon, F. et al. Identification of novel markers for the diagnosis of malignant pleural mesothelioma. American Journal of Pathology 178, 1033–1042 (2011).

82. Servais, E. L. et al. Mesothelin overexpression promotes mesothelioma cell invasion and MMP-9 secretion in an orthotopic mouse model and in epithelioid pleural mesothelioma patients. Clinical Cancer Research 18, 2478–2489 (2012).

83. Grosser, R., Cherkassky, L., Chintala, N. & Adusumilli, P. S. Combination Immunotherapy with CAR T Cells and Checkpoint Blockade for the Treatment of Solid Tumors. Cancer Cell 36, 471–482 (2019).

84. Efremova, M., Vento-Tormo, M., Teichmann, S. A. & Vento-Tormo, R. CellPhoneDB: inferring cell–cell communication from combined expression of multi-subunit ligand–receptor complexes. Nat Protoc 15, 1484–1506 (2020).

85. Schröder, B. The multifaceted roles of the invariant chain CD74 - More than just a chaperone. Biochim Biophys Acta Mol Cell Res 1863, 1269–1281 (2016).

86. Maurer, K. et al. Expansion of a CD8+ Temra Population and Activating T-Cell Interactions Characterize the Graft Versus Leukemia Response in Relapsed AML. Blood 140, 4806–4807 (2022).

87. Wang, J. H. et al. Structure of a heterophilic adhesion complex between the human CD2 and CD58 (LFA-3) counterreceptors. Cell 97, 791–803 (1999).

88. de Palma, M., Biziato, D. & Petrova, T. v. Microenvironmental regulation of tumour angiogenesis. Nat Rev Cancer 17, 457–474 (2017).

89. Asrir, A. et al. Tumor-associated high endothelial venules mediate lymphocyte entry into tumors and predict response to PD-1 plus CTLA-4 combination immunotherapy. Cancer Cell (2022) doi:10.1016/J.CCELL.2022.01.002.

90. Choi, J., Enis, D. R., Koh, K. P., Shiao, S. L. & Pober, J. S. T lymphocyte-endothelial cell interactions. Annu Rev Immunol 22, 683–709 (2004).

91. Motz, G. T. et al. Tumor endothelium FasL establishes a selective immune barrier promoting tolerance in tumors. Nat Med 20, 607–615 (2014).

92. Casneuf, T. et al. Effects of daratumumab on natural killer cells and impact on clinical outcomes in relapsed or refractory multiple myeloma. Blood Adv 1, 2105–2114 (2017).

93. St.Clair, E. W., et al. Clinical Efficacy and Safety of Baminercept, a Lymphotoxin β Receptor Fusion Protein, in Primary Sjögren’s Syndrome: Results From a Phase II Randomized, Double-Blind, Placebo-Controlled Trial. Arthritis and Rheumatology 70, 1470–1480 (2018).

94. Tokunaga, R. et al. CXCL9, CXCL10, CXCL11/CXCR3 axis for immune activation – A target for novel cancer therapy. Cancer Treat Rev 63, 40–47 (2018).

95. Nagarsheth, N., Wicha, M. S. & Zou, W. Chemokines in the cancer microenvironment and their relevance in cancer immunotherapy. Nat Rev Immunol 17, 559–572 (2017).

96. Casrouge, A. et al. Evidence for an antagonist form of the chemokine CXCL10 in patients chronically infected with HCV. Journal of Clinical Investigation 121, 308–317 (2011).

97. Morello, A., Sadelain, M. & Adusumilli, P. S. Mesothelin-targeted CARs: Driving T cells to solid Tumors. Cancer Discov 6, 133–146 (2016).

98. Kachala, S. S. et al. Mesothelin overexpression is a marker of tumor aggressiveness and is associated with reduced recurrence-free and overall survival in early-stage lung adenocarcinoma. Clinical Cancer Research 20, 1020–1028 (2014).

99. Klampatsa, A. et al. Analysis and Augmentation of the Immunologic Bystander Effects of CAR T Cell Therapy in a Syngeneic Mouse Cancer Model. Mol Ther Oncolytics 18, 360–371 (2020).

100. Zhao, Y. et al. Multiple injections of electroporated autologous T cells expressing a chimeric antigen receptor mediate regression of human disseminated tumor. Cancer Res 70, 9053–9061 (2010).

101. Santos, A. M., Jung, J., Aziz, N., Kissil, J. L. & Puré, E. Targeting fibroblast activation protein inhibits tumor stromagenesis and growth in mice. Journal of Clinical Investigation 119, 3613–3625 (2009).

102. Villhauer, E. B. et al. 1-[[(3-Hydroxy-1-adamantyl)amino]acetyl]-2-cyano-(*S*)-pyrrolidine: A Potent, Selective, and Orally Bioavailable Dipeptidyl Peptidase IV Inhibitor with Antihyperglycemic Properties. J Med Chem 46, 2774–2789 (2003).

103. Hu, P. et al. Pharmacokinetics and pharmacodynamics of vildagliptin in healthy Chinese volunteers. J Clin Pharmacol 49, 39–49 (2009).

104. He, Y. L. Clinical pharmacokinetics and pharmacodynamics of vildagliptin. Clin Pharmacokinet 51, 147–162 (2012).

105. Madden, M. Z. & Rathmell, J. C. The complex integration of t-cell metabolism and immunotherapy. Cancer Discov 11, 1636–1643 (2021).

106. DePeaux, K. & Delgoffe, G. M. Metabolic barriers to cancer immunotherapy. Nature Reviews Immunology 2021 21:12 21, 785–797 (2021).

107. Maulana, T. I. et al. Immunocompetent cancer-on-chip models to assess immuno-oncology therapy. Adv Drug Deliv Rev 173, 281–305 (2021).

108. Lam, M. S. Y. et al. G9a/GLP inhibition during ex vivo lymphocyte expansion increases in vivo cytotoxicity of engineered T cells against hepatocellular carcinoma. Nat Commun 14, (2023).

109. Mollica, H. et al. A 3D pancreatic tumor model to study T cell infiltration. Biomater Sci 9, 7420–7431 (2021).

110. Wallstabe, L., et al. ROR1-CAR T cells are effective against lung and breast cancer in advanced microphysiologic 3D tumor models. JCI Insight 4, (2019).

111. Ayuso, J. M. et al. Microfluidic tumor-on-a-chip model to evaluate the role of tumor environmental stress on NK cell exhaustion. Sci Adv 7, 1–15 (2021).

112. Sontheimer-Phelps, A., Hassell, B. A. & Ingber, D. E. Modelling cancer in microfluidic human organs-on-chips. Nat Rev Cancer 19, 65–81 (2019).

113. Nguyen, H. T. et al. Patient-specific vascularized tumor model: Blocking monocyte recruitment with multispecific antibodies targeting CCR2 and CSF-1R. Biomaterials 312, 122731 (2025).

114. Pavesi, A. et al. A 3D microfluidic model for preclinical evaluation of TCR-engineered T cells against solid tumors. JCI Insight 2, (2017).

115. Maulana, T. I. et al. Breast cancer-on-chip for patient-specific efficacy and safety testing of CAR-T cells. Cell Stem Cell (2024) doi:10.1016/j.stem.2024.04.018.

116. Barrett, R. & Puré, E. Cancer-associated fibroblasts: key determinants of tumor immunity and immunotherapy. Curr Opin Immunol 64, 80–87 (2020).

117. Ishii, G., Ochiai, A. & Neri, S. Phenotypic and functional heterogeneity of cancer-associated fibroblast within the tumor microenvironment. Adv Drug Deliv Rev 99, 186–196 (2016).

118. Sahai, E. et al. A framework for advancing our understanding of cancer-associated fibroblasts. Nat Rev Cancer 20, 174–186 (2020).

119. Chen, X. & Song, E. Turning foes to friends: targeting cancer-associated fibroblasts. Nat Rev Drug Discov 18, 99–115 (2019).

120. Sharon, Y. et al. Tumor-Derived Osteopontin Reprograms Normal Mammary Fibroblasts to Promote Inflammation and Tumor Growth in Breast Cancer. Cancer Res 75, 963–973 (2015).

121. Hu, H. et al. Three subtypes of lung cancer fibroblasts define distinct therapeutic paradigms. Cancer Cell 1–17 (2021) doi:10.1016/j.ccell.2021.09.003.

122. Buckanovich, R. J. et al. Endothelin B receptor mediates the endothelial barrier to T cell homing to tumors and disables immune therapy. Nat Med 14, 28–36 (2008).

123. Pathria, P., Louis, T. L. & Varner, J. A. Targeting Tumor-Associated Macrophages in Cancer. Trends Immunol 40, 310–327 (2019).

124. Sautès-Fridman, C., Petitprez, F., Calderaro, J. & Fridman, W. H. Tertiary lymphoid structures in the era of cancer immunotherapy. Nat Rev Cancer 19, 307–325 (2019).

125. Engelhard, V. H. et al. Immune Cell Infiltration and Tertiary Lymphoid Structures as Determinants of Antitumor Immunity. The Journal of Immunology 200, 432–442 (2018).

126. Schumacher, T. N. & Thommen, D. S. Tertiary lymphoid structures in cancer. Science (1979) 375, eabf9419 (2022).

127. Thommen, D. S. et al. A transcriptionally and functionally distinct PD-1+ CD8+ T cell pool with predictive potential in non-small-cell lung cancer treated with PD-1 blockade. Nat Med 24, 994–1004 (2018).

128. de Chaisemartin, L. et al. Characterization of Chemokines and Adhesion Molecules Associated with T cell Presence in Tertiary Lymphoid Structures in Human Lung Cancer. Cancer Res 71, 6391–6399 (2011).

129. Rodriguez-Garcia, A. et al. CAR-T cell-mediated depletion of immunosuppressive tumor-associated macrophages promotes endogenous antitumor immunity and augments adoptive immunotherapy. Nature Communications 2021 12:1 12, 1–17 (2021).

130. Sánchez-Paulete, A. R. et al. Targeting Macrophages with CAR T Cells Delays Solid Tumor Progression and Enhances Antitumor Immunity. Cancer Immunol Res 10, 1354–1369 (2022).

131. Brandt, I. et al. Inhibition of dipeptidyl-peptidase IV catalyzed peptide truncation by Vildagliptin ((2S)-{[(3-hydroxyadamantan-1-yl)amino]acetyl}-pyrrolidine-2-carbonitrile). Biochem Pharmacol 70, 134–143 (2005).

132. Mathieu, C. & Degrande, E. Vildagliptin: a new oral treatment for type 2 diabetes mellitus. Vasc Health Risk Manag 4, 1349 (2008).

133. House, I. G. et al. Macrophage-Derived CXCL9 and CXCL10 Are Required for Antitumor Immune Responses Following Immune Checkpoint Blockade. Clinical Cancer Research 26, 487–504 (2020).

134. Barreira Da Silva, R., et al. Dipeptidylpeptidase 4 inhibition enhances lymphocyte trafficking, improving both naturally occurring tumor immunity and immunotherapy. Nat Immunol 16, 850–858 (2015).

135. Nishina, S. et al. Dipeptidyl Peptidase 4 Inhibitors Reduce Hepatocellular Carcinoma by Activating Lymphocyte Chemotaxis in Mice. Cmgh 7, 115–134 (2019).

136. Ohnuma, K., Hatano, R. & Morimoto, C. DPP4 in anti-tumor immunity: Going beyond the enzyme. Nat Immunol 16, 791–792 (2015).

137. Hou, A. J., Chen, L. C. & Chen, Y. Y. Navigating CAR-T cells through the solid-tumour microenvironment. Nature Reviews Drug Discovery 2021 20:7 20, 531–550 (2021).

138. Young, R. M., Engel, N. W., Uslu, U., Wellhausen, N. & June, C. H. Next-Generation CAR T-cell Therapies. Cancer Discov OF1–OF14 (2022) doi:10.1158/2159-8290.CD-21-1683.

139. Najibi, A. J. & Mooney, D. J. Cell and tissue engineering in lymph nodes for cancer immunotherapy. Adv Drug Deliv Rev 161–162, 42–62 (2020).

140. Gill, S., Maus, M. V. & Porter, D. L. Chimeric antigen receptor T cell therapy: 25 years in the making. Blood Rev 30, 157–167 (2016).

141. Melero, I., Rouzaut, A., Motz, G. T. & Coukos, G. T-Cell and NK-Cell Infiltration into Solid Tumors: A Key Limiting Factor for Efficacious Cancer Immunotherapy. Cancer Discov 4, 522–526 (2014).

142. Spratlin, J. L., Serkova, N. J. & Eckhardt, S. G. Clinical Applications of Metabolomics in Oncology: A Review. Clinical Cancer Research 15, 431–440 (2009).

143. Kiru, L. et al. In vivo imaging of nanoparticle-labeled CAR T cells. Proceedings of the National Academy of Sciences 119, e2102363119 (2022).

144. Cheng, J. et al. Cancer-cell-derived fumarate suppresses the anti-tumor capacity of CD8+ T cells in the tumor microenvironment. Cell Metab (2023) doi:10.1016/J.CMET.2023.04.017.

145. Wang, T. et al. Inosine is an alternative carbon source for CD8+-T-cell function under glucose restriction. Nat Metab 2, 635–647 (2020).

146. McCracken, M. N. et al. Noninvasive detection of tumor-infiltrating T cells by PET reporter imaging. Journal of Clinical Investigation 125, 1815–1826 (2015).

147. O’Rourke, D. M. et al. A single dose of peripherally infused EGFRvIII-directed CAR T cells mediates antigen loss and induces adaptive resistance in patients with recurrent glioblastoma. Sci Transl Med 9, (2017).

148. Thistlethwaite, F. C. et al. The clinical efficacy of first-generation carcinoembryonic antigen (CEACAM5)-specific CAR T cells is limited by poor persistence and transient pre-conditioning-dependent respiratory toxicity. Cancer Immunology, Immunotherapy 66, 1425–1436 (2017).

149. Schmidt, D. R. et al. Metabolomics in cancer research and emerging applications in clinical oncology. CA Cancer J Clin 71, 333–358 (2021).

150. Kumar, C. S. et al. Molecular characterization of the murine interferon γ receptor cDNA. Journal of Biological Chemistry 264, 17939–17946 (1989).

151. Laskowski, T. J., Biederstädt, A. & Rezvani, K. Natural killer cells in antitumour adoptive cell immunotherapy. Nat Rev Cancer 22, 557–575 (2022).

152. Mensurado, S., Blanco-Domínguez, R. & Silva-Santos, B. The emerging roles of γδ T cells in cancer immunotherapy. Nature Reviews Clinical Oncology 2023 1–14 (2023) doi:10.1038/s41571-022-00722-1.

153. Klichinsky, M. et al. Human chimeric antigen receptor macrophages for cancer immunotherapy. Nat Biotechnol (2020) doi:10.1038/s41587-020-0462-y.

154. Tang, H. et al. Facilitating T Cell Infiltration in Tumor Microenvironment Overcomes Resistance to PD-L1 Blockade. Cancer Cell 29, 285–296 (2016).

155. Tumeh, P. C. et al. PD-1 blockade induces responses by inhibiting adaptive immune resistance. Nature 515, 568–571 (2014).

156. Rafiq, S., Hackett, C. S. & Brentjens, R. J. Engineering strategies to overcome the current roadblocks in CAR T cell therapy. Nat Rev Clin Oncol 17, 147–167 (2020).

157. Srivastava, S. et al. Immunogenic Chemotherapy Enhances Recruitment of CAR-T Cells to Lung Tumors and Improves Antitumor Efficacy when Combined with Checkpoint Blockade. Cancer Cell 39, 193–208.e10 (2021).

158. Lynn, R. C. et al. c-Jun overexpression in CAR T cells induces exhaustion resistance. Nature 576, 293–300 (2019).

159. Park, S. E., Georgescu, A., Oh, J. M., Kwon, K. W. & Huh, D. Polydopamine-Based Interfacial Engineering of Extracellular Matrix Hydrogels for the Construction and Long-Term Maintenance of Living Three-Dimensional Tissues. ACS Appl Mater Interfaces 11, 23919–23925 (2019).

160. Moon, E. K. et al. Expression of a Functional CCR2 Receptor Enhances Tumor Localization and Tumor Eradication by Retargeted Human T cells Expressing a Mesothelin-Specific Chimeric Antibody Receptor. Clinical Cancer Research 17, 4719–4730 (2011).

161. Satija, R., Farrell, J. A., Gennert, D., Schier, A. F. & Regev, A. Spatial reconstruction of single-cell gene expression data. Nature Biotechnology 2015 33:5 33, 495–502 (2015).

162. Aran, D. et al. Reference-based analysis of lung single-cell sequencing reveals a transitional profibrotic macrophage. Nat Immunol 20, 163–172 (2019).

163. Trapnell, C. et al. The dynamics and regulators of cell fate decisions are revealed by pseudotemporal ordering of single cells. Nat Biotechnol 32, 381–386 (2014).

164. Cao, J. et al. The single-cell transcriptional landscape of mammalian organogenesis. Nature 566, 496–502 (2019).

165. Vento-Tormo, R. et al. Single-cell reconstruction of the early maternal–fetal interface in humans. Nature 563, 347–353 (2018).

166. Gu, Z., Gu, L., Eils, R., Schlesner, M. & Brors, B. circlize implements and enhances circular visualization in R. Bioinformatics 30, 2811–2812 (2014).

167. Frens, G. Controlled Nucleation for the Regulation of the Particle Size in Monodisperse Gold Suspensions. Nature Physical Science 241, 20–22 (1973).

168. Chen, G. et al. Exosomal PD-L1 contributes to immunosuppression and is associated with anti-PD-1 response. Nature 560, 382–386 (2018).

169. Chambers, M. C. et al. A cross-platform toolkit for mass spectrometry and proteomics. Nature Biotechnology 2012 30:10 30, 918–920 (2012).

170. Agrawal, S. et al. EL-MAVEN: A fast, robust, and user-friendly mass spectrometry data processing engine for metabolomics. Methods in Molecular Biology 1978, 301–321 (2019).

171. Pang, Z. et al. Using MetaboAnalyst 5.0 for LC–HRMS spectra processing, multi-omics integration and covariate adjustment of global metabolomics data. Nature Protocols 2022 17:8 17, 1735–1761 (2022).

