## Supplemental Information for "Microengineered transplantation of human solid tumors for in vitro studies of CAR T immunotherapy"

**Supplementary Information includes:**

**Supplementary Figures 1 to 33**

Supplementary Figure 1. Fabrication of microdevice for in vitro tumor transplantation
Supplementary Figure 2. Injection of cell-hydrogel mixture
Supplementary Figure 3. Effect of initial tumor size on tumor vascularization
Supplementary Figure 4. Tumor nests
Supplementary Figure 5. In vitro transplantation of tumor spheroids and organoids
Supplementary Figure 6. Overexpression of mesothelin in A549 and meso-CAR design
Supplementary Figure 7. Growth of lung tumors and their responses to CAR T cells
Supplementary Figure 8. Analysis of lung tumors infused with non-transduced T cells
Supplementary Figure 9. Analysis of chemokine production
Supplementary Figure 10. Histological analysis of meso- and control tumors
Supplementary Figure 11. Flow cytometric analysis of CAR T cell phenotype
Supplementary Figure 12a. Identification of UMAP clusters
Supplementary Figure 12b. Heatmap of differentially expressed genes in meso-tumor group
Supplementary Figure 13. Gene Ontology (GO) enrichment analysis
Supplementary Figure 14. Expression of cluster-specific marker genes
Supplementary Figure 15. UMAP plots of tumor and stromal cell-related marker genes
Supplementary Figure 16. Comparison of gene expression by fibroblast subtypes
Supplementary Figure 17. UMAP plots of T cell-related marker genes
Supplementary Figure 18. Comparison of gene expression by CAR T cell subtypes
Supplementary Figure 19. Further characterization of gene expression by CAR T cells
Supplementary Figure 20. Heatmap of differentially expressed genes in control group
Supplementary Figure 21. Further characterization and validation of CAR T cell phenotype
Supplementary Figure 22. Comparison of gene expression by activated CAR T cells
Supplementary Figure 23. Flow cytometric analysis of CCR2 expression
Supplementary Figure 24. Ligand-receptor interaction analysis
Supplementary Figure 25. Effects of LAF237 on tumor growth during CAR T infusion
Supplementary Figure 26. Effects of LAF237 alone on tumor growth
Supplementary Figure 27. Effect of blocking CXCR3 on the activity of CAR T cells
Supplementary Figure 28. Measurement of cleaved and intact forms of CXCL10
Supplementary Figure 29. Heatmap overview of significantly changed metabolites
Supplementary Figure 30. Heatmap of top 50 metabolites
Supplementary Figure 31. Metabolite sets enrichment analysis
Supplementary Figure 32. Identification of predictive biomarkers
Supplementary Figure 33. Levels of additional predictive biomarkers

**Supplementary Table 1 to 3**

Supplementary Table 1. List of significant interactions in CellPhoneDB analysis

Supplementary Table 2. Peak intensities of metabolites
Supplementary Table 3. Summary of MSEA and pathway impact analysis

**Captions for Movies 1 to 11**

Movie 1. 3D reconstruction of blood vessels wrapping around a tumor transplant in the device
Movie 2. 3D reconstruction of blood vessels penetrating a tumor transplant in the device
Movie 3. Visualization of the perfusability of vascularized human lung tumor explants using 75-kDa FITC-Dextran
Movie 4. Visualization of the perfusability of vascularized human lung tumor explants using 1-µm fluorescent microbeads
Movie 5. Visualization of the perfusability of vascularized lung tumor spheroids using 1-µm fluorescent microbeads
Movie 6. Comparison of initial CAR T cell infusion between the meso-tumor and control groups
Movie 7. Extravasation and directional migration of CAR T cells in the meso-tumor model
Movie 8. Behavior of CAR T cells in the control tumor group
Movie 9. Time-lapse imaging of CAR T cell-infused lung tumor growth without LAF237
Movie 10. Time-lapse imaging of CAR T cell-infused lung tumor growth treated with 50 nM of LAF237
Movie 11. Time-lapse imaging of CAR T cell-infused lung tumor growth treated with 1000 nM of LAF237

### Supplementary Figure 1. Fabrication of microdevice for in vitro tumor transplantation

**a**. The open-top microdevice consists of three layers including a three-lane culture chamber, an open-top ceiling, and an insert. **b**. To fabricate the device, degassed PDMS prepolymer is dispensed onto 3D printed molds and cured at 65ºC for 2 hours (Step 1). The fully cured PDMS slabs are then peeled off the molds (Step 2) and assembled to generate an open-top device in which the protruding features of the insert are fit into the open wells of the device ceiling (Step 3).

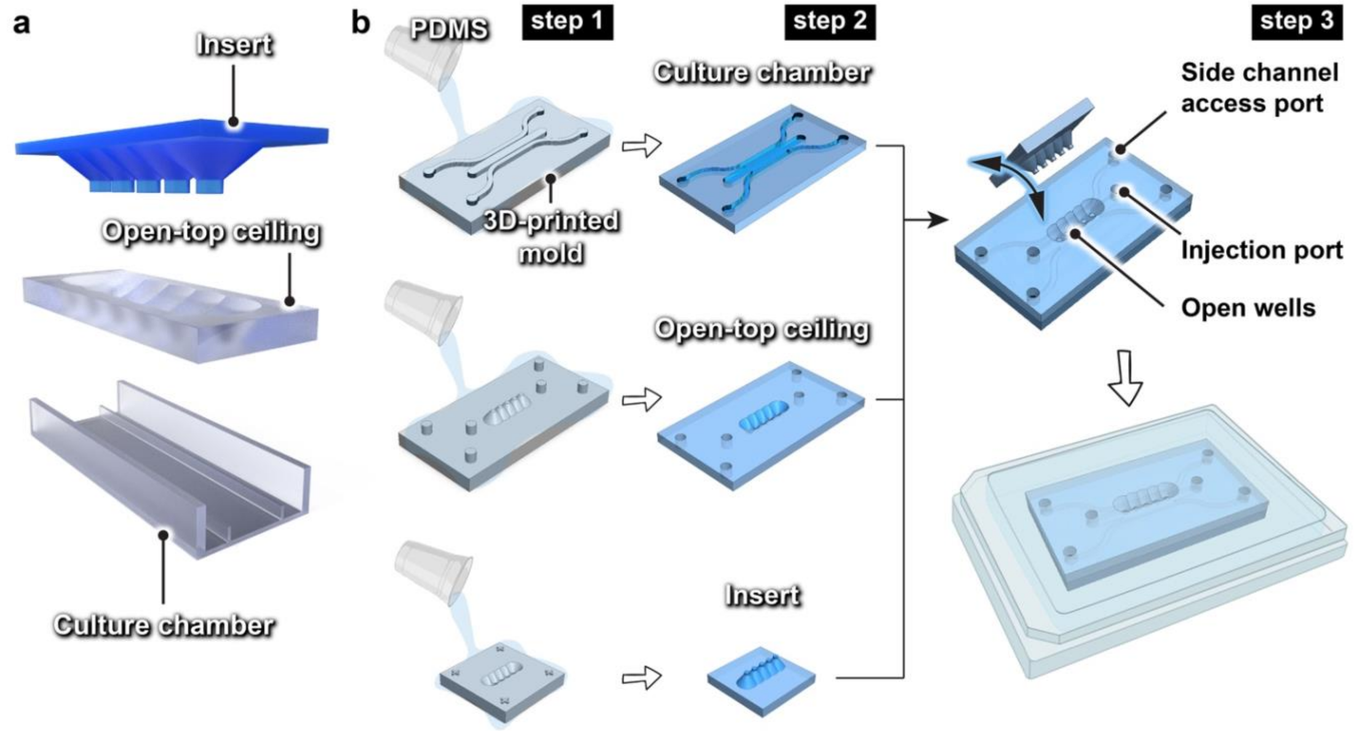

### Supplementary Figure 2. Injection of cell-hydrogel mixture

Time-lapse images demonstrating the injection of cell-hydrogel mixture into the middle chamber of the device. Injection occurs while the open wells in the ceiling of the device are covered by the insert (shown as a square at the center). For visualization purposes, the mixture solution was dyed black in these images.

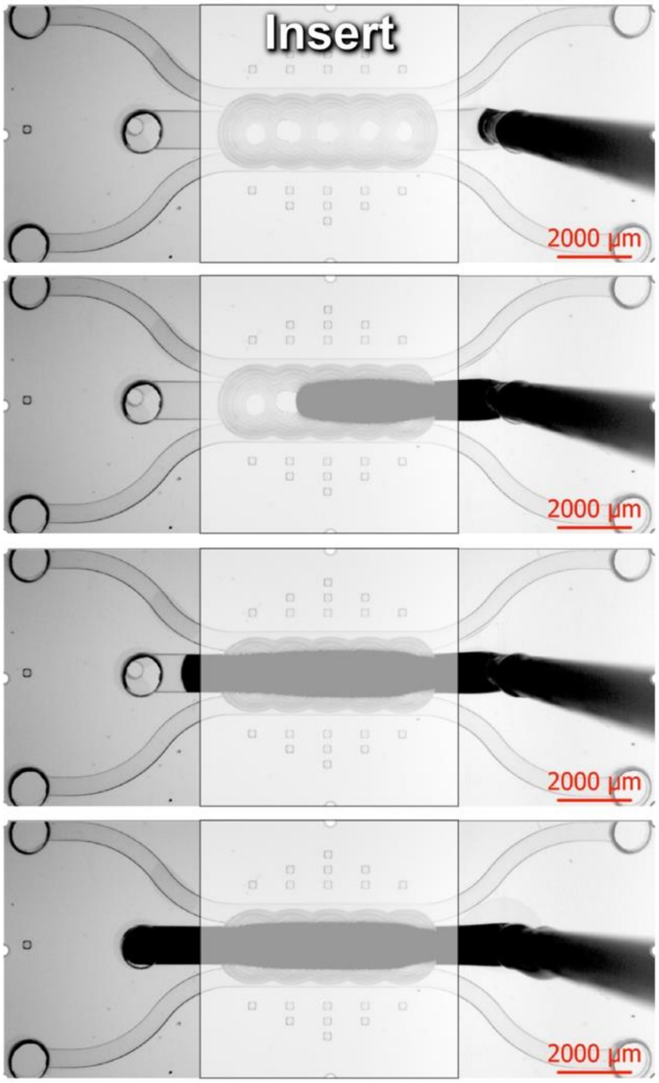

### Supplementary Figure 3. Effect of initial tumor size on tumor vascularization

**a,** Representative confocal micrographs showing vascularization of single lung tumors with different initial sizes in our device. Scale bars, 200 μm (left column) and 400 μm (right column). **b-d,** Quantification and comparison of (**b**) average vessel diameter, (**c**) the number of vessel junctions, and (**d**) the percentage of vessel-covered area. Data are presented as mean ± SEM (n = 3-14).

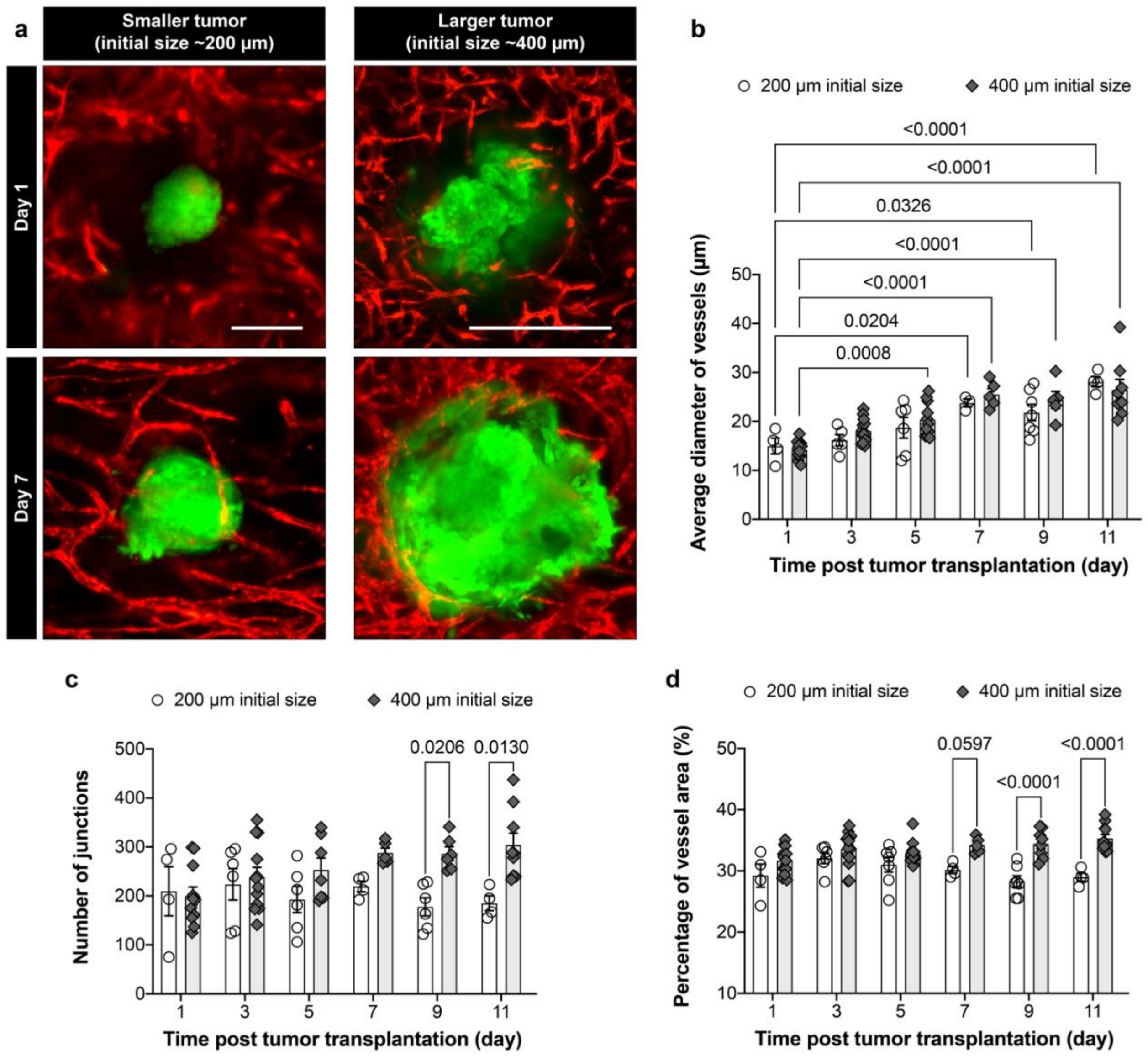

### Supplementary Figure 4. Tumor nests

Histological sections of squamous cell carcinoma in human lung cancer (**left**) and head and neck cancer (**right**). Cancer lesions are seen with microscopic tumor nests in the vascularized stroma.

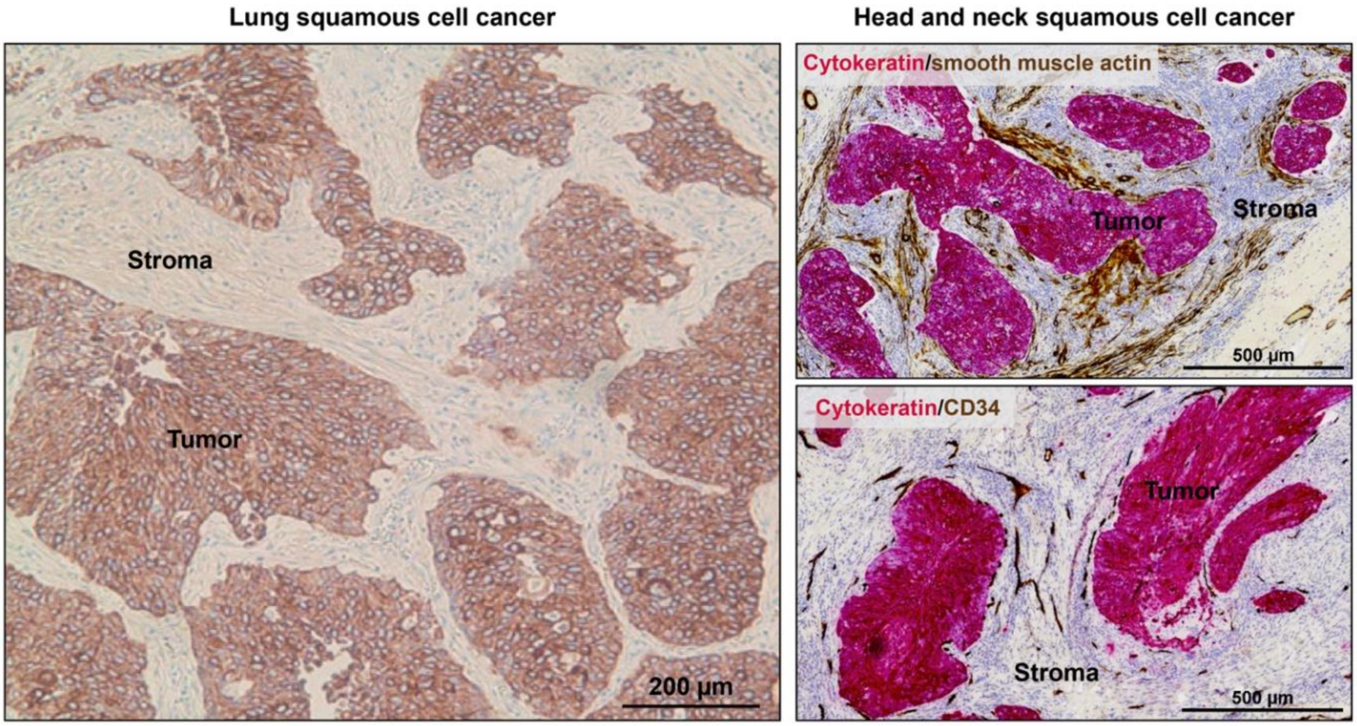

### Supplementary Figure 5. In vitro transplantation of tumor spheroids and organoids

**a,b,** Fluorescence images of lung tumor spheroids composed of A549 human lung adenocarcinoma cells (**a**) and human colorectal cancer organoids (**b**) embedded and vascularized in the in vitro transplantation microdevice.

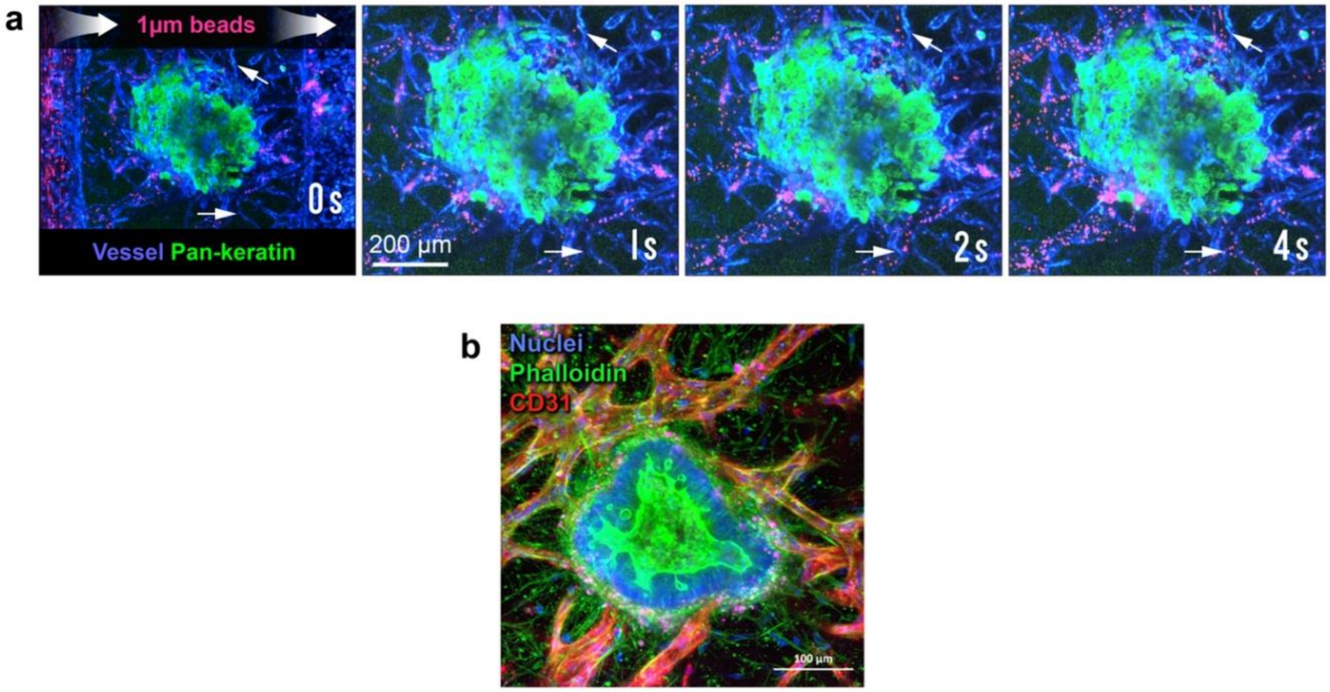

### Supplementary Figure 6. Overexpression of mesothelin in A549 and meso-CAR design

**a-c**, Comparison of wild-type A549 cells (**Ctrl-A549**) and those transduced to overexpress mesothelin (**Meso-A549**). **a**, Fluorescence images show mesothelin expression (red) by GFP-expressing A549 cells cultured in 2D monolayers (top row) and in A549-CDX tumors (bottom row). **b**, Quantification of mesothelin expression. Data are presented as mean ± SEM ($n \geq 6$). **c**, Expression of mesothelin and GFP by both A549 cell lines was further confirmed and quantified by flow cytometry. **d**, Schematic representation of the mesothelin-binding chimeric receptors. The SS1 scFv domain is designed to recognize human mesothelin.

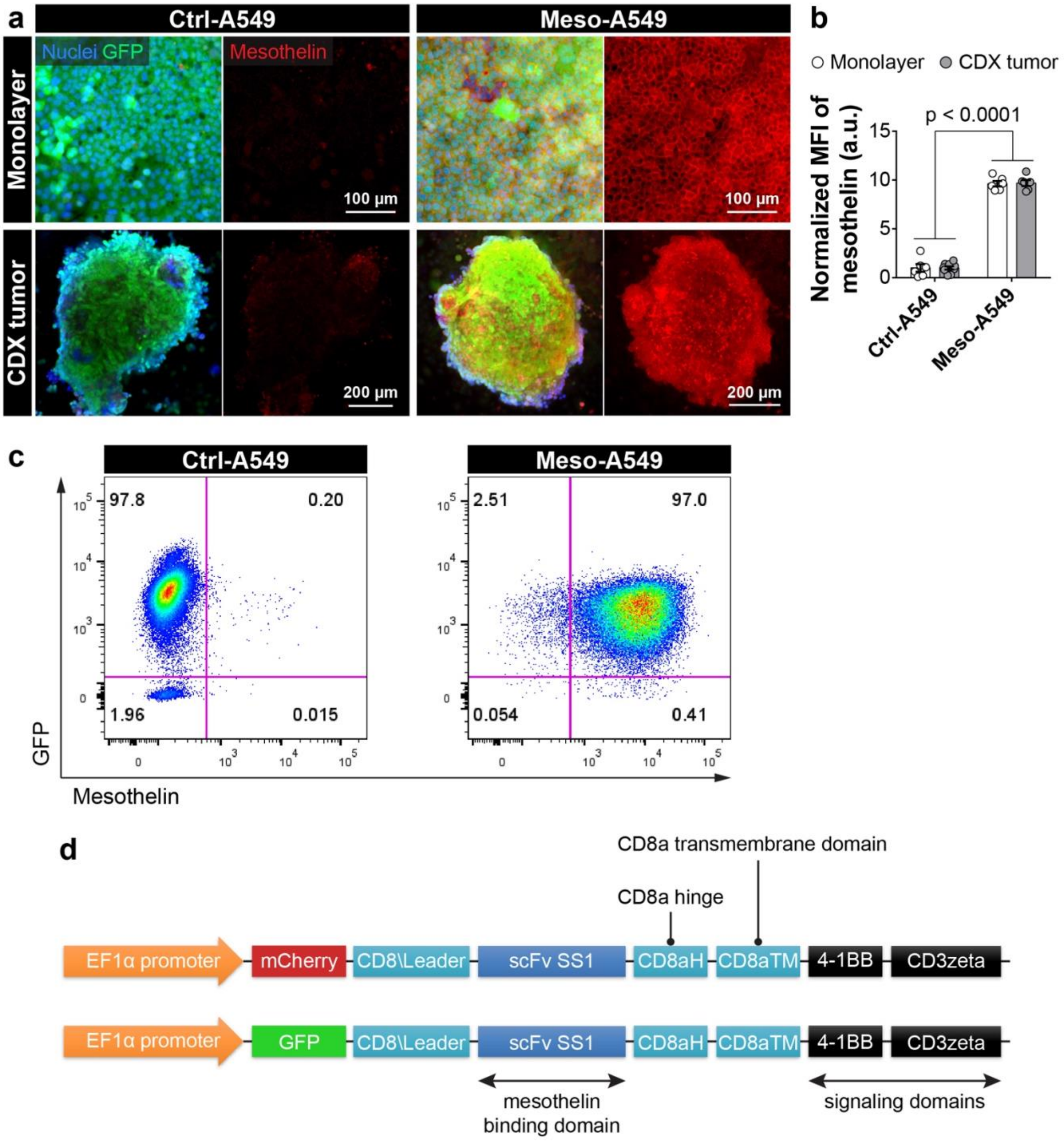

### Supplementary Figure 7. Growth of lung tumors and their responses to CAR T cells

Fluorescence time-lapse images of meso- (left) and control (right) tumors in the device prior to (days 1 to 11) and after (days 14 to 19) meso-CAR T cell infusion. Compared to the control group, the post-infusion meso-tumor model is seen with noticeably higher levels of CAR T cell adhesion and retention in tumor masses. Note that meso-CAR T cell trafficking in areas distant to the tumors is also greater in the meso-tumor model than in the control group, which is consistent with the difference in endothelial expression of ICAM-1 between the two models shown in **Figure 2p**. Scale bars, 500 μm.

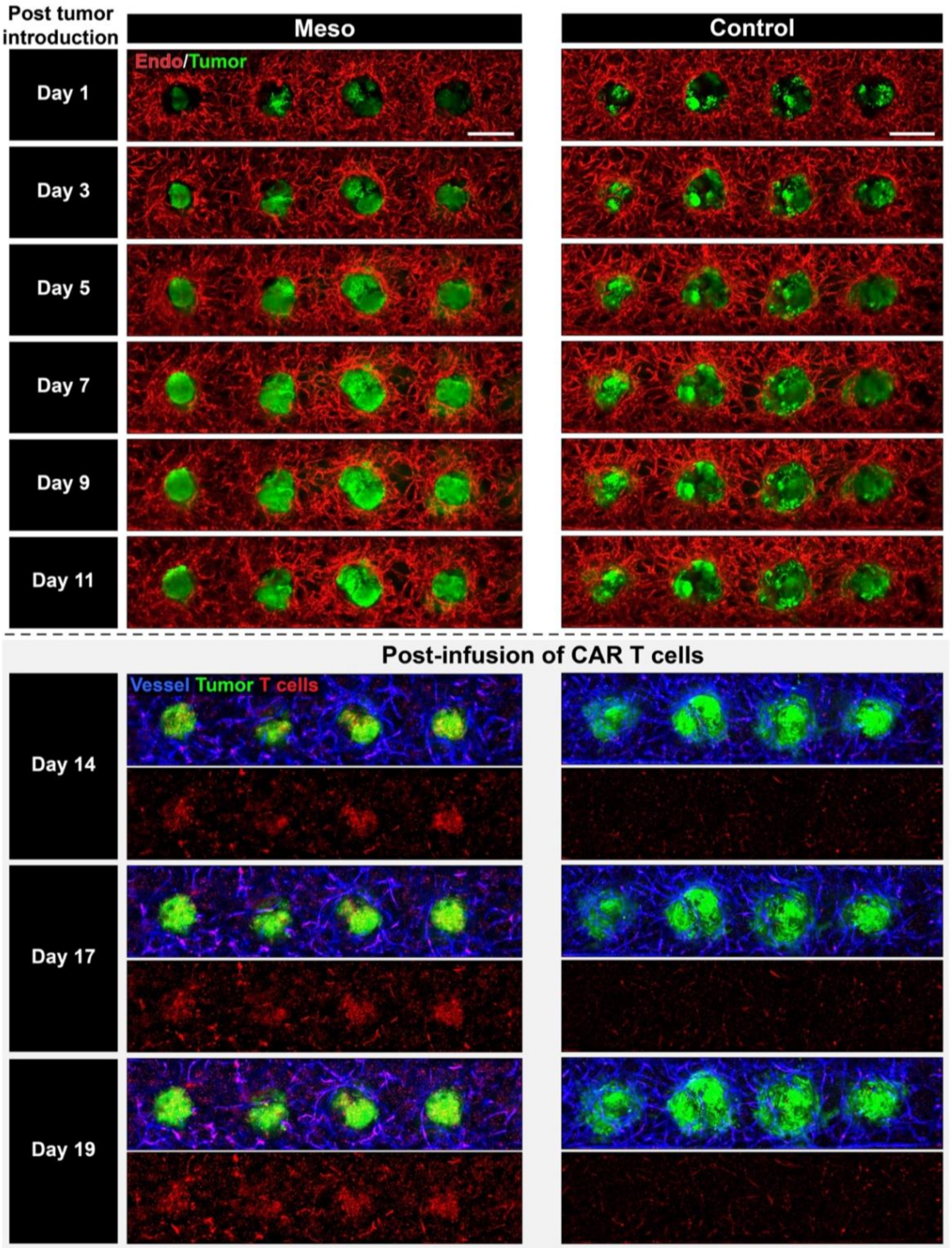

### Supplementary Figure 8. Analysis of lung tumors infused with non-transduced T cells

**a,b,** Representative confocal micrographs of single meso-tumor constructs treated with non-transduced (NTD) T cells (**a**) or meso-CAR T cells (**b**). Scale bars, 250 µm. **c,d,** Quantification of T cell area (**c**) and normalized tumor area (**d**) over time. Data are presented as mean ± SEM (n ≥ 3). CAR T cells derived from one healthy donor were tested.

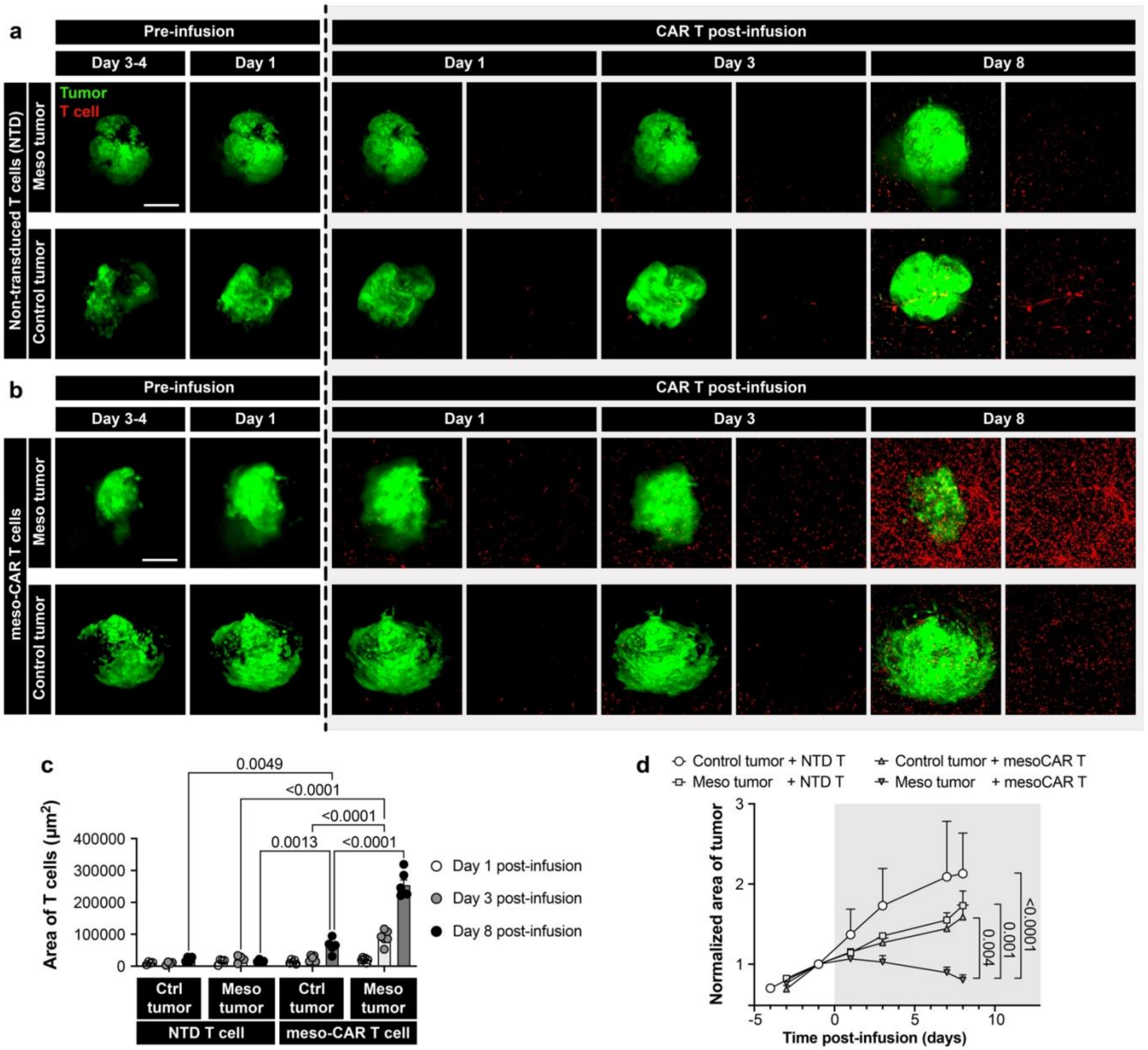

### Supplementary Figure 9. Analysis of chemokine production

**a**, Array screening of 38 chemokines using device effluent samples collected from microengineered constructs containing blood vessels only (vessel only control) (**b**), vascularized control tumors (Control tumor model) (**c**), and vascularized meso-tumors (Meso-tumor model) (**d**). **e**, Quantification and comparison of the level of chemokine expression.

**a**

| | A | B | C | D | E | F | G | H | I | J | K | L | M |
|---|---|---|---|---|---|---|---|---|---|---|---|---|---|
| 1 | POS1 | POS2 | POS3 | NEG | NEG | BLC | CCL28 | Ckb8-1 | CTACK | CXCL16 | ENA-78 | Eotaxin | Eotaxin-2 |
| 2 | POS1 | POS2 | POS3 | NEG | NEG | BLC | CCL28 | Ckb8-1 | CTACK | CXCL16 | ENA-78 | Eotaxin | Eotaxin-2 |
| 3 | Eotaxin-3 | Fractalkine | GCP-2 | GRO | GROa | HCC-4 | I-309 | I-TAC | IL-8 | IP-10 | Lymphotactin | MCP-1 | MCP-2 |
| 4 | Eotaxin-3 | Fractalkine | GCP-2 | GRO | GROa | HCC-4 | I-309 | I-TAC | IL-8 | IP-10 | Lymphotactin | MCP-1 | MCP-2 |
| 5 | MCP-3 | MCP-4 | MDC | MIG | MIP-1a | MIP-1b | MIP-1d | MIP-3a | MIP-3b | MPIF-1 | NAP 2 | PARC | RANTES |
| 6 | MCP-3 | MCP-4 | MDC | MIG | MIP-1a | MIP-1b | MIP-1d | MIP-3a | MIP-3b | MPIF-1 | NAP 2 | PARC | RANTES |
| 7 | SDF-1a | SDF-1b | TARC | TECK | NEG | NEG | NEG | NEG | NEG | NEG | NEG | NEG | NEG |
| 8 | SDF-1a | SDF-1b | TARC | TECK | NEG | NEG | NEG | NEG | NEG | NEG | NEG | NEG | NEG |

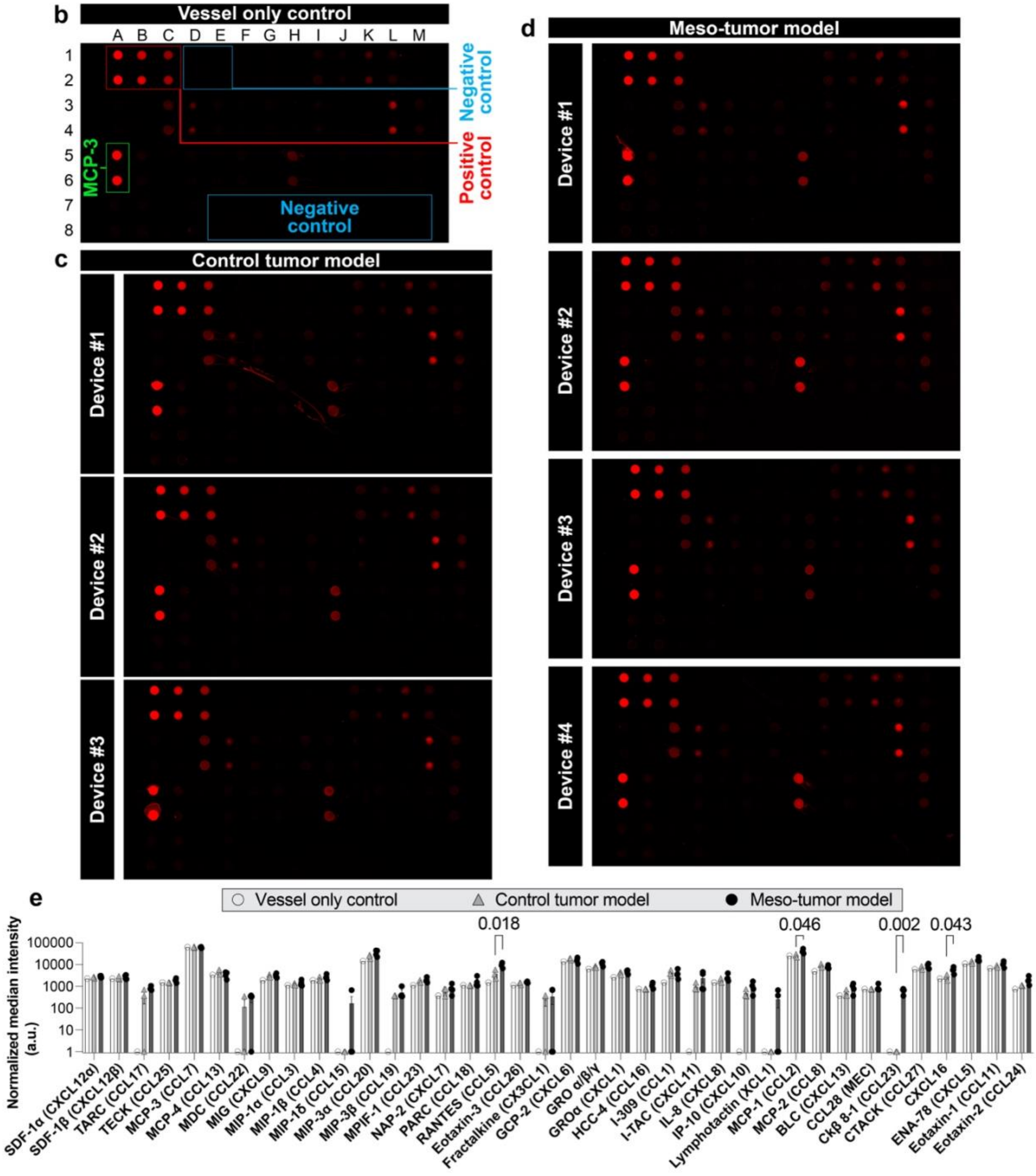

### Supplementary Figure 10. Histological analysis of meso- and control tumors

Representative images showing histological sections and H&E staining of CAR T cell-infused meso- and control tumors. The green dashed lines show the outline of tumors. Tissue sections were cut from similar locations of respective tissue blocks. After device delamination, more than 90% of the tumors remained intact in the vascularized constructs. As shown in the images below, the retrieved tumors in both groups retained their shape and overall tissue structure. Scale bars, 100 μm.

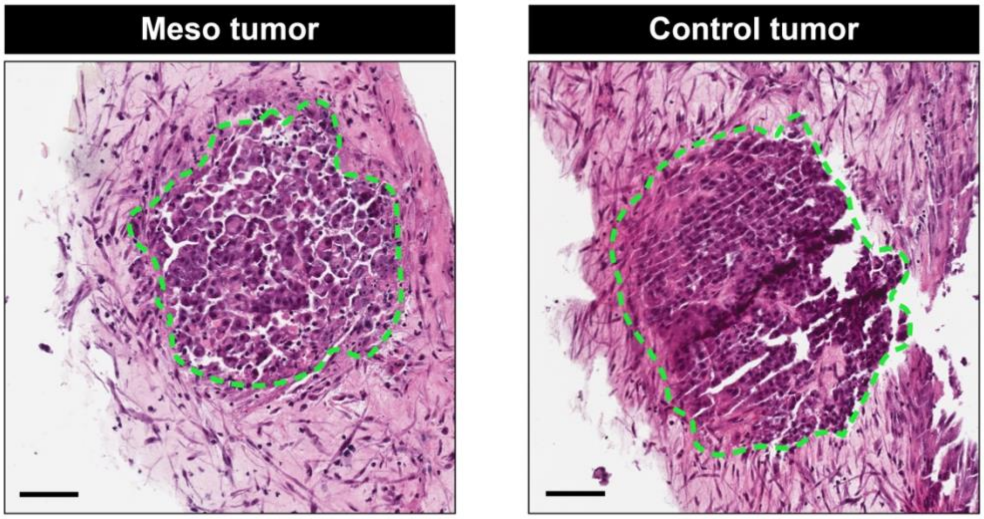

### Supplementary Figure 11. Flow cytometric analysis of CAR T cell phenotype

**a**, Gating strategy of flow cytometric analysis. Flow cytometry is applied to the single cell suspensions derived from our model to identify the phenotypic states and changes of CAR T cells post-infusion. The live singlet lymphocytes are first gated for CD3+ CD8+ T cells, and then gated based on the CAR expression to identify meso-CAR T cells. Within meso-CAR T cells, we further gate using CD45RO and CD62L as markers for central memory cells (CD45RO+ CD62L+) and effector memory cells (CD45RO+ CD62L-); using CD103 as positive marker for tissue resident cells; using CD69 as positive marker for activated and tissue resident cells; using PD-1 as marker for recently activated cells. **b,c,d,** Representative flow cytometry plots and quantification of surface marker expression by CAR T cells, including CD69 (**b**), CD45RO and CD62L (**c**), and CD103 (**d**). Data are presented as mean ± SEM. *P < 0.05, **P < 0.01, and ***P < 0.001. (n = 3-4).

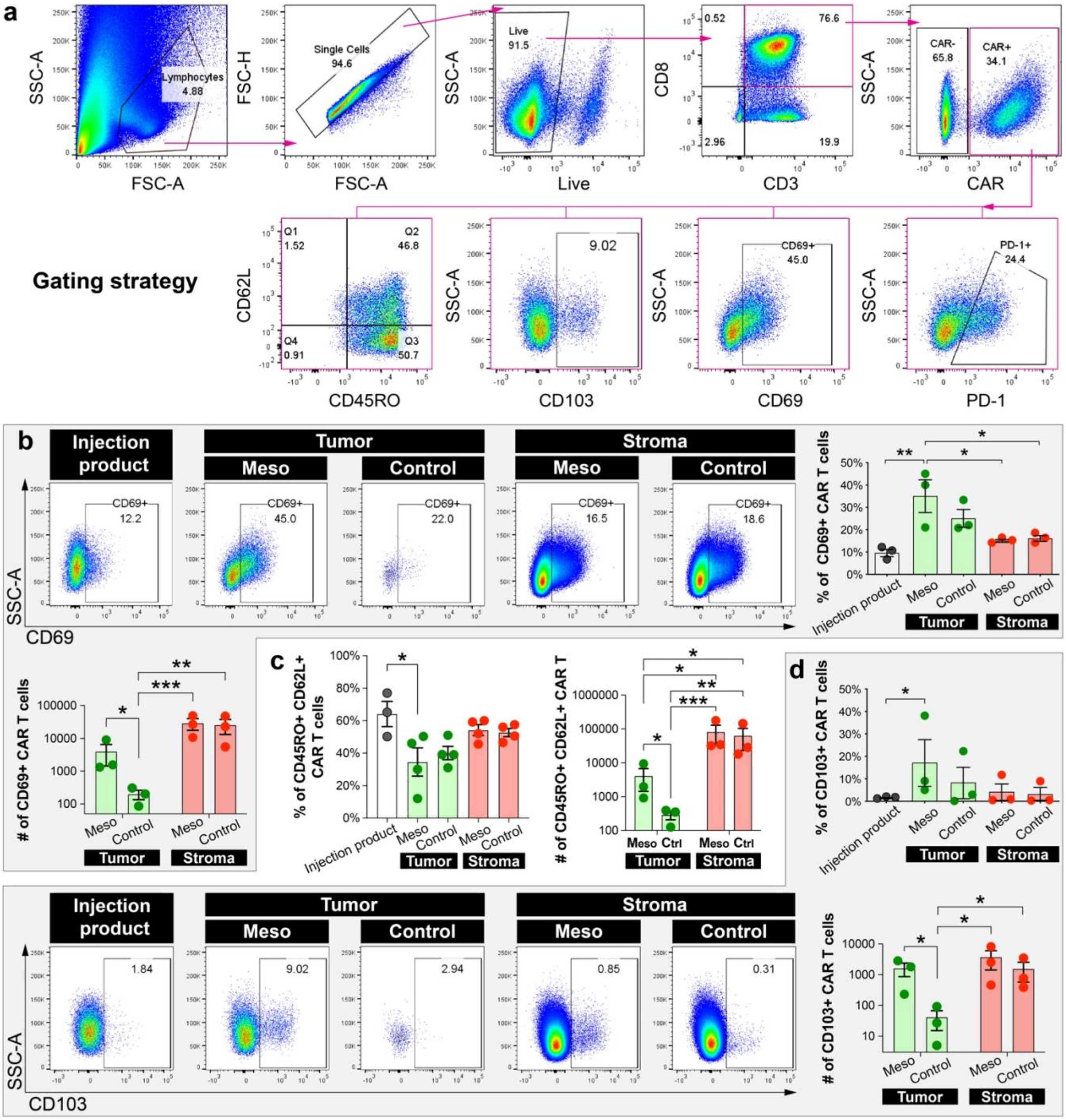

### Supplementary Figure 12a. Identification of UMAP clusters

**a,** UMAP projection of cell populations in the CAR T cell-treated meso-tumor model at Day 6 post-infusion. Endo represents endothelial cells. **b,** UMAP plots of cell type-specific markers. **c,** UMAP clusters annotated with their location within the model.

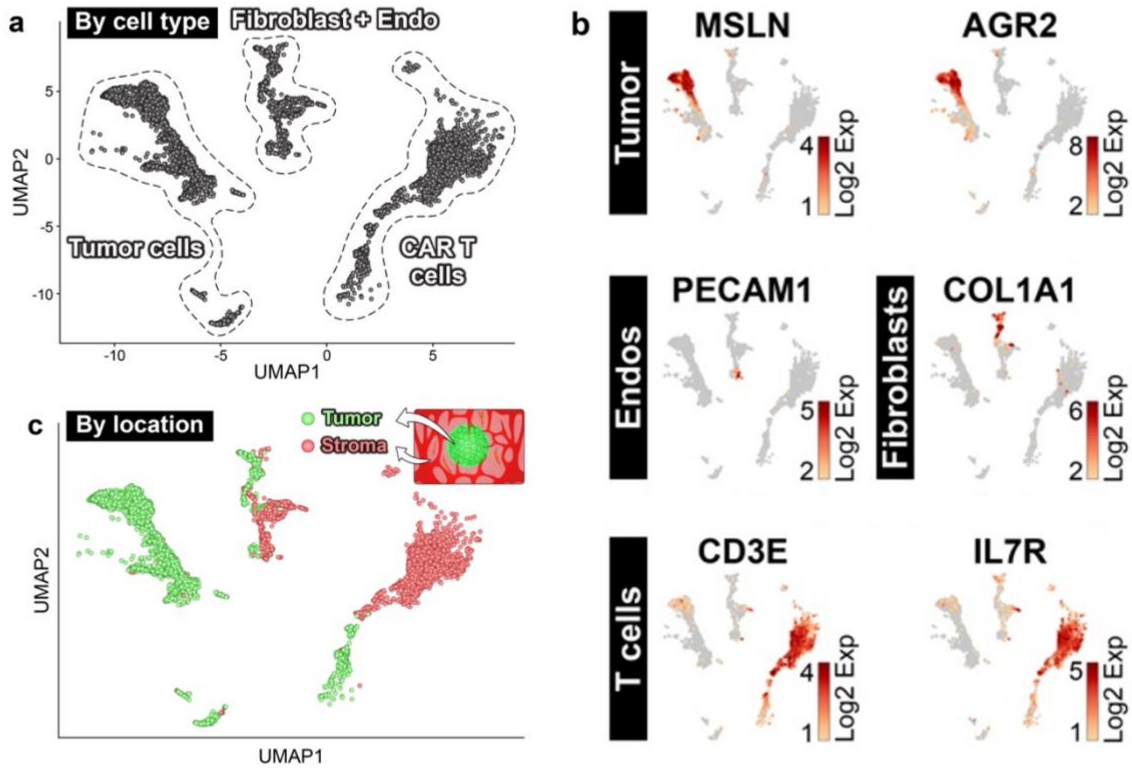

### Supplementary Figure 12b. Heatmap of differentially expressed genes in meso-tumor group

Heatmap showing top 20 differentially expressed genes for each cluster in the meso-tumor group.

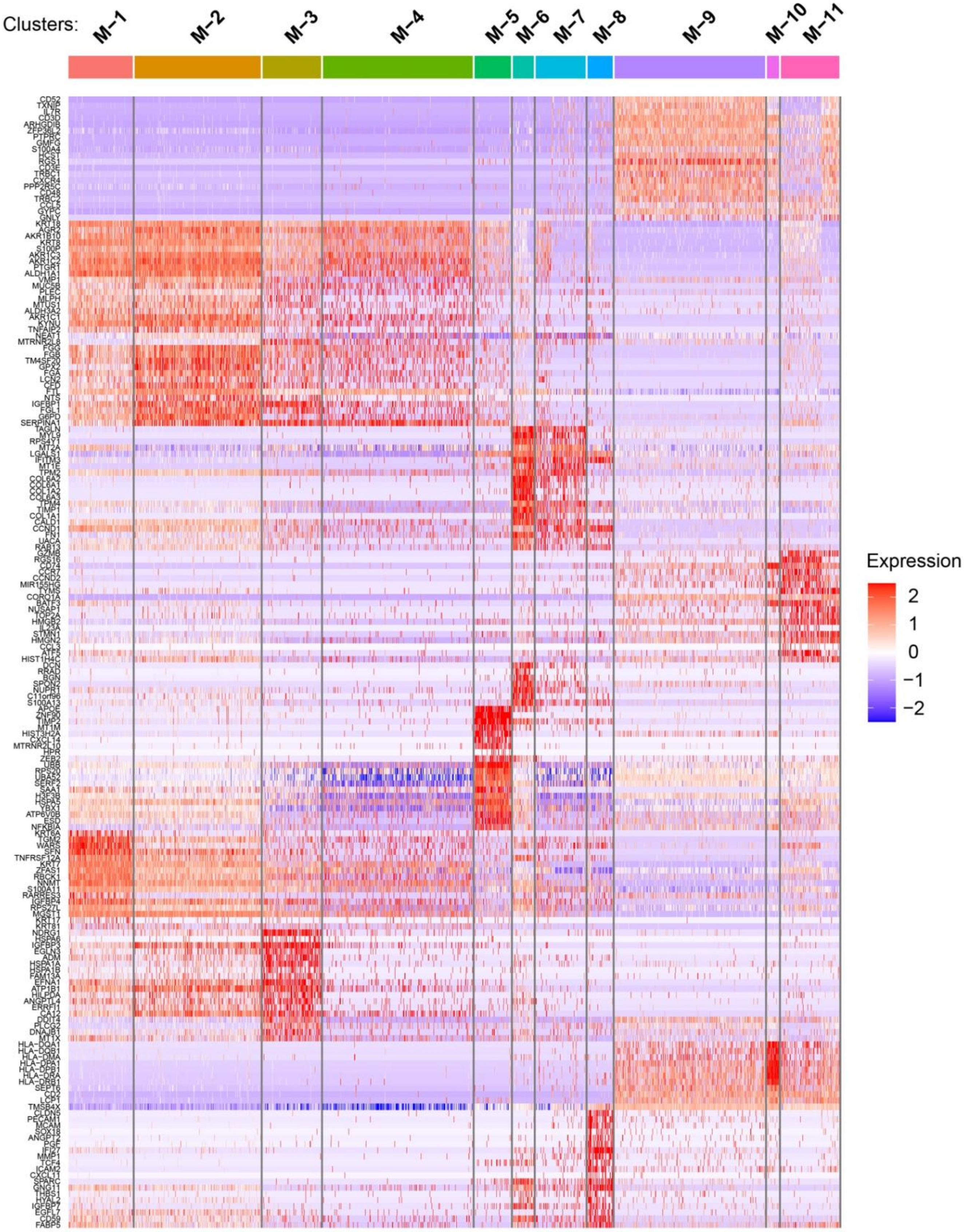

### Supplementary Figure 13. Gene Ontology (GO) enrichment analysis

GO enrichment analysis for each cluster in the meso-tumor group. A Fisher's exact test was performed and corrected by the calculation of False Discovery Rate (FDR), with the enrichment cutoff at FDR $p < 0.05$.

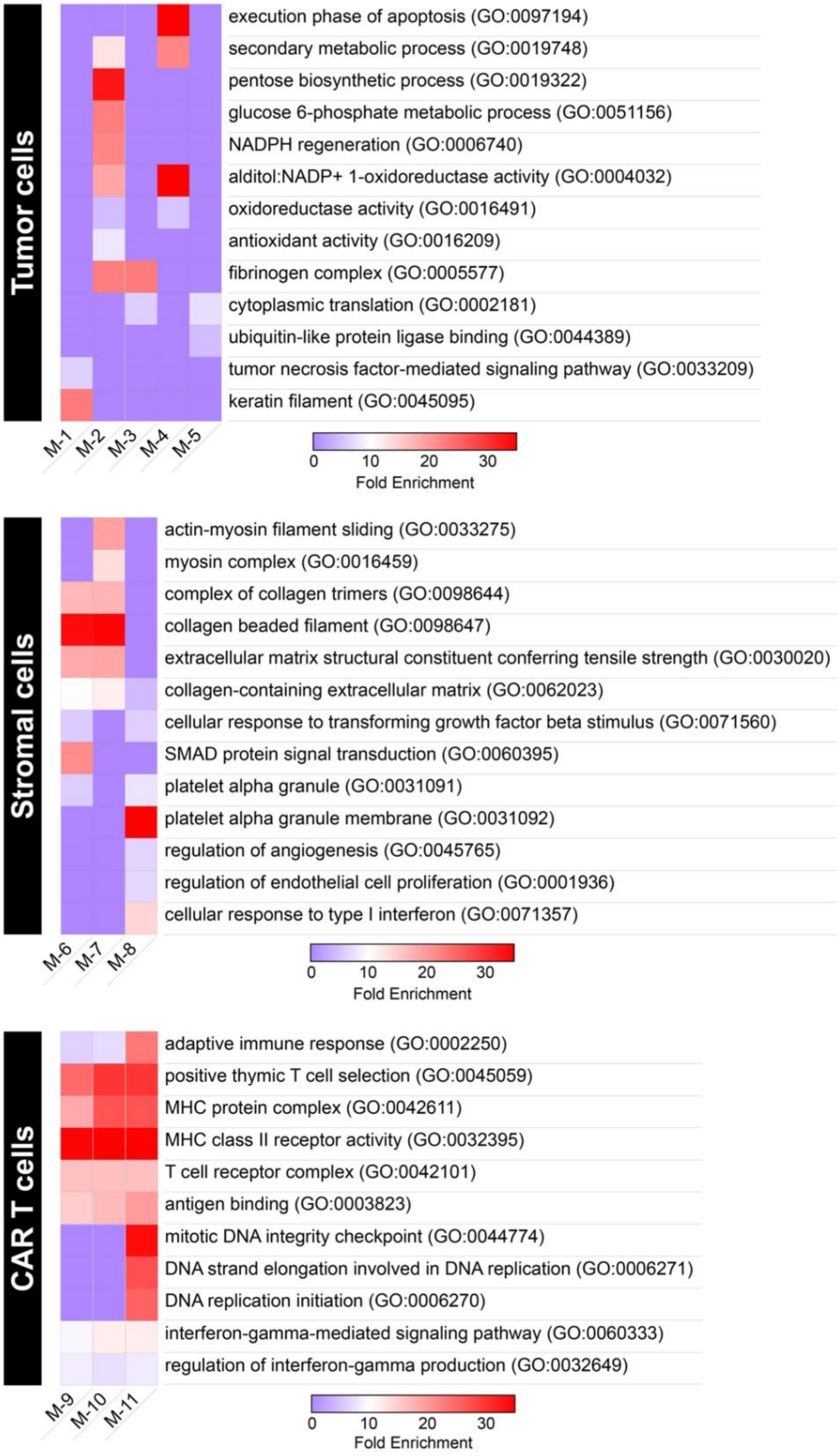

### Supplementary Figure 14. Expression of cluster-specific marker genes

Violin plots showing select top differentially expressed genes by individual clusters identified in the meso-tumor model.

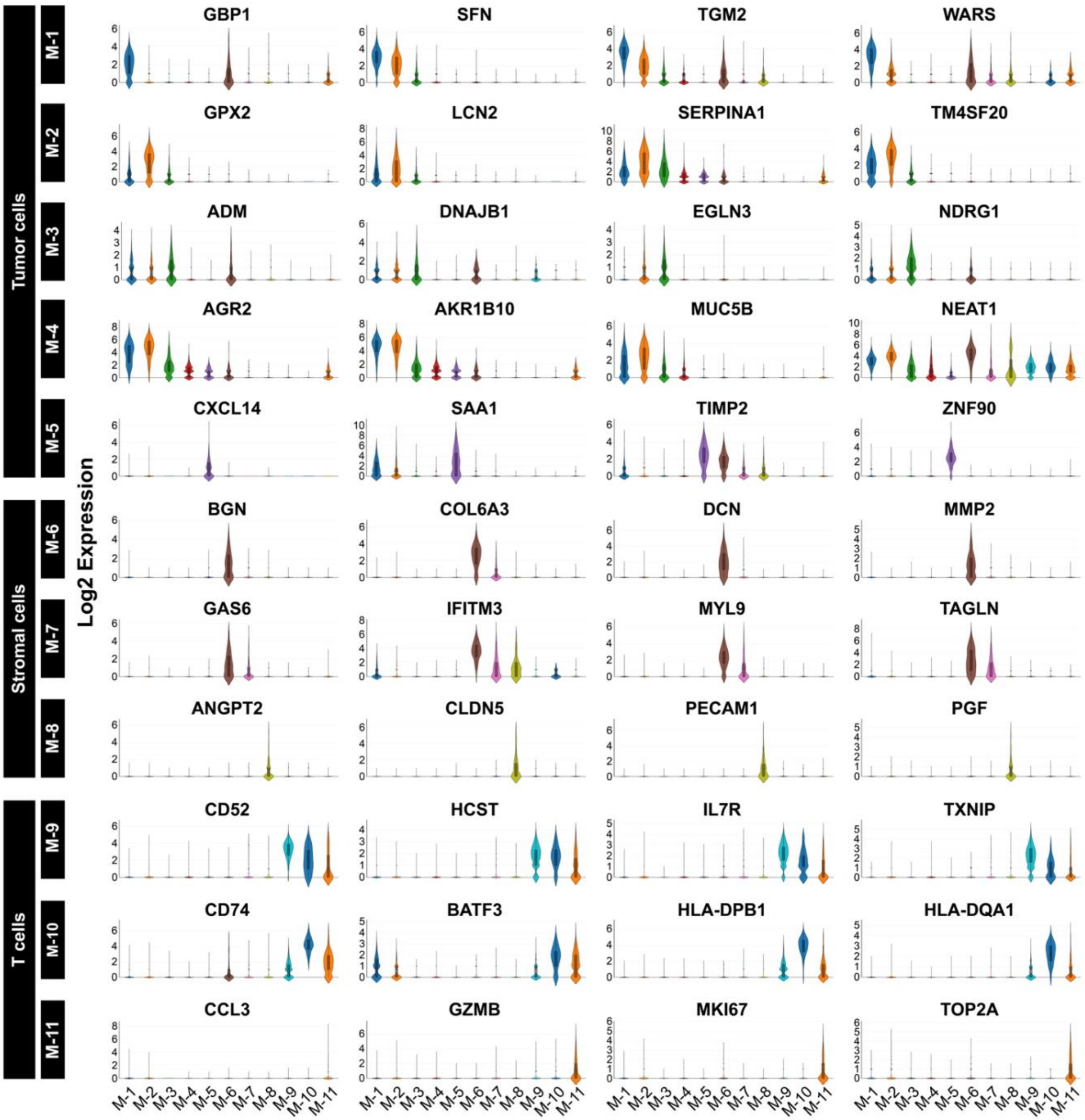

### Supplementary Figure 15. UMAP plots of tumor and stromal cell-related marker genes

UMAP plots showing the expression levels and spatial distribution of select genes for tumor and stromal cells in the meso-tumor group.

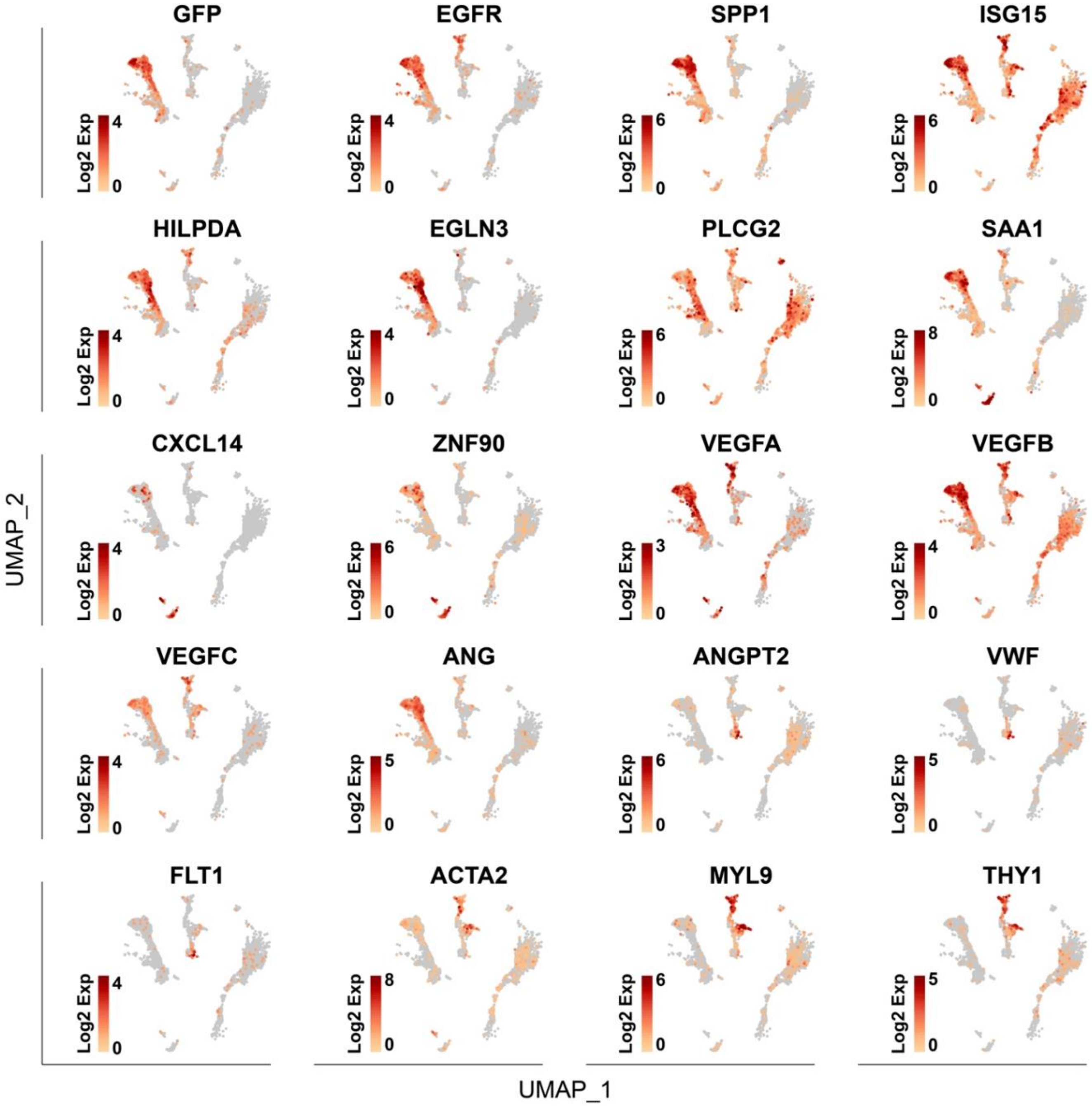

### Supplementary Figure 16. Comparison of gene expression by fibroblast subtypes

Violin plots depicting the expression of select genes of interest in fibroblast subtypes from meso-tumor and control groups. Genes including TGFB1, TGFBR2, SMAD2, FAP, ACTA2, SPARC, COL1A1, and FN1 are significantly upregulated in the subpopulations of fibroblast in which the majority of cells are located in the tumor compartment (i.e., M-6 and C-4) as shown in **Figs. 3e, 3j**. Additional genes associated with fibroblasts in tumors in vivo such as HGF, FGF7, DCN, TWIST1, and LTBP1 are also upregulated in M-6 and C-4.

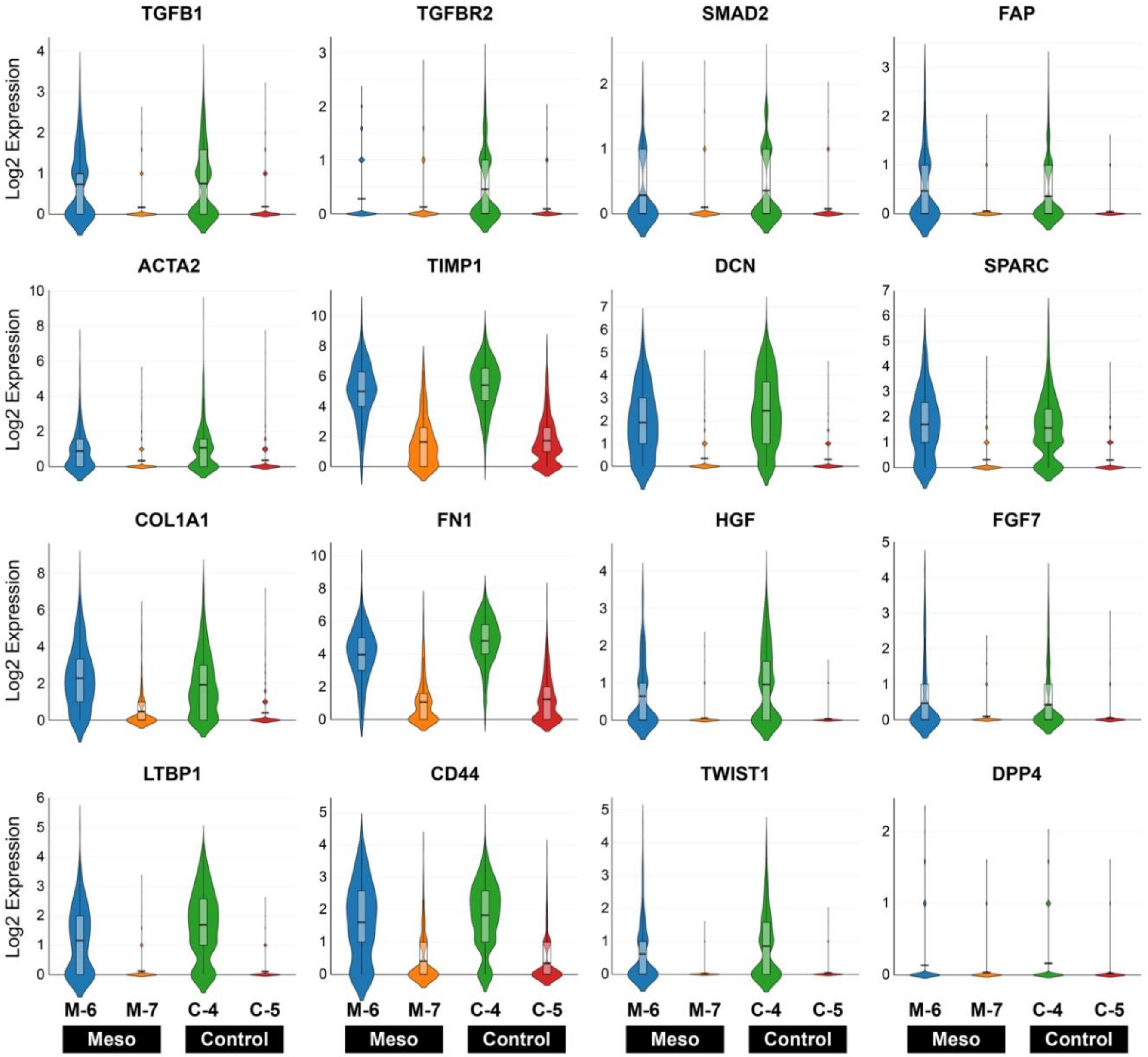

### Supplementary Figure 17. UMAP plots of T cell-related marker genes

UMAP plots showing the expression level and spatial distribution of select T cell genes.

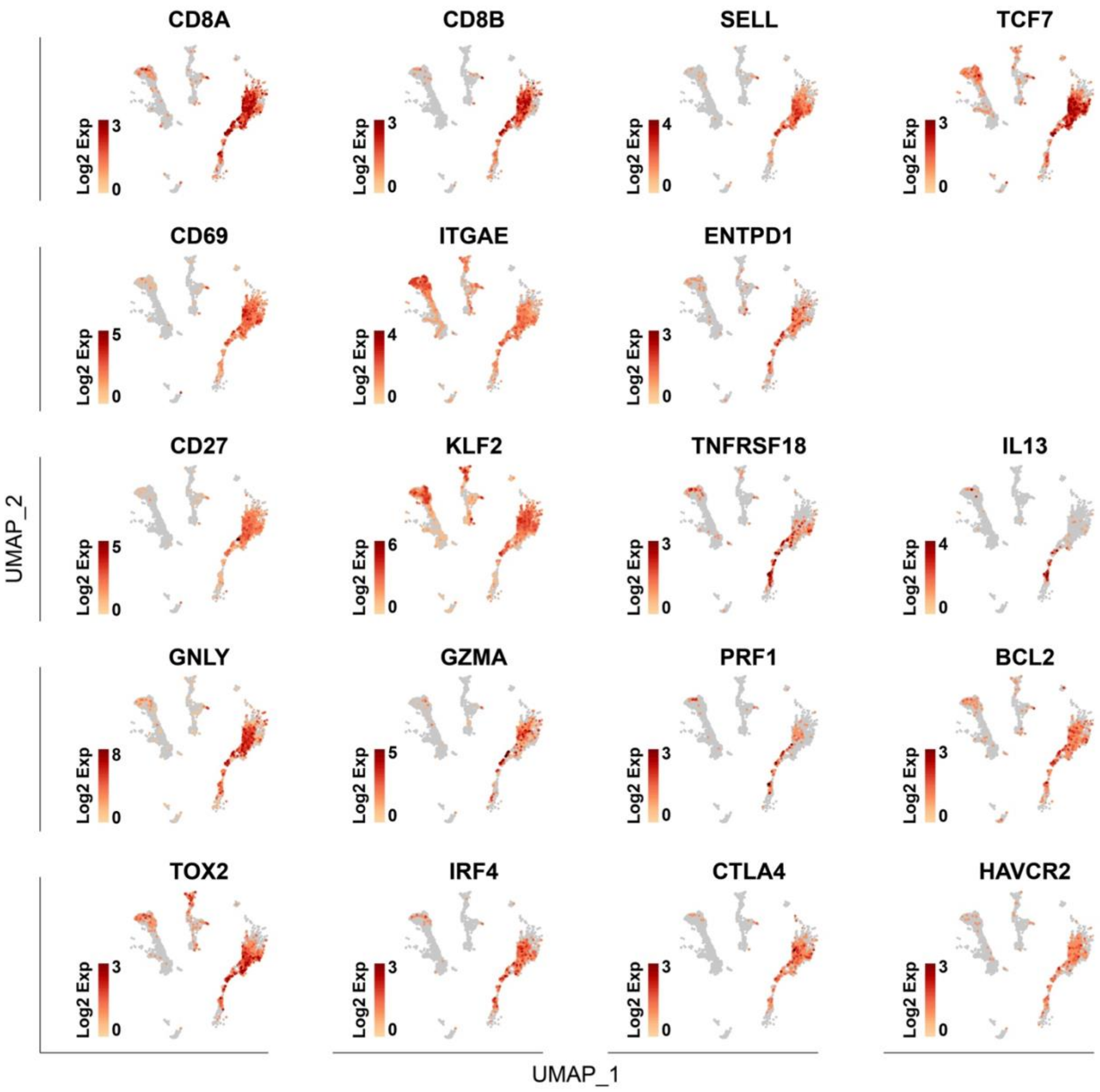

### Supplementary Figure 18. Comparison of gene expression by CAR T cell subtypes

Violin plots showing the expression of gene signatures and individual marker genes by the subpopulations of CAR T cells in the meso-tumor group. ****P* < 1e-7 against M-11 for LEF1 and SELL, ****P* < 1e-7 against M-9 for MIR155HG, ****P* < 1e-7 against other clusters for the rest of genes.

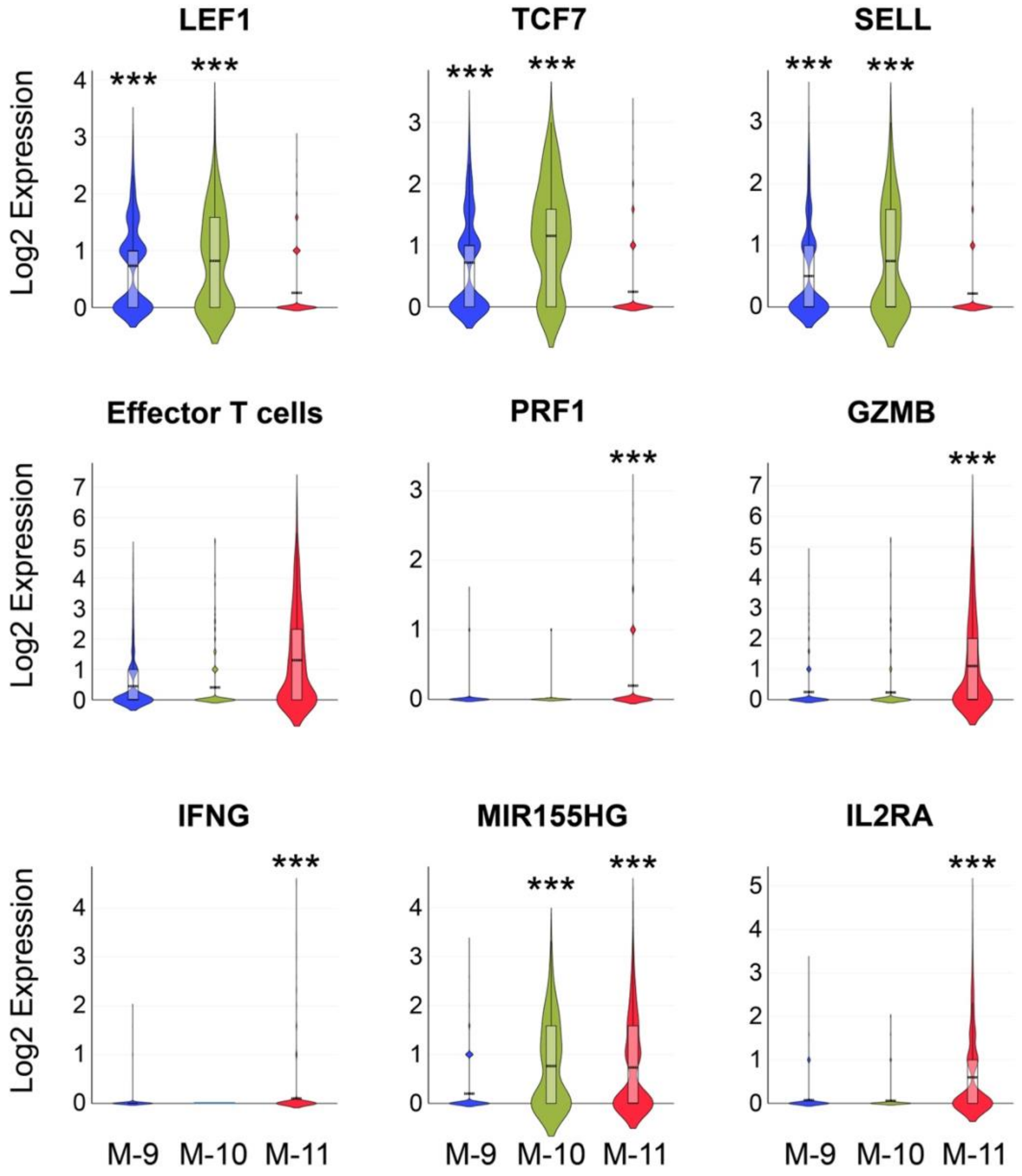

### Supplementary Figure 19. Further characterization of gene expression by CAR T cells

UMAP plots showing the expression level and spatial distribution of select genes in CAR T cells associated with 4-1BB signaling (left) and T cell activation (right).

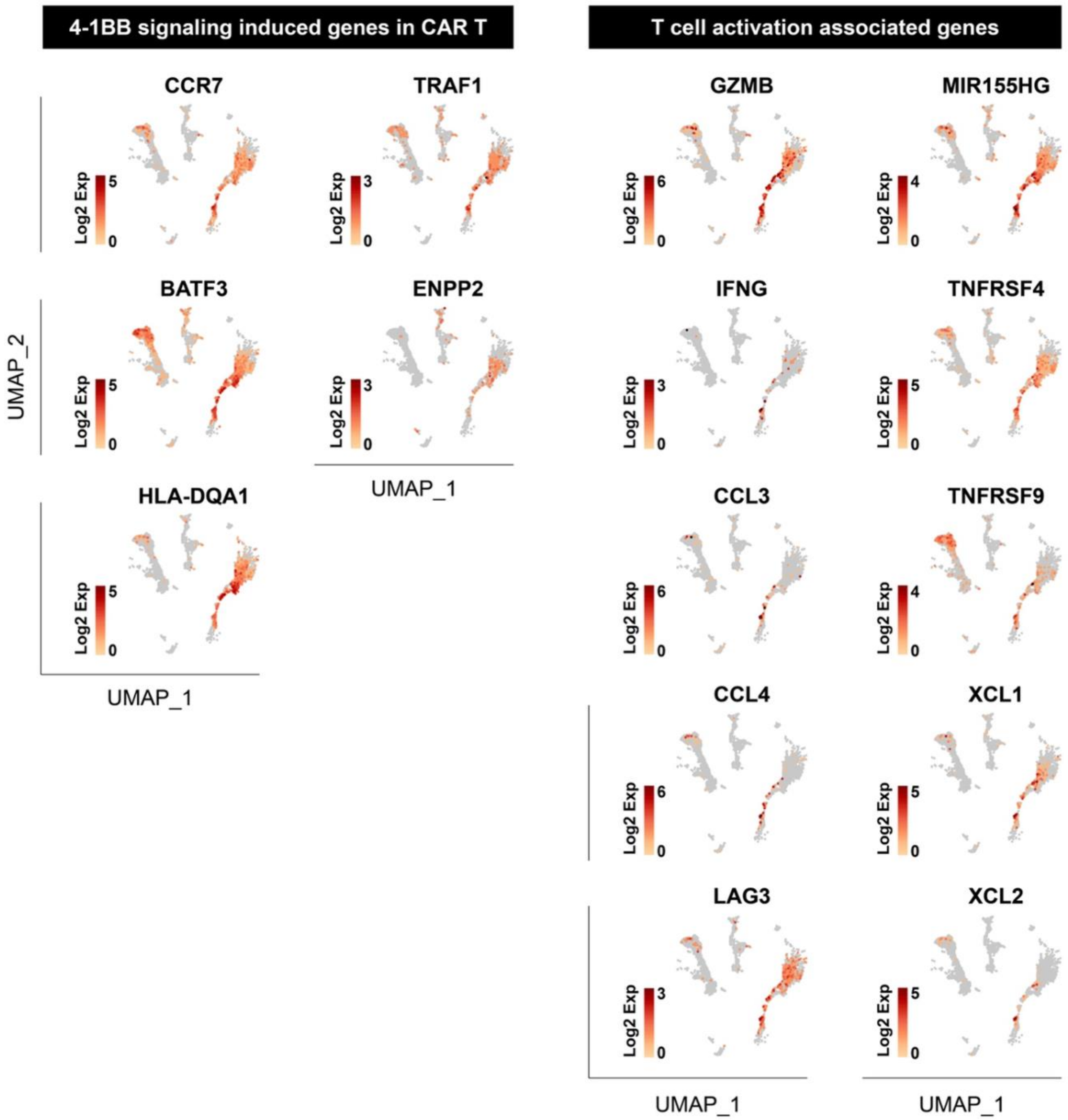

### Supplementary Figure 20. Heatmap of differentially expressed genes in control group

Heatmap showing top 20 differentially expressed genes for each cluster in the control group.

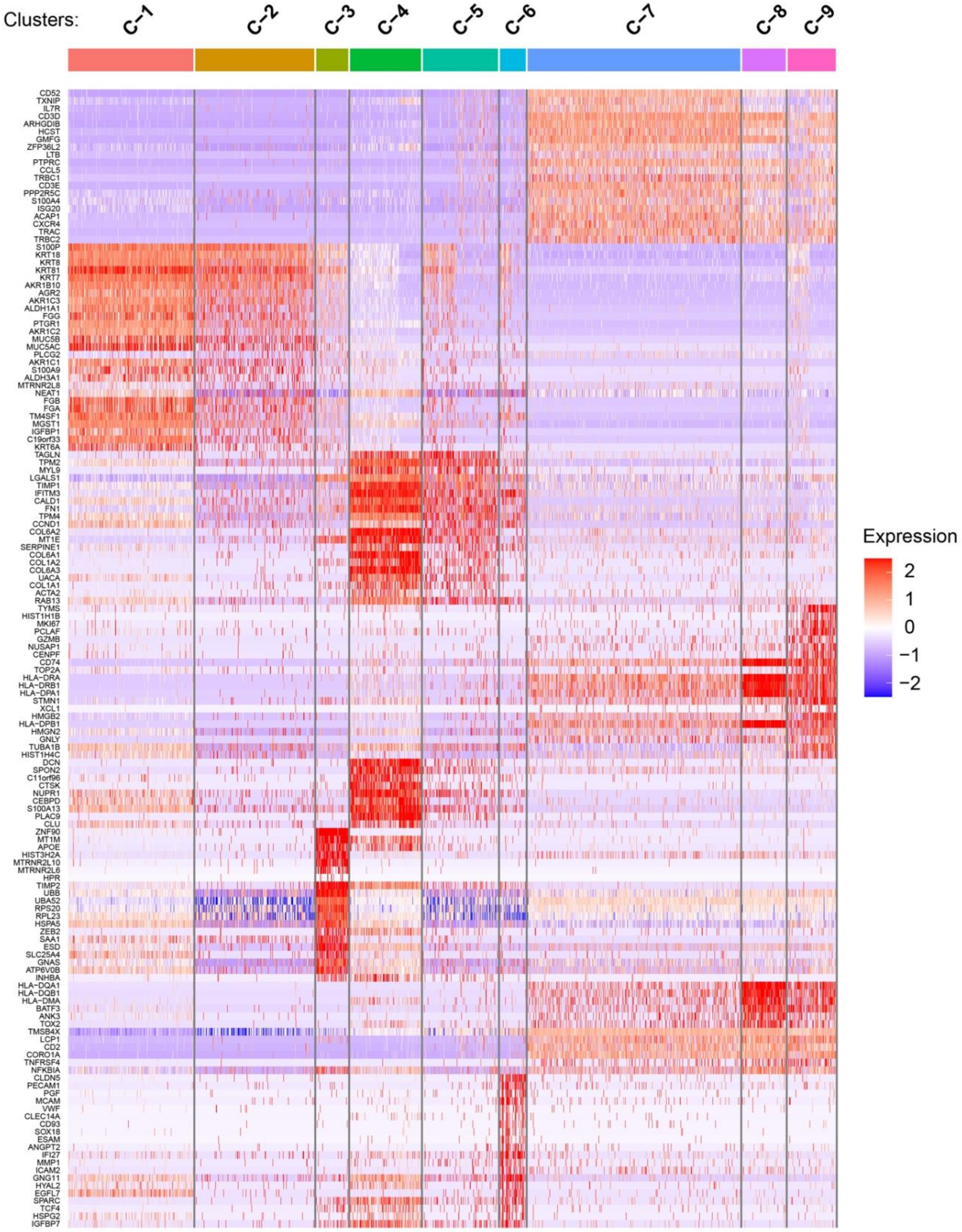

### Supplementary Figure 21. Further characterization and validation of CAR T cell phenotype

**a,** Violin plots of select gene signatures expressed by tumor-infiltrating activated effector CAR T cells from the meso-tumor (**M-11**) and control (**C-9**) models. **b,** Heatmap showing SingleR scores for CAR T cells in the meso-tumor model computed against each CD8 T cell subtype described in the GSE99254 reference dataset of human endogenous tumor-infiltrating T cells from NSCLC patients. Each vertical line represents a single cell. **c,** UMAP projection of CAR T cell clusters in the meso-tumor model annotated using labels from the reference human dataset. The results are consistent with what is shown by cell type-specific marker-based manual annotation and characterization (**Fig. 3**). In the meso-tumor model, for example, most of the cells in Cluster **M-9** – a subpopulation of CAR T cells that were less activated and expressed higher levels of naïve markers – were recognized by SingleR as CD8_C1-LEF1, which represents patient naïve T cells mostly from the peripheral blood. In comparison, a large portion of the cells in Clusters **M-10** and **M-11**, especially tumor-infiltrating CAR T cells in **M-11**, were annotated as either CD8_C5-ZNF683 (tissue-resident memory T cells) or CD8_C6-LAYN (exhausted T cells), both of which originate from patient tumors. Very few T cells were identified as CD8_C3-CX3CR1 effector cells or CD8_C4-GZMK "pre-exhausted" T cells, suggesting the lack of terminal effector differentiation and thus potential for persistence of CAR T cells.

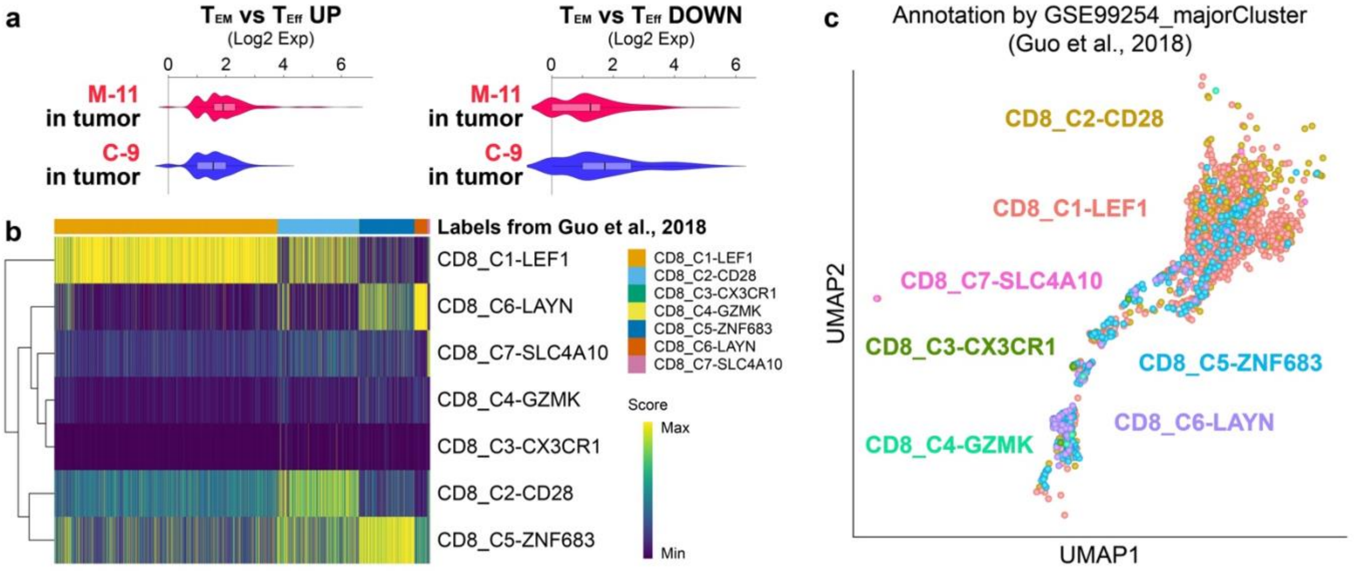

### Supplementary Figure 22. Comparison of gene expression by activated CAR T cells

**a,** Violin plots comparing the expression of select genes by activated effector CAR T cells in the stromal and tumor compartments in the meso-tumor (**M-11**) and control (**C-9**) groups. **b,c,** Comparison of gene expression by CAR T cell infiltrates in the control and meso-tumor groups. **$P < 0.0005$, ***$P <$ 1e-6 against meso-tumor and control tumor for MKI67, TCF7, SELL, and LEF1. ***$P <$ 1e-7 against other groups for AGR2 and SERPINA1.

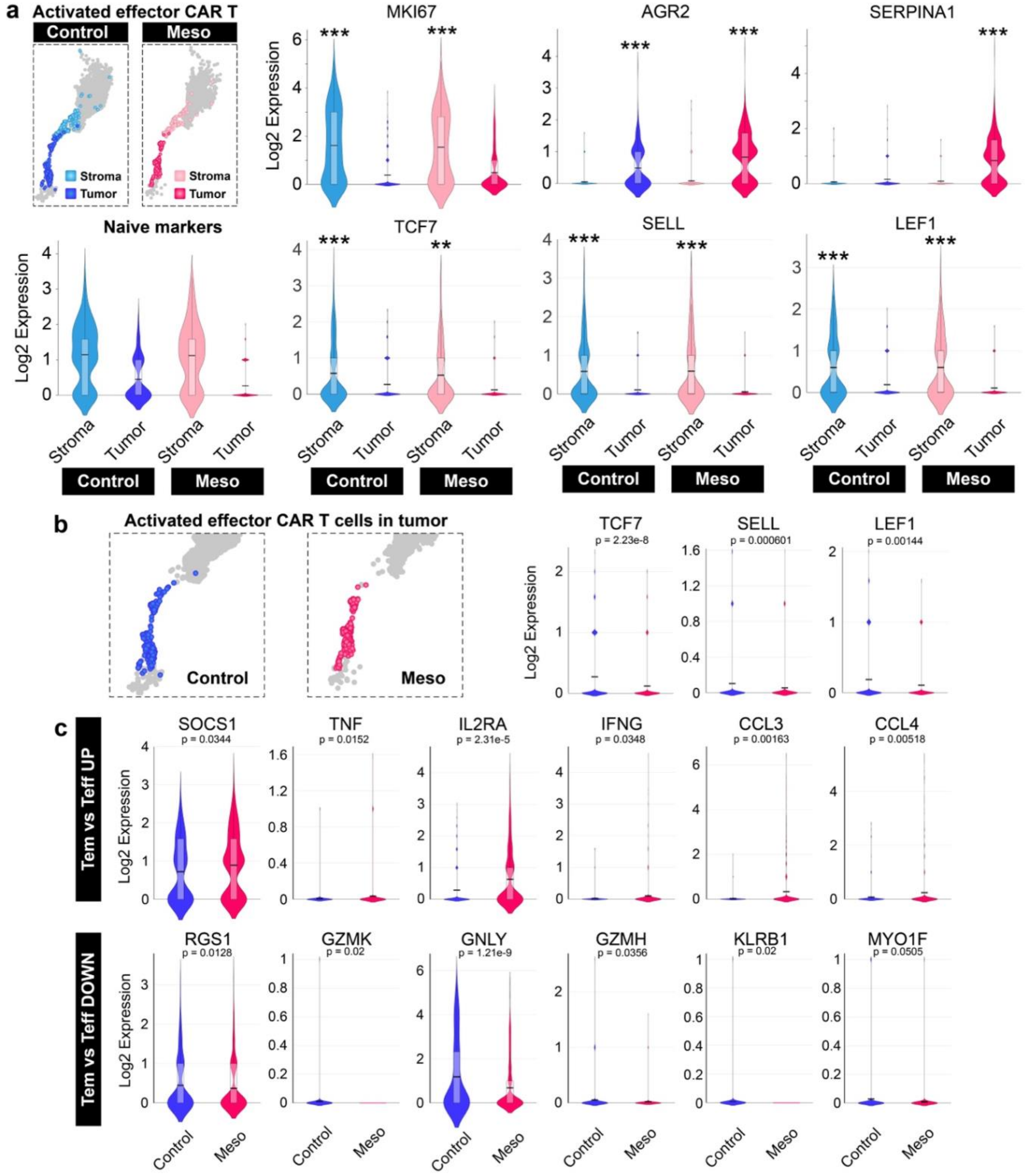

### Supplementary Figure 23. Flow cytometric analysis of CCR2 expression

Quantification of CCR2 expression in non-transduced donor T cells (NTD), meso-CAR T cells (meso-CAR), and meso-CAR T cells with lentiviral transduction of CCR2 (meso-CAR-CCR2).

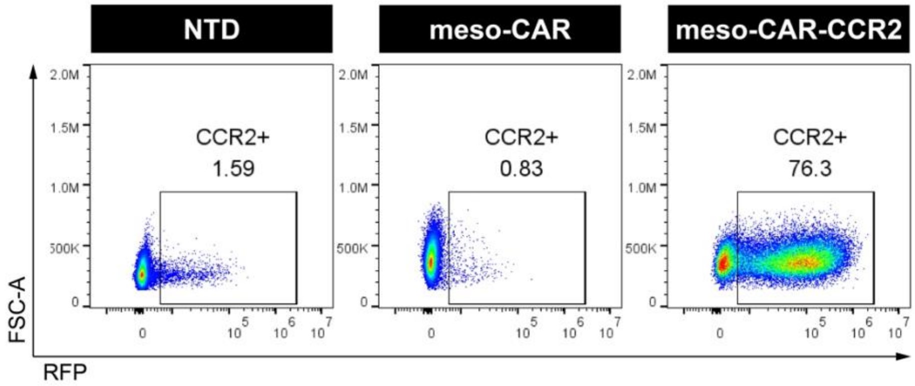

### Supplementary Figure 24. Ligand-receptor interaction analysis

**a,b**, Separate UMAP plots showing the clustering of cells from the control (**a**) and meso-tumor (**b**) groups by cell type (top) and by location of origin (bottom). **c-e**, Chord diagrams and violin plots showing the cell type- and individual cluster-specific expression of interacting gene pairs mediating the crosstalk between tumor cells and activated CAR T cells, including CCL3-IDE (**c**), IL13-TMEM219 (**d**), and TNFSF10-TNFRSF10B (**e**).

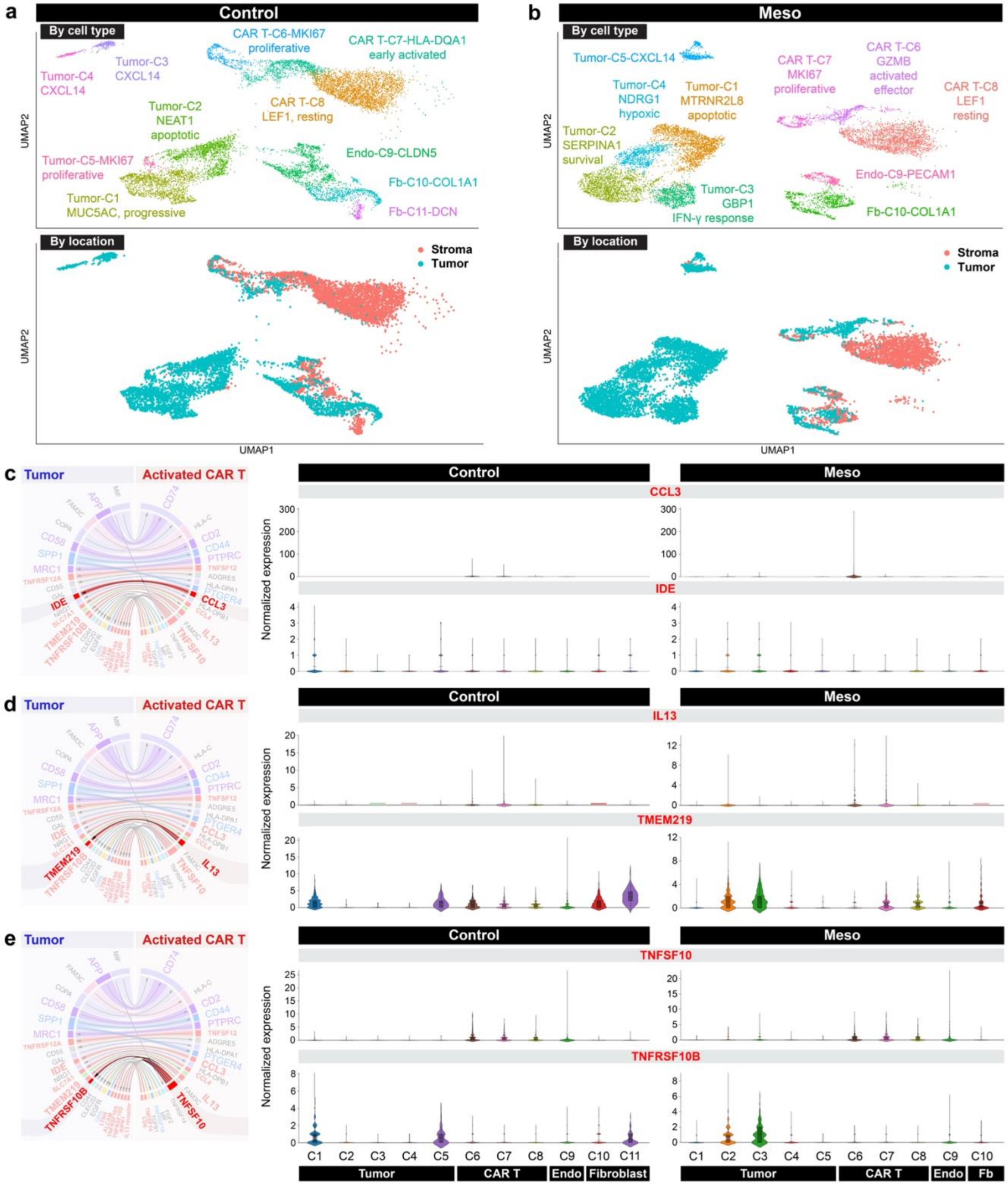

### Supplementary Figure 25. Effects of LAF237 on tumor growth during CAR T infusion

**a,** Representative confocal micrographs of meso-CAR T cell-infused lung tumors treated with different concentrations of LAF237 during long-term culture. Scale bars, 250 µm. **b,** Quantification of CAR T cell trafficking within the region of interest at Day 26. Data are presented as mean ± SEM (n ≥ 4).

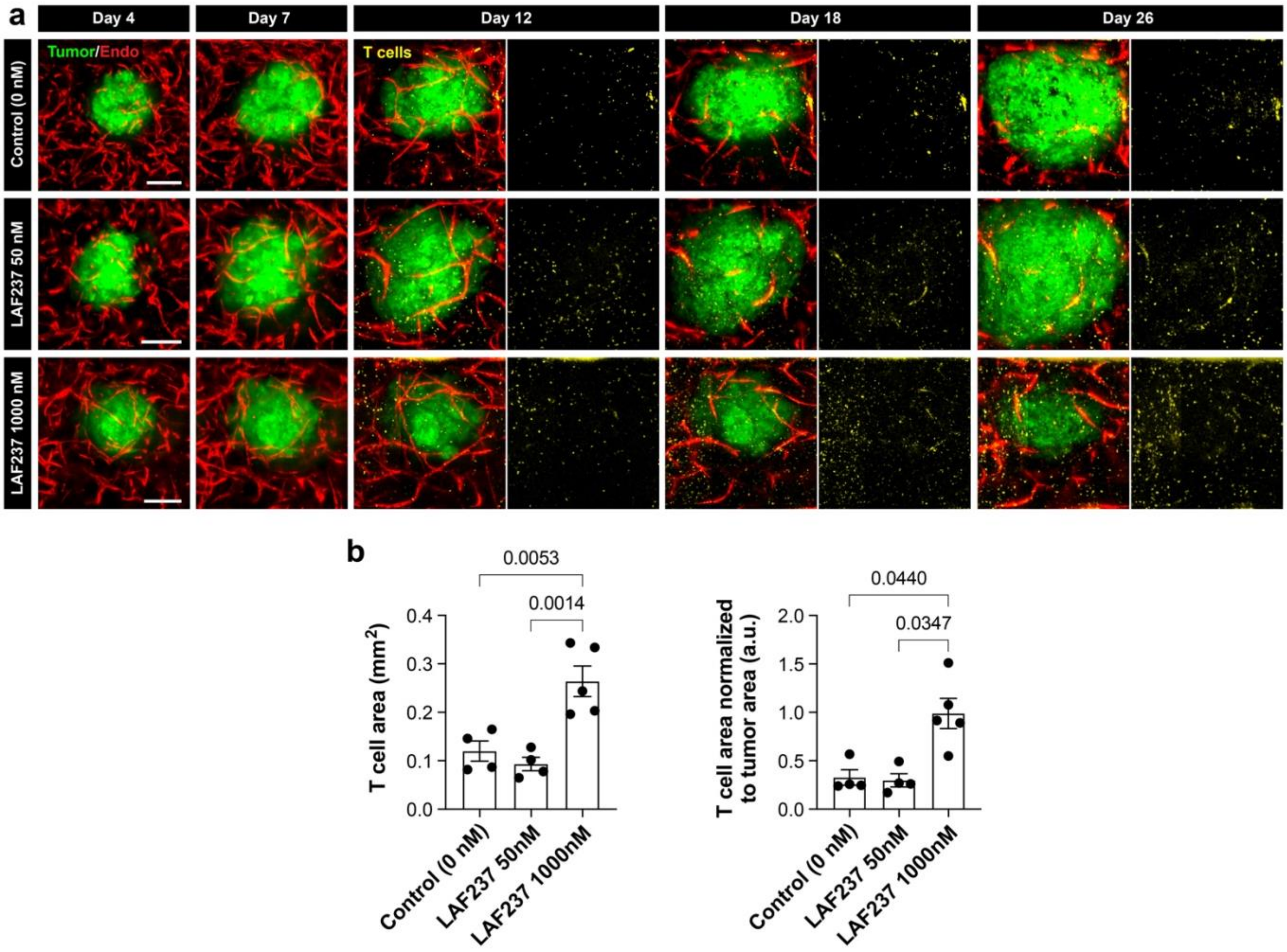

### Supplementary Figure 26. Effects of LAF237 alone on tumor growth

**a**. Representative confocal micrographs of lung tumors during daily treatment with LAF237 in the absence of CAR T cells. **b**. Quantification and comparison of normalized tumor area over time. Data are presented as mean ± SEM (n = 3-6).

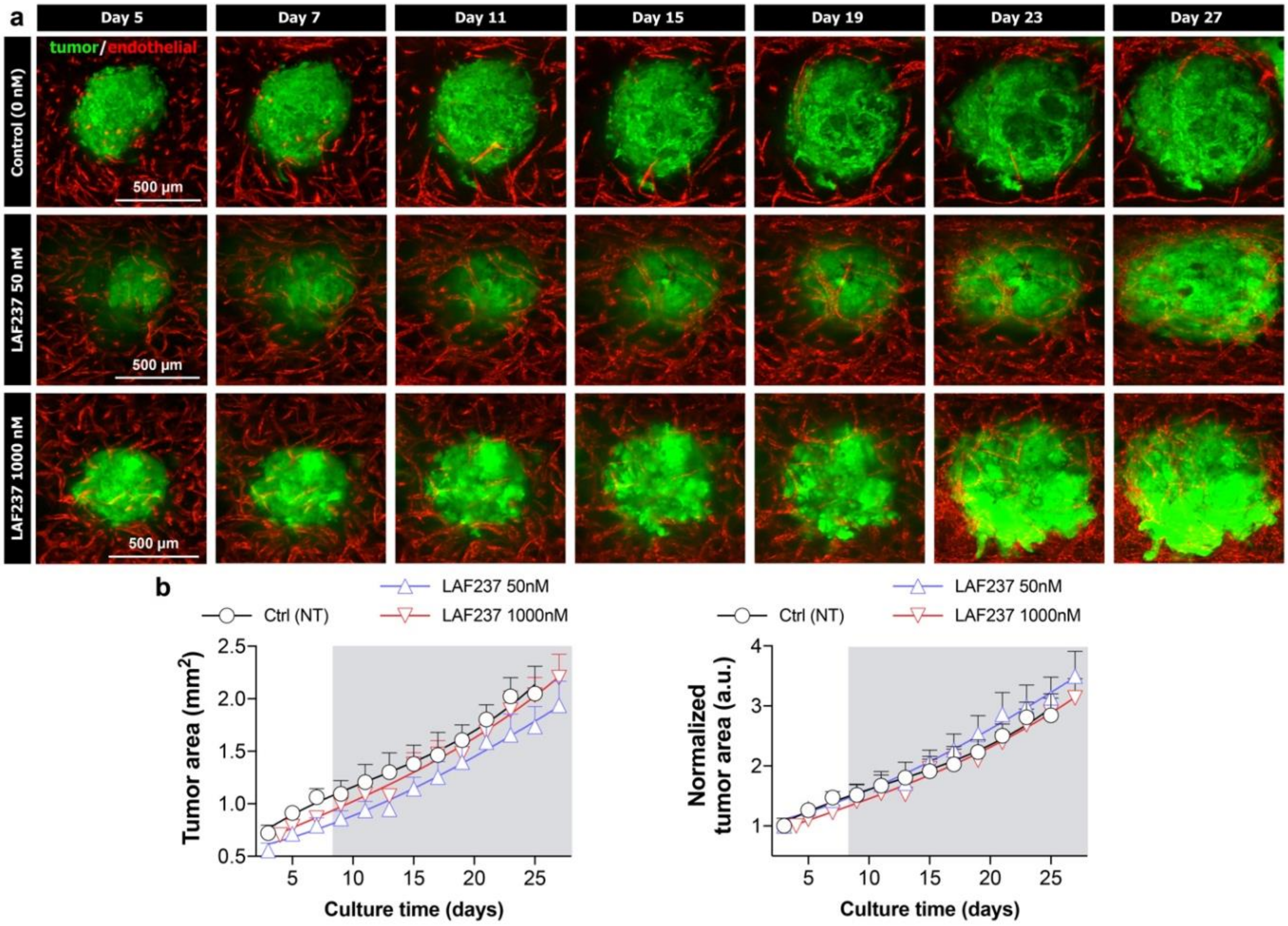

**Supplementary Figure 27. Effect of blocking CXCR3 on the activity of CAR T cells**

**a,** Experimental timeline for meso-CAR T cell infusion and drug treatment with LAF237. **b,** Representative fluorescence micrographs of single meso-tumors infused with meso-CAR T cells without drug treatment (no LAF237), treated with LAF237 at 1000 nM (second row), 1000 nM of LAF237 and anti-CXCR3 antibody (third row), or 1000 nM of LAF237 with isotype-matched control antibody (fourth row). Blood vessels are not shown in these images. Scale bars, 250 µm. **c,d,** Quantification of T cell area (**c**) and normalized tumor area (**d**) over time. Shaded in grey in **d** shows a post-CAR T cell infusion period. Data are presented as mean ± SEM (n ≥ 3). CAR T cells from one healthy donor were tested.

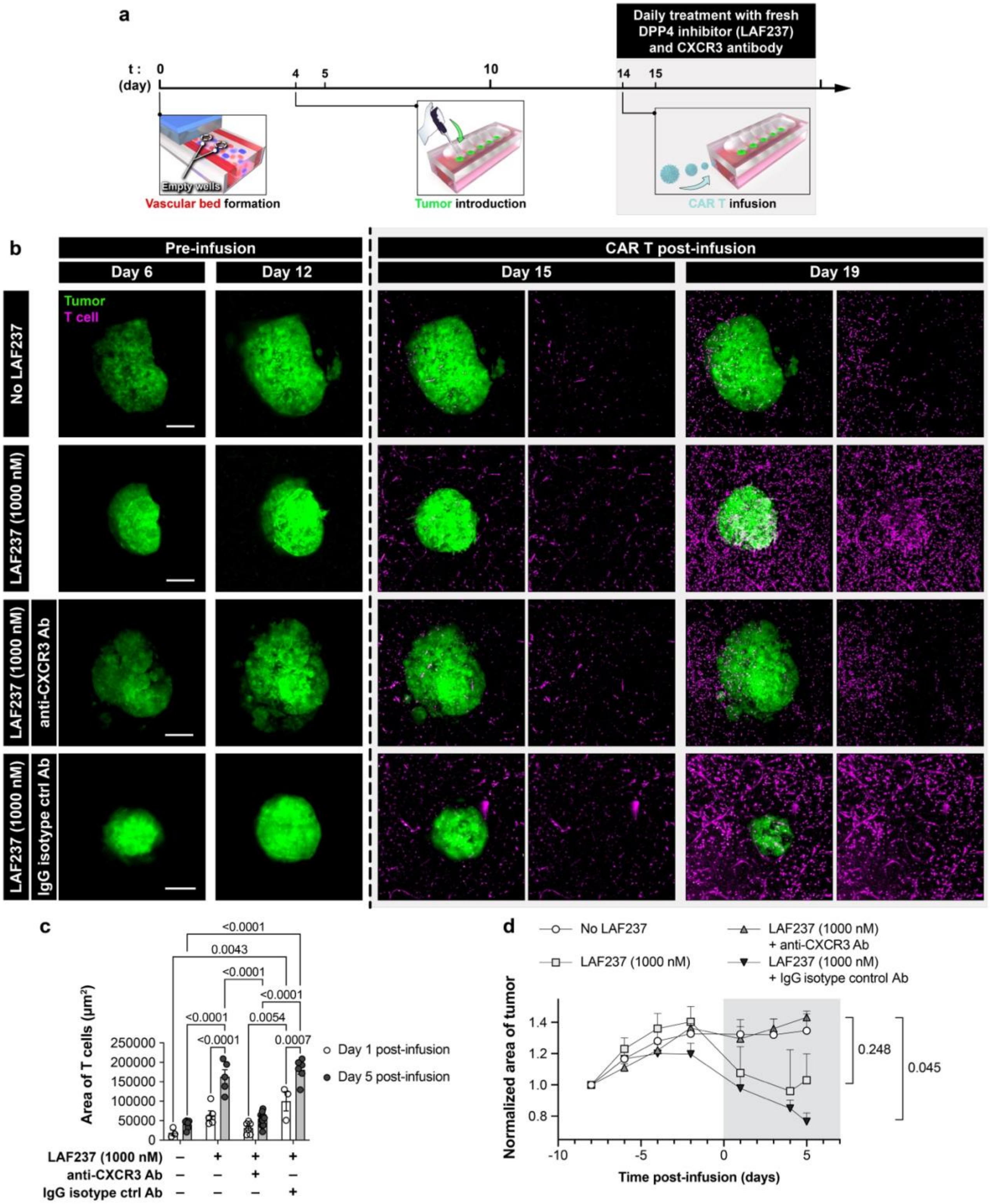

### Supplementary Figure 28. Measurement of cleaved and intact forms of CXCL10

**a**. Design of gold nanoparticles (GNPs) engineered to present valine-specific aptamers on the surface for the detection of valine-proline dipeptides truncated off the intact CXCL10 by DPP4. **b**. UV-vis measurement of the aptamer-modified GNPs after mixing with device effluent samples collected from the experiments described in **Fig. 6** at various time points after CAR T cell infusion. As compared to control (CAR T cell infusion without LAF237 treatment), the high-dose LAF237 group shows smaller peak shifts, indicating reduced truncation of intact CXCL10 due to the inhibition of DPP4. **c**. Raman measurement confirming the presence of valine and proline on the surface of the aptamer-modified GNPs used for the measurement shown in (**b**). **d**. Western blot measurement showing intact CXCL10 protein control (first column), artificially truncated CXCL10 protein control generated by incubation with DPP4 (second column), increased truncation of CXCL10 in the control group without LAF237 treatment from the experiments described in **Fig. 6** at day 8 post-infusion (third column), and preservation of intact CXCL10 in the high-dose LAF237 group from the experiments described in **Fig. 6** at day 8 post-infusion (fourth column). **e**. Unprocessed scan of western blots presented in (**d**).

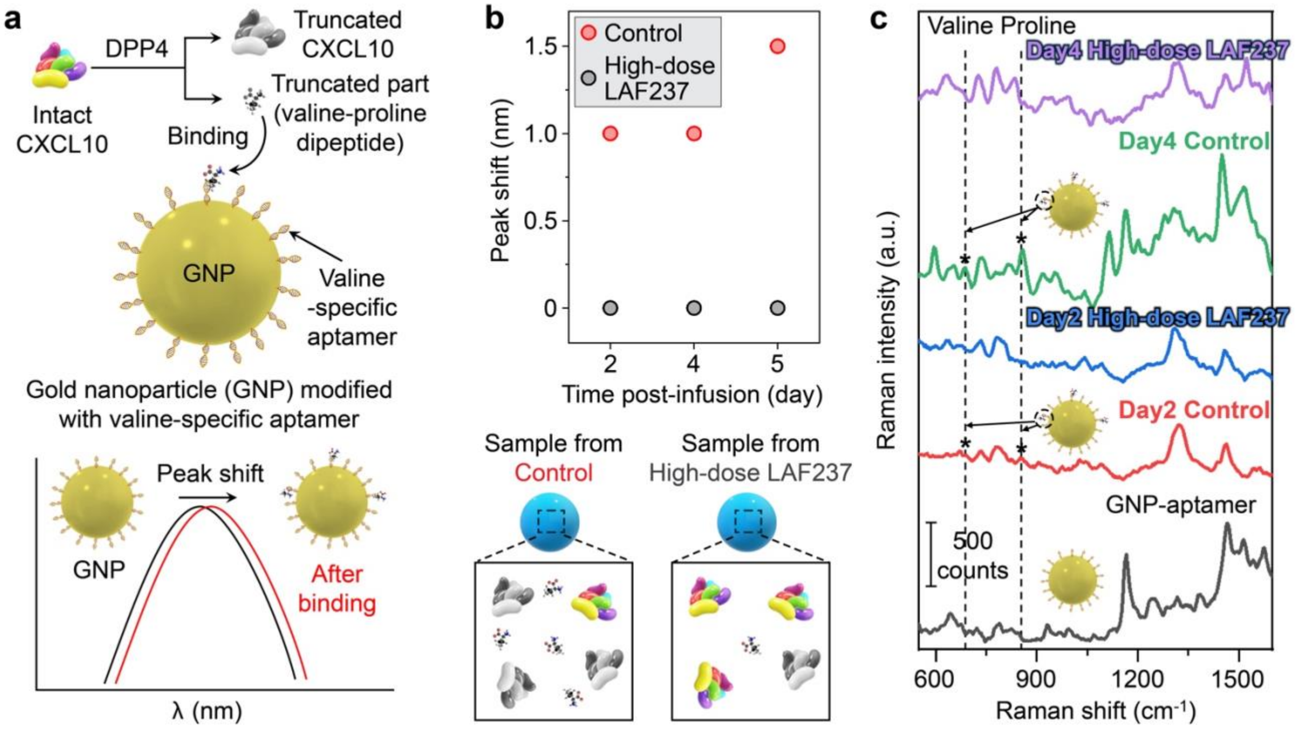


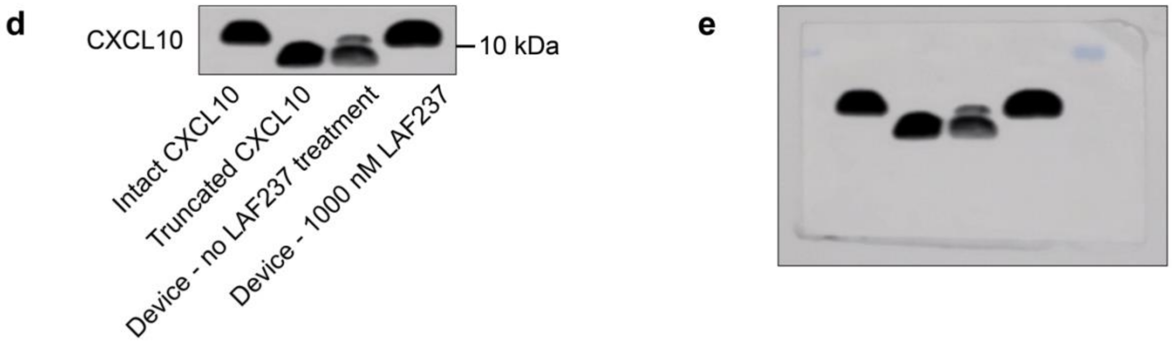

### Supplementary Figure 29. Heatmap overview of significantly changed metabolites

The color in the heatmap indicates the relative abundance of metabolites with red and blue representing higher and lower abundance, respectively.

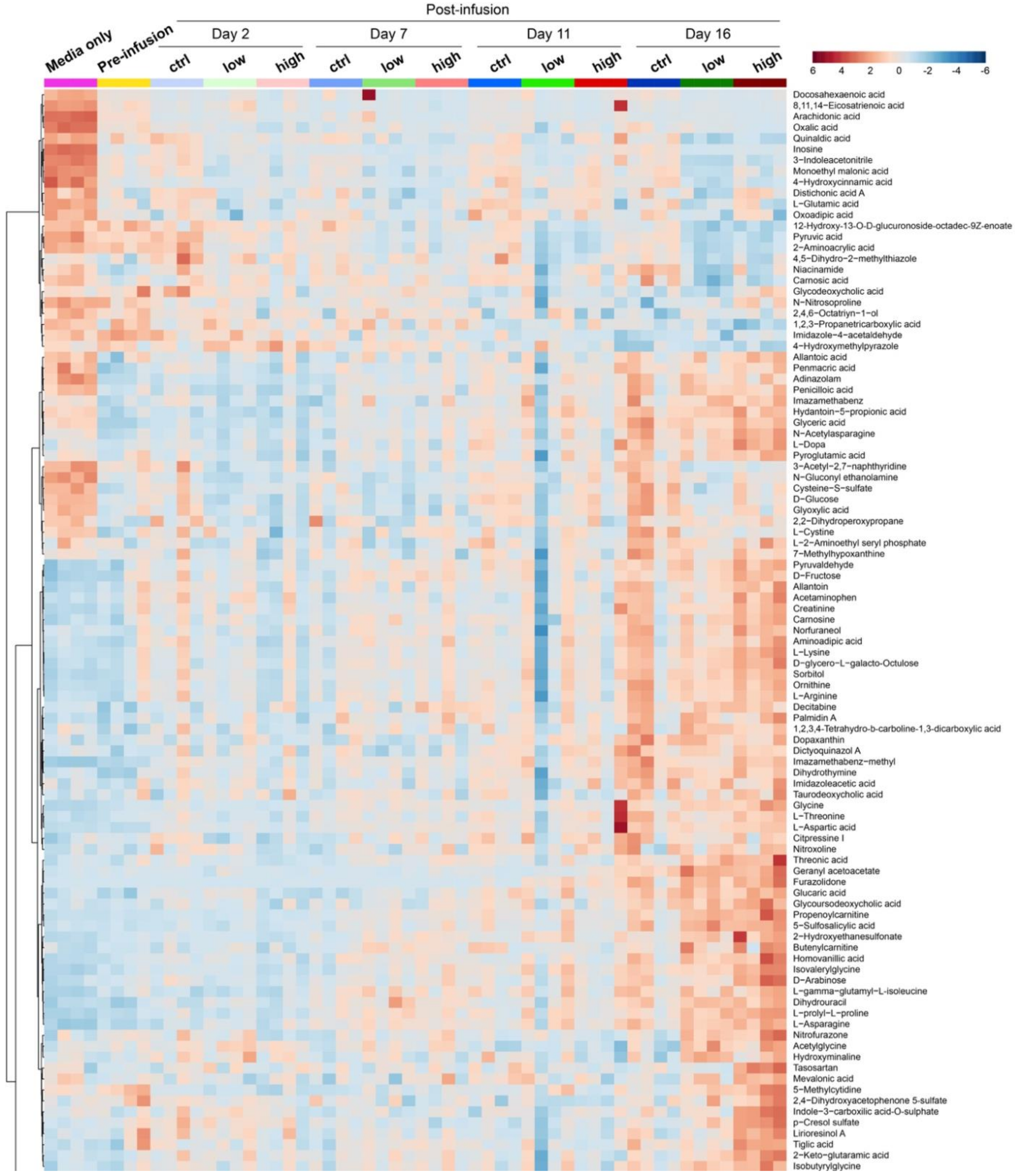

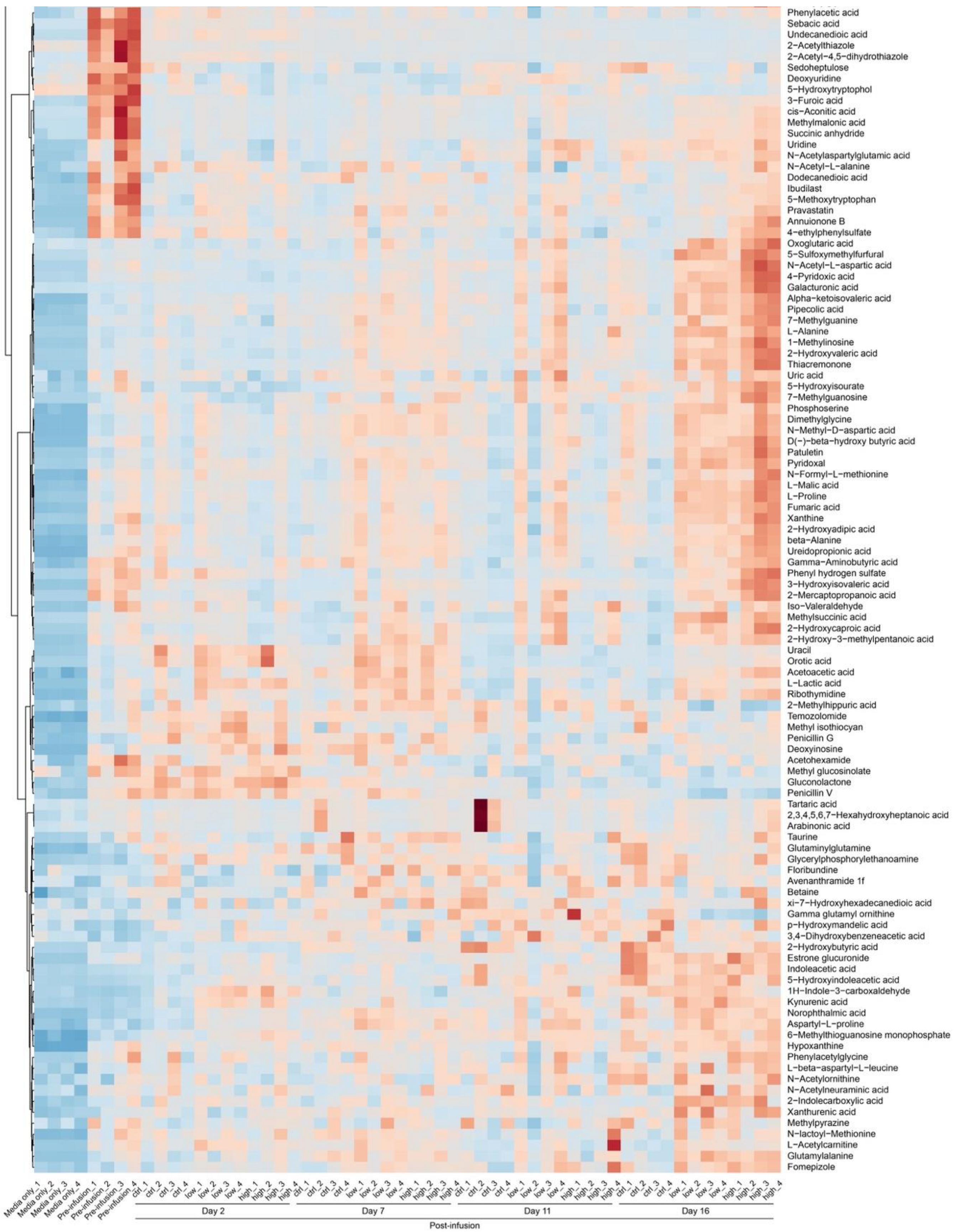
Phenylacetic acid
Sebacic acid
Undecanedioic acid
2−Acetylthiazole
2−Acetyl−4,5−dihydrothiazole
Sedoheptulose
Deoxyuridine
5−Hydroxytryptophol
3−Furoic acid
cis−Aconitic acid
Methylmalonic acid
Succinic anhydride
Uridine
N−Acetylaspartylglutamic acid
N−Acetyl−L−alanine
Dodecanedioic acid
Ibudilast
5−Methoxytryptophan
Pravastatin
Annuionone B
4−ethylphenylsulfate
Oxoglutaric acid
5−Sulfoxymethylfurfural
N−Acetyl−L−aspartic acid
4−Pyridoxic acid
Galacturonic acid
Alpha−ketoisovaleric acid
Pipecolic acid
7−Methylguanine
L−Alanine
1−Methylinosine
2−Hydroxyvaleric acid
Thiacremonone
Uric acid
5−Hydroxyisourate
7−Methylguanosine
Phosphoserine
Dimethylglycine
N−Methyl−D−aspartic acid
D(−)−beta−hydroxy butyric acid
Patuletin
Pyridoxal
N−Formyl−L−methionine
L−Malic acid
L−Proline
Fumaric acid
Xanthine
2−Hydroxyadipic acid
beta−Alanine
Ureidopropionic acid
Gamma−Aminobutyric acid
Phenyl hydrogen sulfate
3−Hydroxyisovaleric acid
2−Mercaptopropanoic acid
Iso−Valeraldehyde
Methylsuccinic acid
2−Hydroxycaproic acid
2−Hydroxy−3−methylpentanoic acid
Uracil
Orotic acid
Acetoacetic acid
L−Lactic acid
Ribothymidine
2−Methylhippuric acid
Temozolomide
Methyl isothiocyan
Penicillin G
Deoxyinosine
Acetohexamide
Methyl glucosinolate
Gluconolactone
Penicillin V
Tartaric acid
2,3,4,5,6,7−Hexahydroxyheptanoic acid
Arabinonic acid
Taurine
Glutaminylglutamine
Glycerylphosphorylethanoamine
Floribundine
Avenanthramide 1f
Betaine
xi−7−Hydroxyhexadecanedioic acid
Gamma glutamyl ornithine
p−Hydroxymandelic acid
3,4−Dihydroxybenzeneacetic acid
2−Hydroxybutyric acid
Estrone glucuronide
Indoleacetic acid
5−Hydroxyindoleacetic acid
1H−Indole−3−carboxaldehyde
Kynurenic acid
Norophthalmic acid
Aspartyl−L−proline
6−Methylthioguanosine monophosphate
Hypoxanthine
Phenylacetylglycine
L−beta−aspartyl−L−leucine
N−Acetylornithine
N−Acetylneuraminic acid
2−Indolecarboxylic acid
Xanthurenic acid
Methylpyrazine
N−lactoyl−Methionine
L−Acetylcarnitine
Glutamylalanine
Fomepizole
Media only_1
Media only_2
Media only_3
Media only_4
Pre−infusion_1
Pre−infusion_2
Pre−infusion_3
Pre−infusion_4
ctrl_1
ctrl_2
ctrl_3
ctrl_4
low_1
low_2
low_3
low_4
high_1
high_2
high_3
high_4
ctrl_1
ctrl_2
ctrl_3
ctrl_4
low_1
low_2
low_3
low_4
high_1
high_2
high_3
high_4
ctrl_1
ctrl_2
ctrl_3
ctrl_4
low_1
low_2
low_3
low_4
high_1
high_2
high_3
high_4
ctrl_1
ctrl_2
ctrl_3
ctrl_4
low_1
low_2
low_3
low_4
high_1
high_2
high_3
high_4
Day 2
Day 7
Day 11
Day 16
Post-infusion

### Supplementary Figure 30. Heatmap of top 50 metabolites

Heatmap of top 50 metabolites identified by the partial least squares discriminant analysis (PLS-DA). The color in the heatmap indicates the relative abundance of metabolites with red and blue representing higher and lower abundance, respectively.

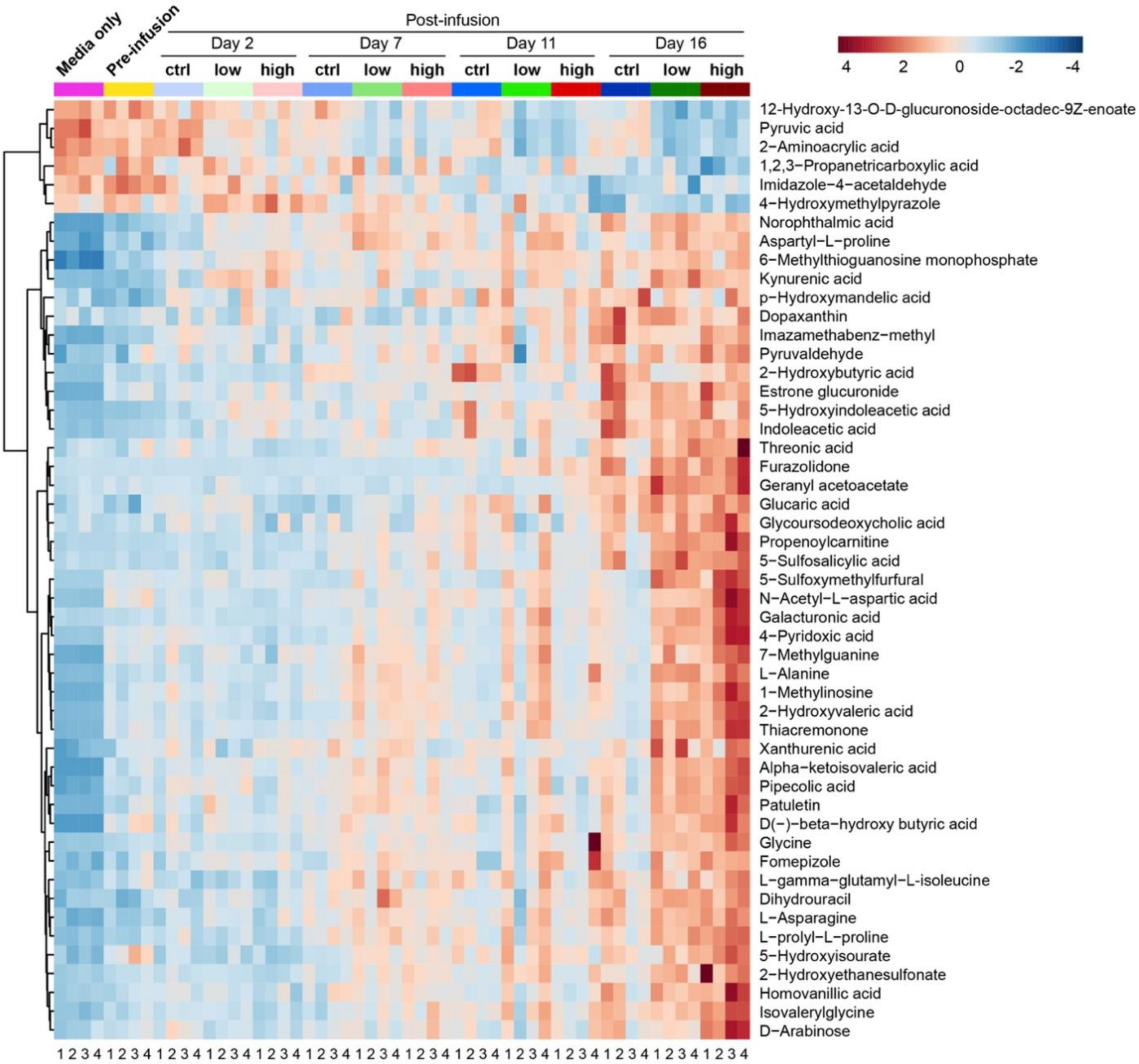

### Supplementary Figure 31. Metabolite sets enrichment analysis

Overview of top 25 significantly enriched metabolite sets from the analysis of 65 significantly upregulated metabolites identified in the high-dose LAF237 group.

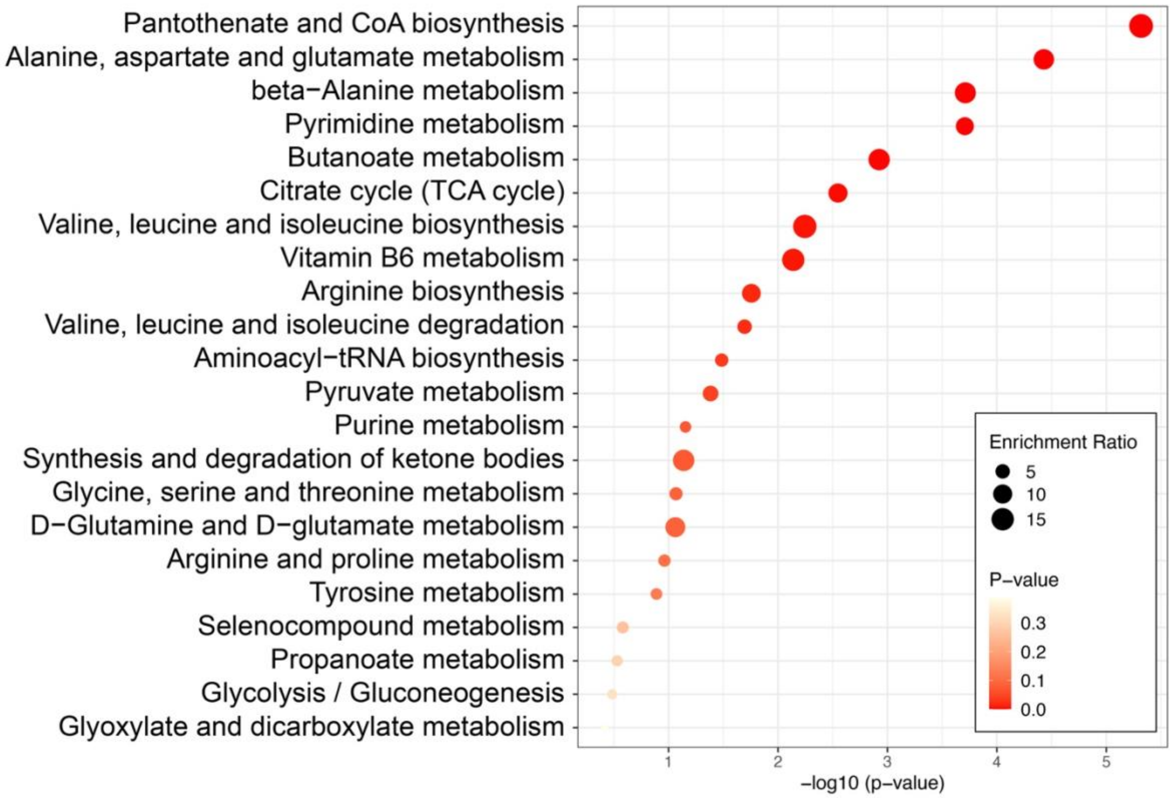

### Supplementary Figure 32. Identification of predictive biomarkers

**a,b,** ROC curves for six biomarker prediction models, with increasing numbers of constituent metabolite features as indicated by the color legend, to identify biomarkers that can differentiate more efficacious CAR T cell treatment with high-dose LAF237 from control CAR T cell treatment without LAF237 at day 16 post-infusion (**a,c,e**) and all time points (**b,d,f**). **c,d,** Predictive accuracy of biomarker models with increasing numbers of features. The most accurate biomarker model is highlighted with a red dot. **e,f**, Predicted class probabilities for all samples using selected biomarker models of (**e**) 25 features and (**f**) 15 features. The classification boundary is at the center (x=0.5, dotted line).

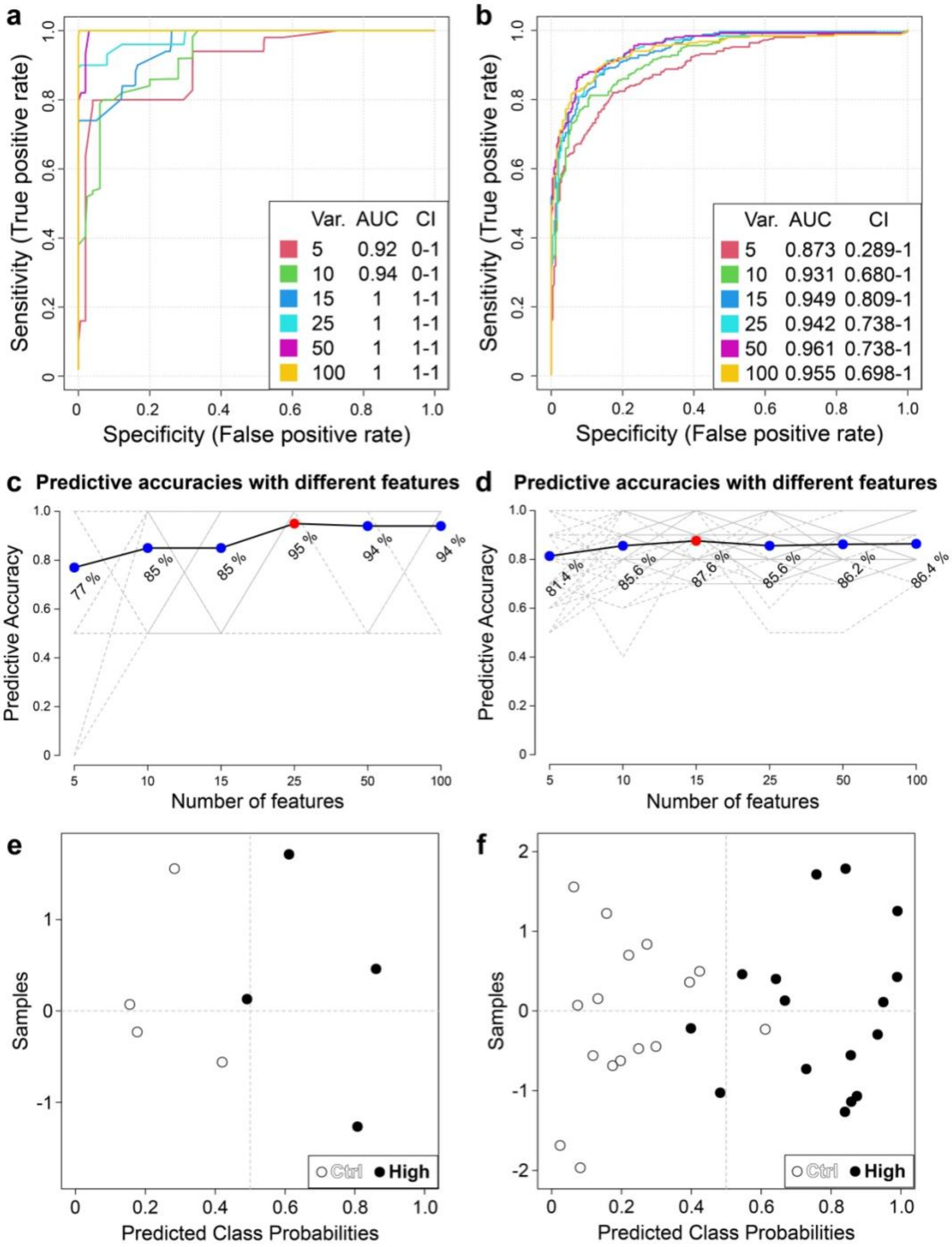

### Supplementary Figure 33. Levels of additional predictive biomarkers

Boxplots show minimum, 25th percentile, mean, 75th percentile, and maximum. N = 4 for each group. Note that post-hoc pairwise comparison is only calculated between the high-dose and control groups at all time points with *p < 0.05.

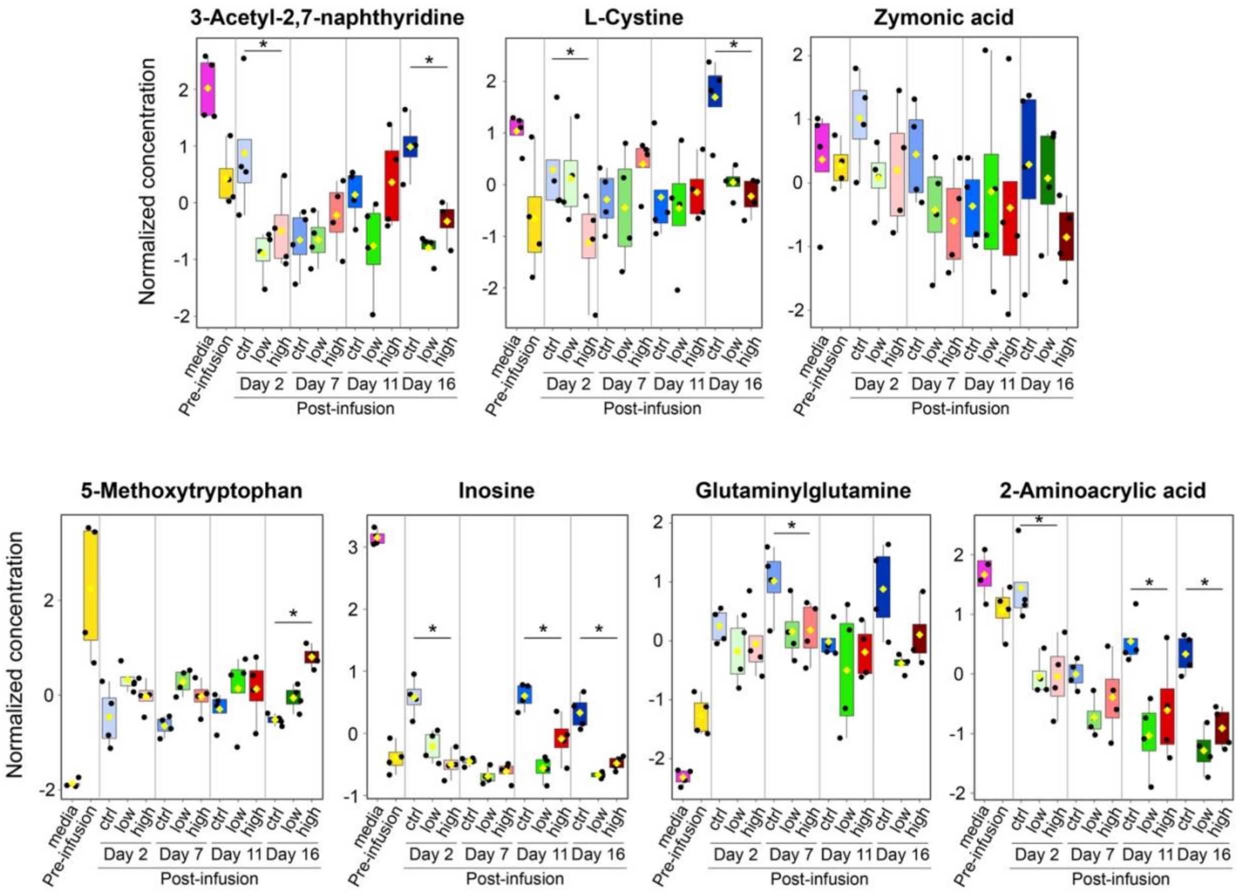

### Supplementary Table 1. List of significant interactions in CellPhoneDB analysis

This table provides the list of significant ligand-receptor interaction pairs from each permutation of individual cluster pair in the control and meso-tumor groups as shown in **Supplementary Fig. 24**. The values in the table represent the mean expression of only significant interactions that have $p < 0.05$. Abbreviations: partner_a, CellPhoneDB ID for the first interaction partner protein; partner_b, CellPhoneDB ID for the second interaction partner protein; source, reference from CellPhoneDB; secreted, True if one of the partners is secreted and False if both are membrane bound; is_integrin, True if one of the partners is integrin.

### Supplementary Table 2. Peak intensities of metabolites

This table contains the matrix of peak intensities of identified and annotated metabolites from all samples in conditions including fresh media only, pre-infusion, CAR T infusion alone (ctrl), CAR T infusion with 50 nM LAF237 (low), CAR T infusion with 1000 nM LAF237 (high) at days 2, 7, 11, and 16 post-infusion.

### Supplementary Table 3. Summary of MSEA and pathway impact analysis

This table shows a summary of metabolite sets enrichment analysis (MSEA) and pathway impact analysis (in separate sheets) of the upregulated metabolites identified in the high-dose LAF237 group, as shown in **Fig. 7c** and **Supplementary Fig. 31**.
